# STRANDS OF CONNECTION: LIVESTOCK GRAZING, VEGETATION DENSITY, AND ORB-WEAVER SPIDER PERSISTENCE

**DOI:** 10.1101/2023.11.19.567706

**Authors:** Guilherme Oyarzabal, Murilo Guimarães

## Abstract

Studies on the effects of grazing disturbances in grasslands have shown mixed results for spider diversity, mainly regarding their guilds. While ungrazing, low, and moderate grazing potentially enhance diversity of orb-weavers, heavy grazing seems to reduce specieś richness. On the population level, studies of orb-weavers are scarce, and the effects of grazing are unknow. In this way, we investigated the effects of different levels of grazing on population persistence of orb-weaver spiders, hypothesizing that low to intermediate disturbances benefit populations. We predict that high grazing, due to the removal of vegetation structure, will negatively affect occupancy and abundance of orb-weavers. For that, we experimentally controlled grazing pressure and obtained population occurrence and counts of two orb-weaver spider species, *Argiope argentata* and *Alpaida quadrilorata*. We found that *A. argentata* was directly affected by grazing, as it relies on higher vegetation for web-building. In contrast, *A. quadrilorata*, which occurs in cattle-resistant rosette plants, showed no effects of grazing. **Implications for insect conservation:** Our study emphasizes the need for balanced grazing practices and habitat conservation to protect orb-weaver spiders and other arthropods, as well as, species-specific effects for species from the same guild, underscoring their ecological significance in maintaining ecosystem stability.

## Introduction

Generalist predators are midranking carnivorous that vary in many shapes and sizes, playing an important role in regulating, mainly, herbivores (Gagnon et al. 2019; Macé et al. 2019; Michalko et al. 2019; Wray et al. 2021). Such predators usually occur sympatrically and have similar needs of food and energy intakes, as well as broader diets than top predators (Lesmeister et al. 2015; Whitney et al. 2018; Wray et al. 2021), providing them superior resilience to disturbances (Wimp et al. 2019). However, generalist predators are largely affected by chronic anthropogenic disturbances (Rito et al. 2017; Antongiovanni et al. 2020) such as agriculture and livestock, since these activities may threat them through human conflict, environment degradation and declines in prey population (Wang et al. 2015; Newsome et al. 2017; Wimp et al. 2019).

Among arthropod generalist predators, spiders have been one of the main focus in studies of grazing chronic disturbance (Macé et al. 2019; Filazzola et al. 2020). Despite some controversy and inconclusive results (Silva and Ott 2017; Muvengwi et al. 2018; Samu et al. 2018), authors usually suggest that ungrazing, low and moderate grazing may enhance spider diversity while heavy grazing reduce species richness and abundance, as well as species turnover (Szmatona-Túri et al. 2017; Wang et al. 2019; Ferreira et al. 2020; Oyarzabal and Guimarães 2021). Moreover, grazing disturbance seems to affect spider guilds in opposite ways, where diversity of ground dweller spiders may be enhanced while orb-weaver spiders appear to be particularly sensitive to vegetation removal (Nogueira and Pinto-da-Rocha 2016; Neilly et al. 2020), losing species richness in high grazing environments (Oyarzabal and Guimarães 2021).

The removal of above-ground plant biomass provoked by chronic grazing (Tälle et al. 2016; Pett and Bailey 2019; Ferreira et al. 2020; da Silva Bomfim et al. 2021) directly affects the primary hunting strategy of orb-weavers, the ability to build webs using the tridimensional vegetal structure (Nogueira and Pinto-da-Rocha 2016). Without physical structures to build their web, species are unable to find prey (Torma et al. 2019; Helden et al. 2020; Fischer et al. 2021) and mates (Cory and Schneider 2018; Weiss and Schneider 2021), as well as avoid predation (Blackledge and Wenzel 1999; da Silva Bomfim et al. 2021; Narimanov et al. 2021). Consequently, the simplification of habitat structure induced by grazing can culminate in the decline of local populations, resulting in the exclusion of orb-weaver species from grasslands (Oyarzabal and Guimarães 2021). However, the emphasis has remained on the community aspects of orb-weavers predation (Rodrigues et al. 2009; da Silva Bomfim et al. 2021), leaving a notable gap in understanding how distinct species, and even genera, respond to the persistent disturbances induced by grazing.

In this way, our objective was to assess the effects of grazing pressure on populations of two abundant grassland orb-weaver spiders (Rodrigues et al. 2009; Nogueira and Pinto-da-Rocha 2016), *Argiope argentata* (Fabricius, 1775) and *Alpaida quadrilorata* (Simon, 1897) (Fig. 1). We hypothesize that the population of both spider species are directly affected by different levels of grazing disturbance. We predict that low and intermediate grazing will benefit both species but they will respond differently from each other. Hence their habitat use and abundance will be negatively affected, mostly by heavy grazing, due to the removal of vegetation.

**Figure 1.**
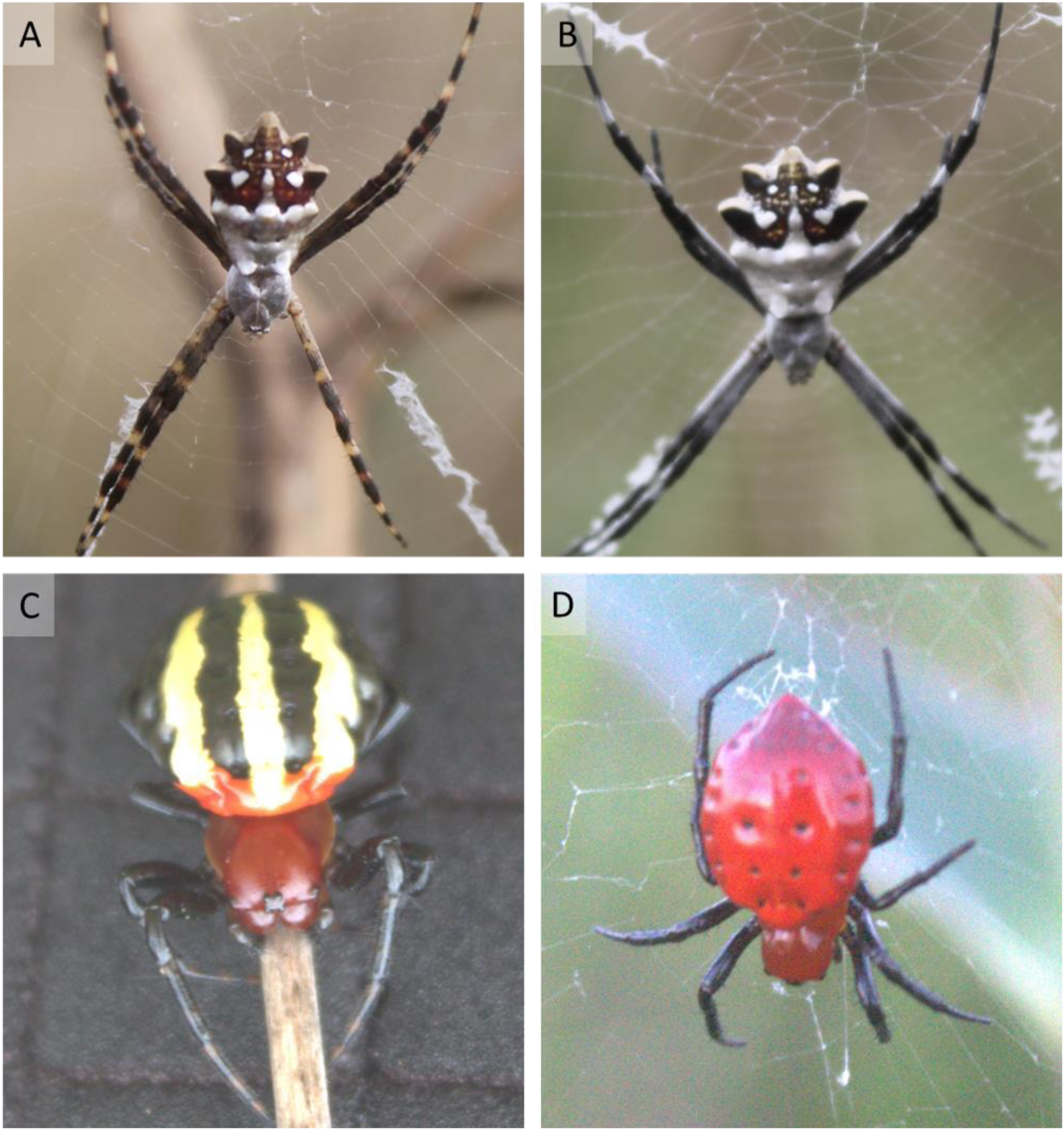
Orb-weaver spider species from the Araneidae family. Photos A and B represent *Argiope argentata* and photos C and D represent *Alpaida quadrilorata*.

## Methods

### Species studied

The first species, *A. argentata*, have a broad range distribution, from Canada to Argentina (Agnarsson et al. 2016; World Spider Catalog 2023) and inhabits low above ground plants on grassland and margins of roads and trails (Robinson 1969). The second species, *A. quadrilorata*, is distributed in Argentina, Brazil, Paraguay and Uruguay (Vasconcellos-Neto et al. 2017; World Spider Catalog 2023) and it is known to inhabit, almost exclusively, plants with rosette-shaped leaves that has similar architecture to bromeliads Uruguay (Levi 1988; Vasconcellos-Neto et al. 2017; Hesselberg et al. 2023).

### Study site and sampling design

Our study took place in the Pampa grasslands, southern South America. The climate is subtropical with hot-dry summers and humid-cold winters. Temperatures surpass 40°C in summer and vary from 4°C to 28°C in winter. Rainfall ranges between 1,200 and 1,600 mm through the year (Kottek et al. 2006). Sampling occurred at Estação Experimental Agronômica da Universidade Federal do Rio Grande do Sul (UFRGS) located in Eldorado do Sul municipality, Rio Grande do Sul, Brazil (30°06’08’’S; 51°40’56’’W). Since 1987, an experiment called *Nativão* is conducted to assess the effects of different intensities of cattle grazing in an area that covers about 52 hectares (Nabinger et al. 2009). In the year 2000, the area was subdivided in 14 plots with different cattle grazing treatments that vary in fixed and daily levels of grass forage supply for cattle, expressed in kg of vegetal dry matter [DM]/100 kg of live weight [LW] (% LW). In this way, these areas are defined by a percentage of vegetal dry matter remaining, meaning the less vegetal dry matter that remains, the greater the grazing pressure (Nabinger et al. 2009).

Six plots were selected for sampling: two plots (3.05 ha and 3.14 ha) with high grazing disturbance (4% LW, around 0.86 Animal Units (AU)/ha/year); two plots (2.73 ha and 3.67 ha) of moderate disturbance (8% LW, around 0.59 AU/ha/year); and two plots (5.27 ha and 5.42 ha) of low grazing disturbance (16% LW, around 0.45 AU/ha/year) (Nabinger et al. 2009). Considering the known home range and movement capacity of one of our target species (Craig et al. 2001), we superimposed a grid on the top of each of the six plots with cell size 5x5m, using the QGIS software (QGIS.org 2020). From the total of cells per plot, a subgroup of 50 cells was randomly sorted for all surveys (50 cells per plot, 300 in total). Then, on each campaign 16 randomly selected cells were surveyed from the 50 pre-selected cells of each plot (96 in total per campaign). Lastly, the order that the plots were surveyed was always randomized in each campaign.

Seven-monthly campaigns were conducted in year 1, from October 2017 to April 2018, and six-monthly campaigns in year 2, from October 2018 to April 2019, during austral spring and summer, when spiders are more active (Nei et al. 2015). Each campaign was composed by two days (surveys) and species were surveyed in the field from dawn to mid-day (06:00 am to 12:30 pm) and from afternoon to dusk (03:00 pm to 09:00 pm). Cells were surveyed until exhaustion, counting adults and juveniles of *A. argentata* and *A. quadrilorata* species. Two to three trained observers were deployed on each campaign (a total of eight people through the experiment).

### Data analysis

Environmental variables were registered through the campaigns and surveys to be used as occupancy, abundance, and detection predictor variables. To estimate occupancy probability and abundance, we considered vegetation density and the quadratic effect of vegetation density as spatial variables. Although vegetation density is directly correlated with the different grazing treatments in our field site, during our field work we detected variation on vegetation height within the same plots. Therefore, vegetation density was obtained taking four photos of the vegetation on each cell and year, using a 1x1-m white cardboard as a background to measure vegetation density on every photo. Then, we used ImageJ software (Schneider et al. 2012) to convert images to black and white scale, hence, the black pixels were counted as a measure of vegetation density in contrast with the white cardboard background (Ford et al. 2017). The arithmetic mean of black pixels between the four photos was considered as a proxy of vegetation density for each cell and in each year. To estimate detection probability, we included air temperature (degrees Celsius) and time (expressed as minutes after midnight) as temporal predictors. Air temperature was measured three times during each survey (beginning, middle and end). Time was taken on the beginning of each cell survey. Moreover, detection probability was also estimated as a function of vegetation density, and the quadratic effect of vegetation density. All numeric variables (temperature, time, and vegetation density) were standardized to have zero mean and one standard deviation.

We fitted single-season models independently for each sampled year and species. We estimated occupancy (Ψ) and detection (p) probabilities using occupancy modeling (MacKenzie et al. 2002). To estimate abundance (N) we fitted N-Mixture models (Royle 2004) using counts of spiders per cell as our response. We used Akaike’s Information Criterion (AIC) to compare and rank occupancy and N-Mixture models considering models with Delta AIC ≤ 2 those best supported (Arnold 2010) (Supplementary Material 1). Models were model-averaged to provide parameters estimates. Models were built using ‘unmarked’ package (Chandler et al. 2021), and model-averaged using ‘MuMIn’ package (Bartoń 2019), both in the software R (R Core Team 2022). The detection probability estimates and the effects of temporal variables are not discussed but are presented in the supplementary material (Supplementary Material 1).

## Results

A total of 889 individuals of *A. argentata* (25 in high, 278 in moderate and 586 in low grazing) were found in 324 cells in year one and 883 individuals (83 in high, 510 in moderate and 290 in low grazing) were found in 289 cells in year two. For *A. quadrilorata*, a total of 430 individuals (one in high, 266 in moderate and 163 in low grazing) were found in 198 cells in year one and 348 individuals (three in high, 263 in moderate and 82 in low grazing) were found in 178 cells in year two.

### Argiope argentata occupancy and abundance estimates

In year one, *A. argentata* occupancy estimates strongly increased with vegetation density but not significantly (occupancy β_VegY1_ = 40.404, p = 0.087) while its abundance estimates decreased with the quadratic effect of vegetation density (abundance β_Veg_^2^_Y1_ = -0.147, p = 0.050) (Fig. 2). In year two, occupancy estimates increased with vegetation density (occupancy β_VegY2_ = 2.953, p = 0.017) while abundance estimates decreased with the quadratic effect of vegetation density (abundance β_Veg_^2^_Y2_ = - 0.343, p = 7.51e-4).

**Figure 2.**
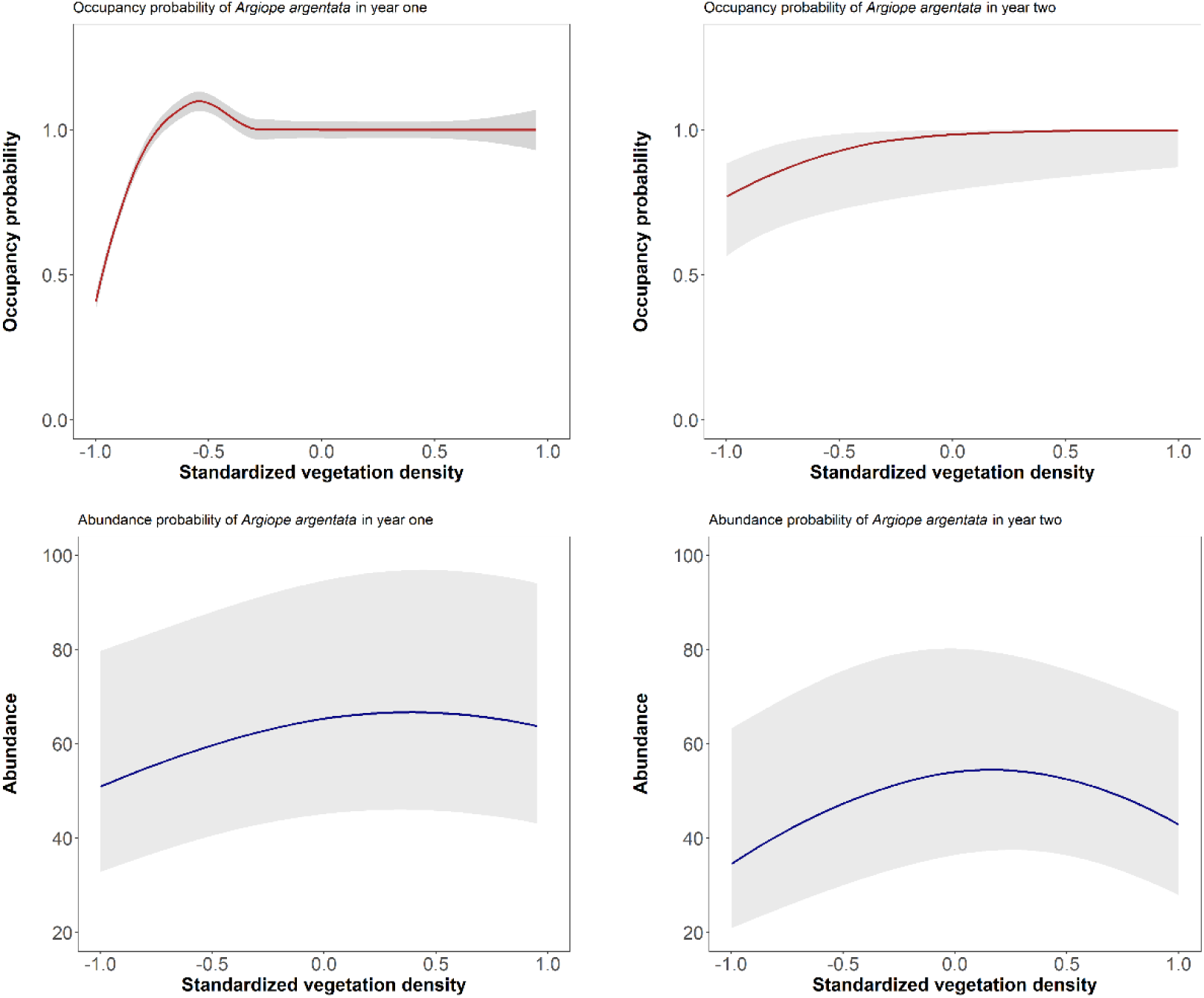
Occupancy (upper) and abundance (lower) estimates for *Argiope argentata* in year one (right) and year two (left). Red lines indicate mean occupancy estimates for year one and two. Blue lines indicate mean abundance estimates for year one and two. Grey shadows indicate standard deviations.

### Alpaida quadrilorata occupancy and abundance estimates

In year one, neither occupancy nor abundance estimates of *A. quadrilorata* were affected by vegetation density (occupancy β_VegY1_ = -0.192, p = 0.342; and abundance β_VegY1_ = -0.008, p = 0.946). In year two, we found the same trend, neither occupancy nor abundance estimates of *A. quadrilorata* were affected by vegetation density (occupancy β_VegY2_ = 1.714, p = 0.059; and abundance β ^2^ = 0.132, p = 0.814) (Fig. 3).

**Figure 3.**
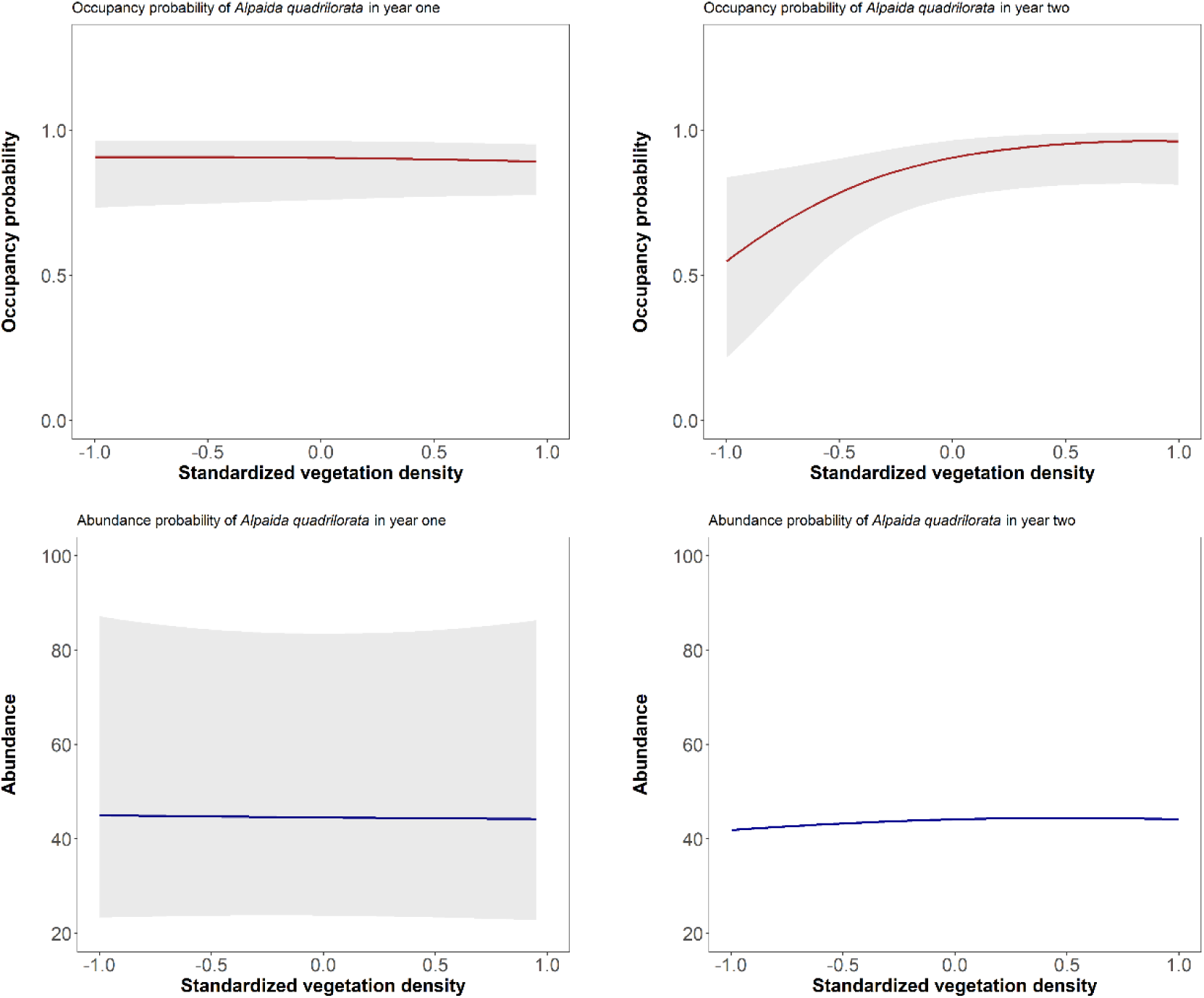
Occupancy and abundance estimates of *Alpaida quadrilorata* in year one and year two of sampling. Red lines indicate occupancy estimates for year one and two. Blue lines indicate abundance estimates for year one and two. Grey shadows indicate standard deviations.

## Discussion

Based on the results, our findings partially align with our hypothesis that both spider populations are influenced by grazing disturbances, consistent with the broad trend observed in the orb-weaver community. Only *A. argentata*, responded to vegetation density, corroborating past findings where orb-weaver spiders seem to benefit from intermediate levels of grazing (Hu et al. 2019; Wang and Tang 2019; Ferreira et al. 2020; Oyarzabal and Guimarães 2021). Interestingly though, our results support the controversy of grazing impact being positive, negative or neutral on spider species, both at the community and population levels (Szmatona-Túri et al. 2017; Samu et al. 2018; Hu et al. 2019; Wang and Tang 2019; Ferreira et al. 2020; da Silva Bomfim et al. 2021; Oyarzabal and Guimarães 2021).

Considering our results, both orb-weaver spiders are responding differently to grazing pressure, likely due to their specific microhabitat requirements. The species *A. argentata* can be easily found on the grass, using the leaves, stems and patches to construct their webs (Robinson 1969; Ayoub et al. 2023; Hesselberg et al. 2023), which may reach up to 100 centimeters above ground (Oyarzabal, personal observation). Consequently, the impact of grazing on *A. argentata* would be direct. As the cattle consume or trample the grass, they inadvertently eliminate potential anchoring points for webs, occasionally even consuming the spiders themselves (Ben-Ari and Inbar 2013; Gish et al. 2017). For *A. quadrilorata* though, we found the species exclusively inhabiting *Eryngium horridum* Malme (referred to as Gravatá or Caraguatá), a plant characterized by rosette-shaped leaves and thorns. This plant is not consumed by cattle and is also resistant to its trampling (Balph and Malecheck 1985; Fidelis et al. 2009; Kurtz et al. 2018; Boavista et al. 2019). Consequently, given this intricate ecological association between the spider, its host plant, and cattle, the grazing effect on *A. quadrilorata* would be indirect.

In face of disturbance, orb-weavers and other spiders are known to disperse through ballooning (Eberhard 1987; Sheldon et al. 2017; Piacentini et al. 2021). Therefore, dispersal would be a good alternative for both species, since the constant grazing could increase the energy cost of rebuilding a web while it lowers their feeding capacity (Prestwich 1977; Janetos 1982; Tanaka 1989; Uetz 1992; Fischer et al. 2019). However, concrete evidence for ballooning behavior exists solely for *A. argentata* (Agnarsson et al. 2016), whereas support for *A. quadrilorata* ballooning is restricted to its genus, *Alpaida* (Eberhard 1987). The challenge though would not be the dispersion, but rather finding plant structures for *A. argentata* and a host plant for *A. quadrilorata*. In this case, the species would need some chemical, physiological or mechanical mechanism to identify the plants. Spiders have a complex chemical communication system that involves pheromones for mate and offspring recognition (Guimarães et al. 2018; Fischer et al. 2019; Beyer et al. 2021). However, few authors suggest the attractiveness of plant chemicals for spiders (Fischer et al. 2018, 2019, 2021). Moreover, these species have a poor vision compared to other spiders like Jumping spiders (Salticidae) (Pollard et al. 1987; Richman and Jackson 1992). Hence, they would only perceive light or shade incidence in the environment, which we know that is important for hunting strategies (Herberstein et al. 2000; Blamires et al. 2007; da Silva et al. 2021). Even so, the mechanisms of how orb-weavers spider perceive a suitable habitat to build their webs it is still a mystery. Hence, delving into light incidence and shade coverage emerges as promising avenues to investigate habitat choice in orb-weaver spiders.

As advocated by other authors (Meadows et al. 2017; Barton et al. 2020) and partially supported by our results, the use of arthropod generalists seems promising as a proxy to study management, chronic anthropogenic disturbances, and conservation. Besides the study of grazing disturbance, arthropods in general may be amazing candidates to study fine scale climate change (Staude et al. 2018; Høye 2020). Variations in temperature and rainfall have been affecting arthropod survival, reproduction, body size, clutch size, behavior, and physiology (Supriya et al. 2019; Walsh et al. 2019; Høye 2020), as observed in chordates such as anurans (González-del-Pliego et al. 2020), reptiles (Diele-Viegas et al. 2020), birds (Bateman et al. 2020) and mammals (Mitchell et al. 2018). The conservation and protection of charismatic animal species (e.g., mammals), who may function as umbrella species for the environment (Schlagloth et al. 2018; Wang et al. 2021), is undoubtfully important. However, given the importance of arthropods in food webs, both as predators and prey, and in the ecosystem functioning, their disappearance induced by anthropogenic actions may lead to unpredictable ecosystem dynamics, undoubtedly cascading onto ecosystem services and hence jeopardizing landscape conservation (Klaus et al. 2013; Blubaugh et al. 2017; Goulson 2019; Samways et al. 2020).

In conclusion, our study provides valuable insights into the complex interplay between cattle grazing and its impact on orb-weaver spider populations, specifically *A. argentata* and *A. quadrilorata*. While our findings partially align with our initial hypothesis that grazing disturbances influence spider populations, they also reveal the intricate nature of these effects. While *A. argentata* seems to respond directly to grazing, *A. quadrilorata*, which exclusively inhabits the cattle-resistant *Eryngium horridum*, is indirectly affected by grazing. Therefore, although grazing may not be the best solution for all kinds of grassland management (Helden et al. 2020), moderate grazing using limited animal load, up to 0.6 AU/ha/year (Jansen et al. 2013; Clendenin 2016; Toupet et al. 2020), seems to have the potential to maintain and preserve spiders and other generalist predators that are intrinsically linked to vegetation structure. Moreover, the ability of these spiders to disperse through ballooning offers a potential survival strategy in the face of grazing pressure, but it presents challenges related to identifying suitable habitats. This highlights promising avenues for future research, particularly regarding how light incidence and shade coverage influence habitat choice for orb-weaver spiders. In light of these findings, it is imperative that we strive for an optimal balance in managing grazing practices and conserving habitats to ensure the survival of orb-weaver spiders and other arthropods. Recognizing the value of these small but ecologically significant creatures is essential for maintaining the health and stability of our ecosystems and the services they provide (Meadows et al. 2017; Fernández-Tizón et al. 2020).

## Acknowledgements

We are grateful for CAPES and the scholarship granted to the first author (process #88882.439370/2019-01). We are also grateful for Neotropical Grassland Conservancy and the Student Grant granted to the first author. The authors declare that they have no known competing financial interests or personal relationships that could have appeared to influence the work reported in this paper.

## Author contributions

All authors contributed to the study conception and design. All authors performed the screening and analysis of the data, as well as the writing and revision of the text, tables, and images. Finally, all authors read and approved the final manuscript.

## Data availability statement

The data that supports the findings of this study are available in the supplementary material of this article.

## Supporting materials

## SUPPLEMENTARY MATERIAL

**Figure 1.**
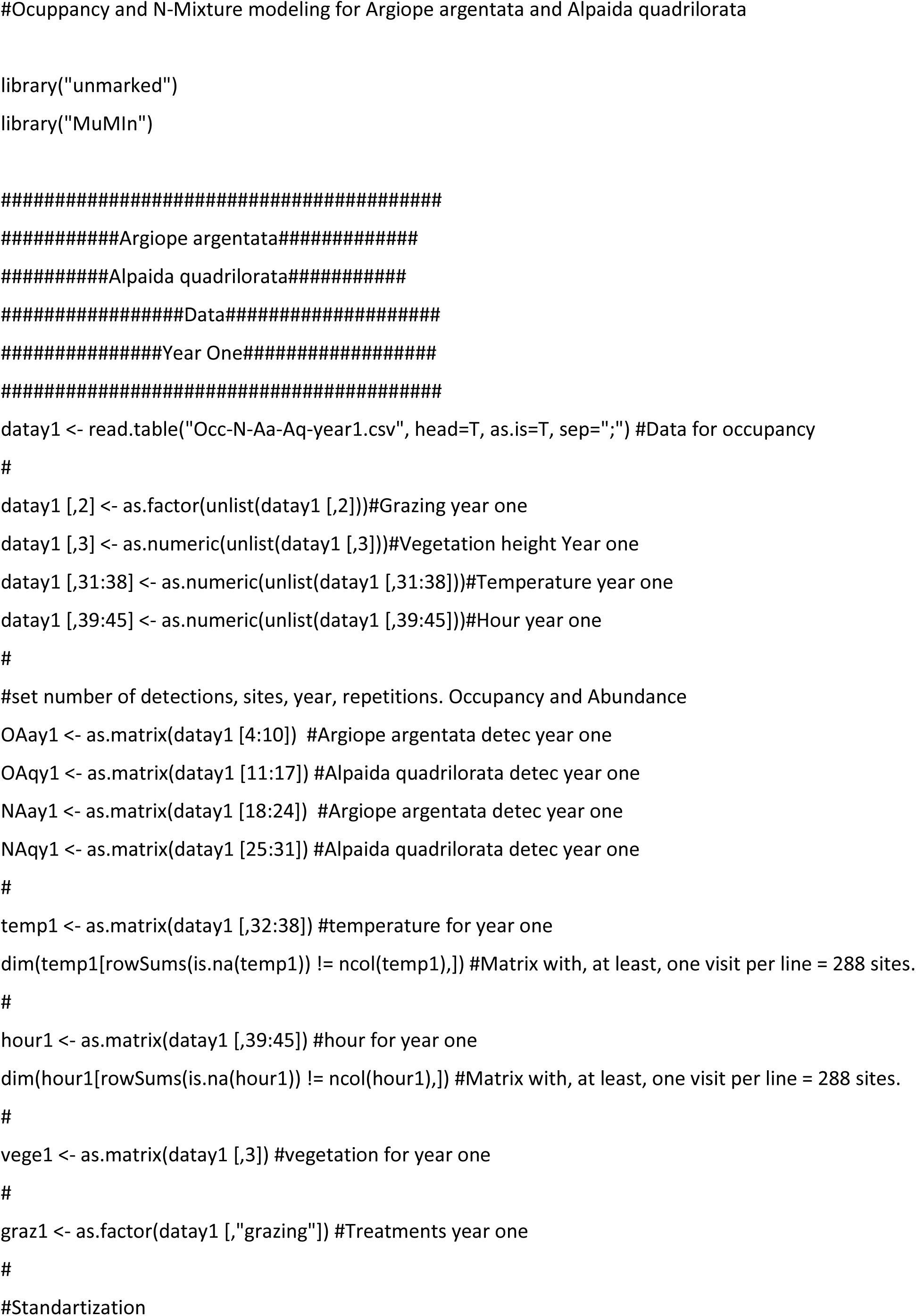
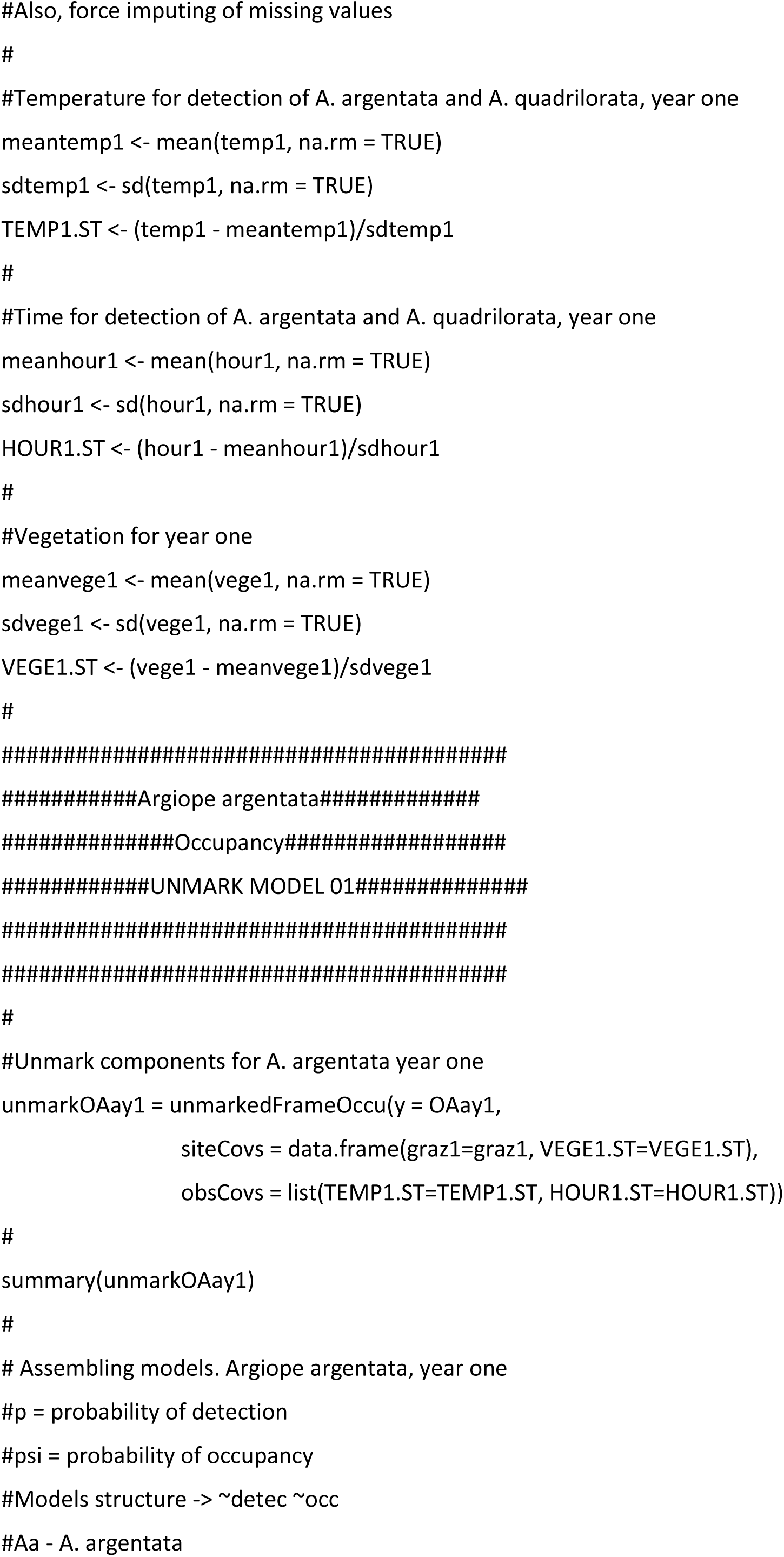
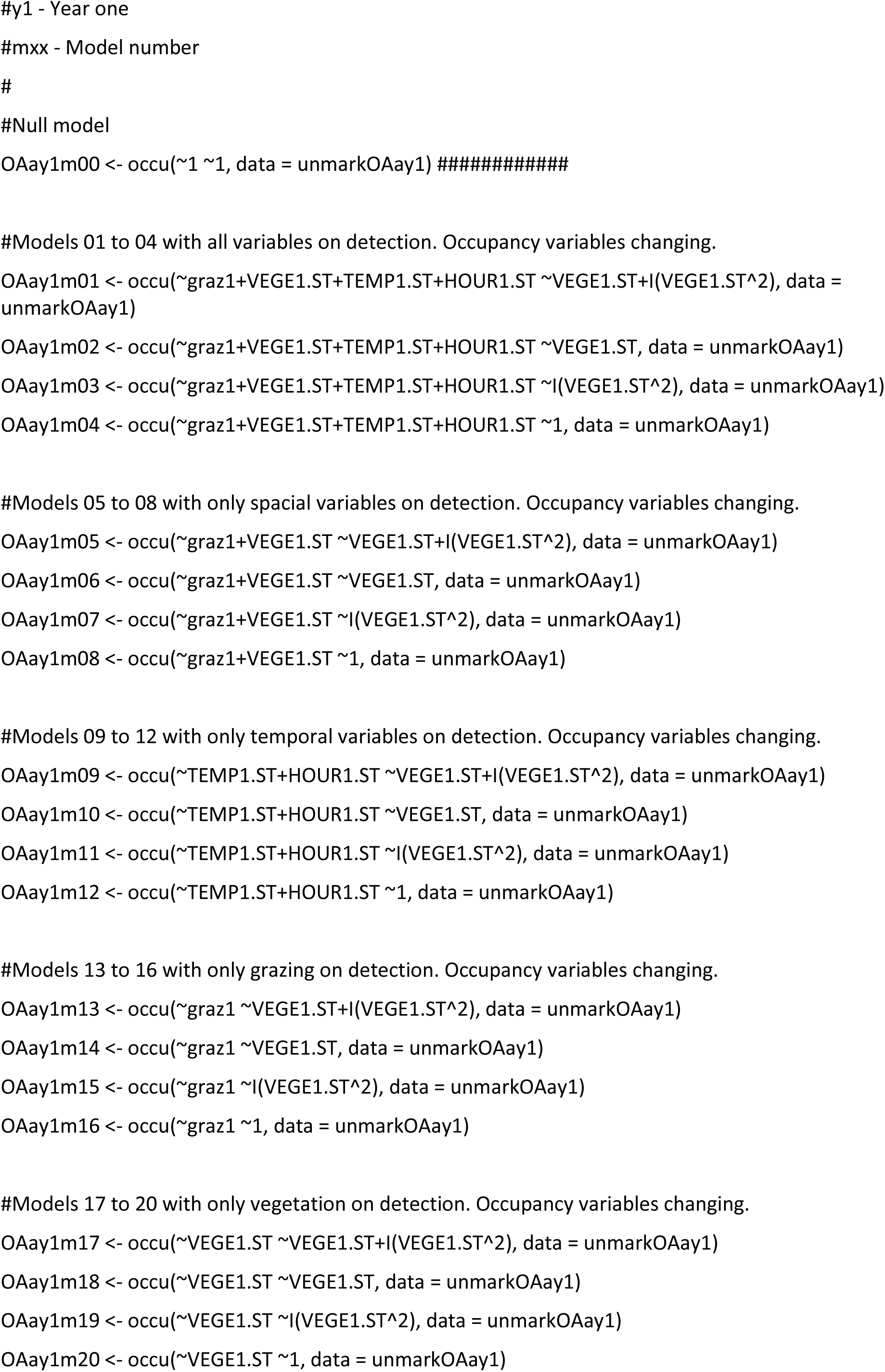
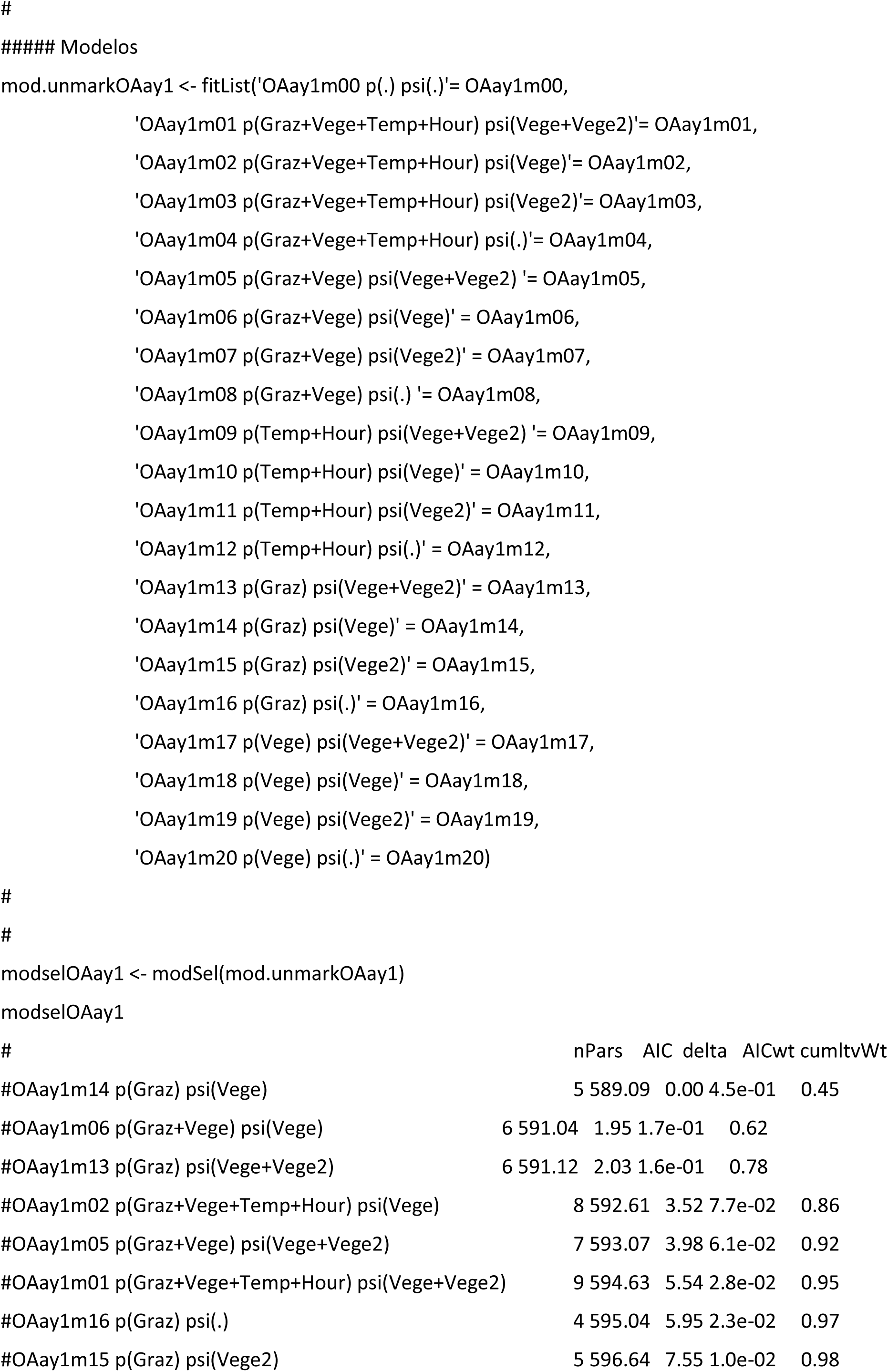
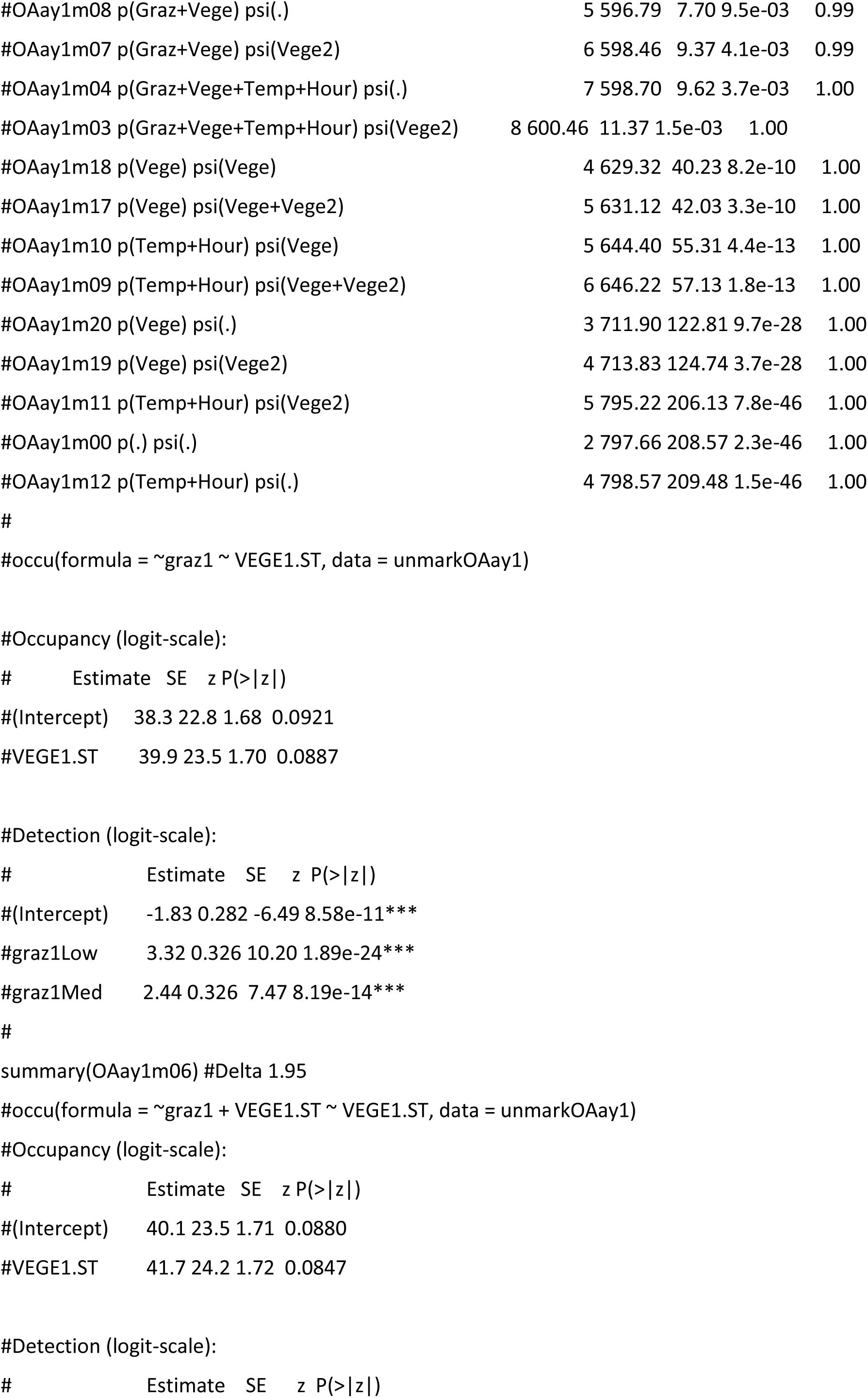
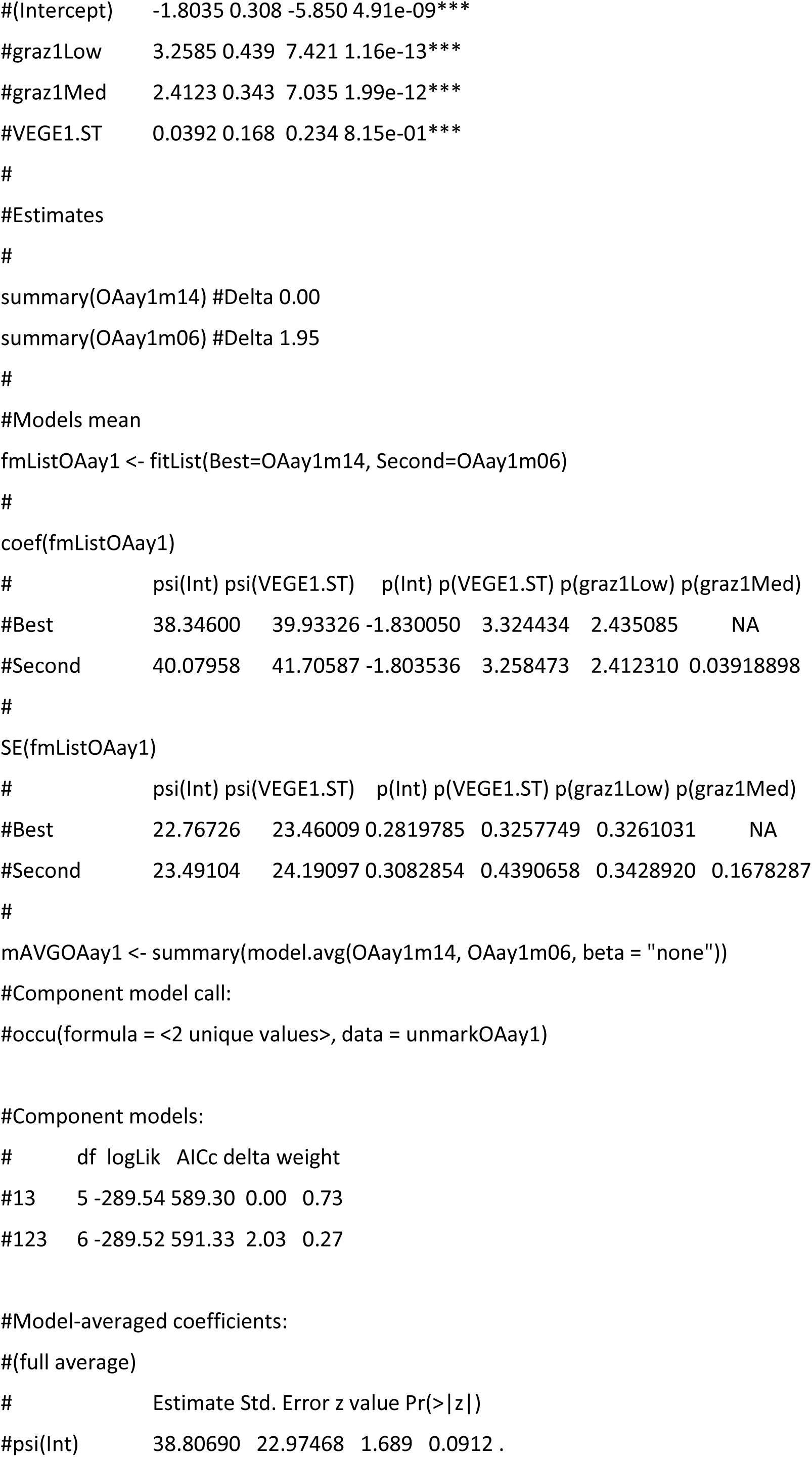
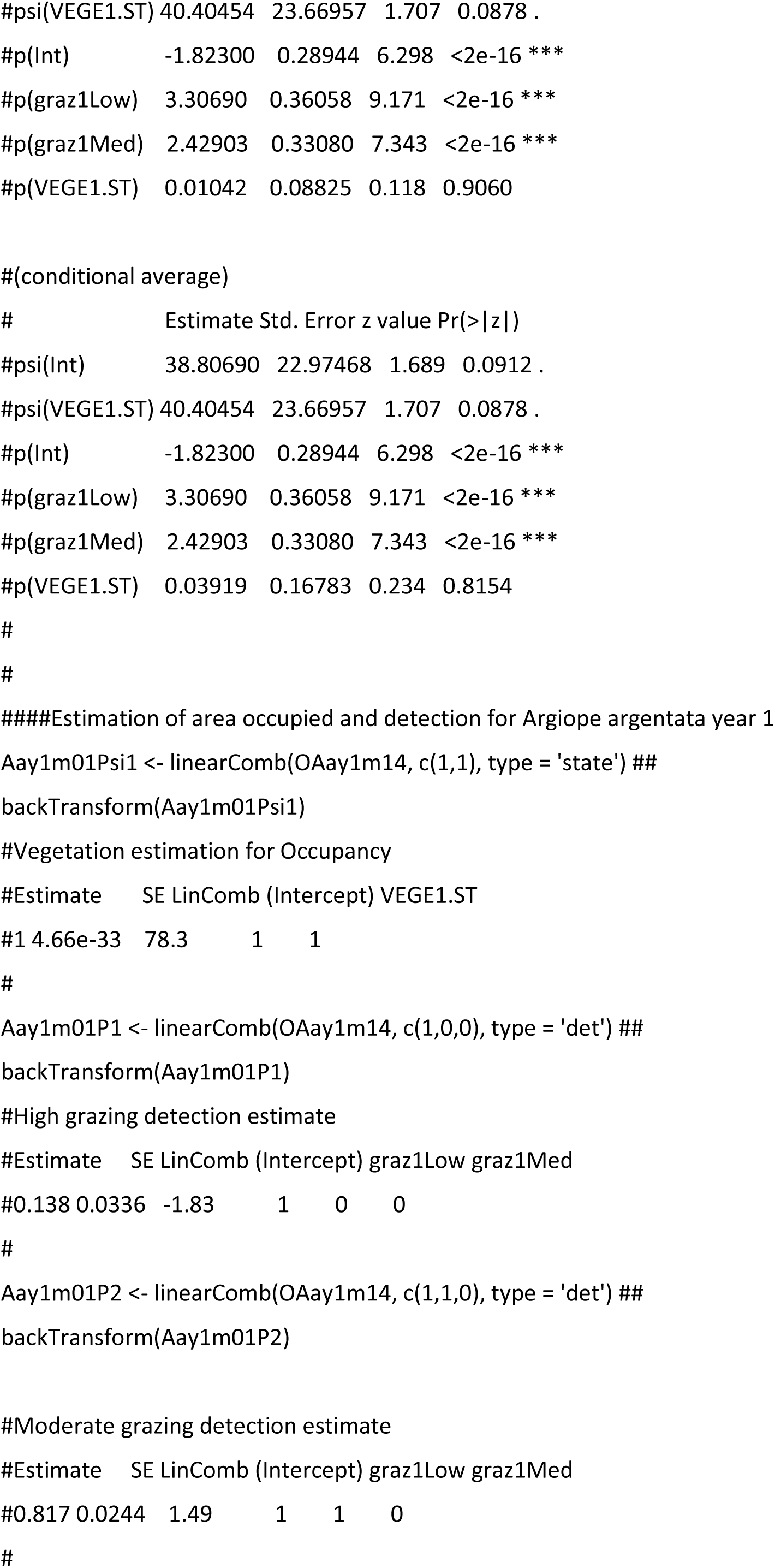
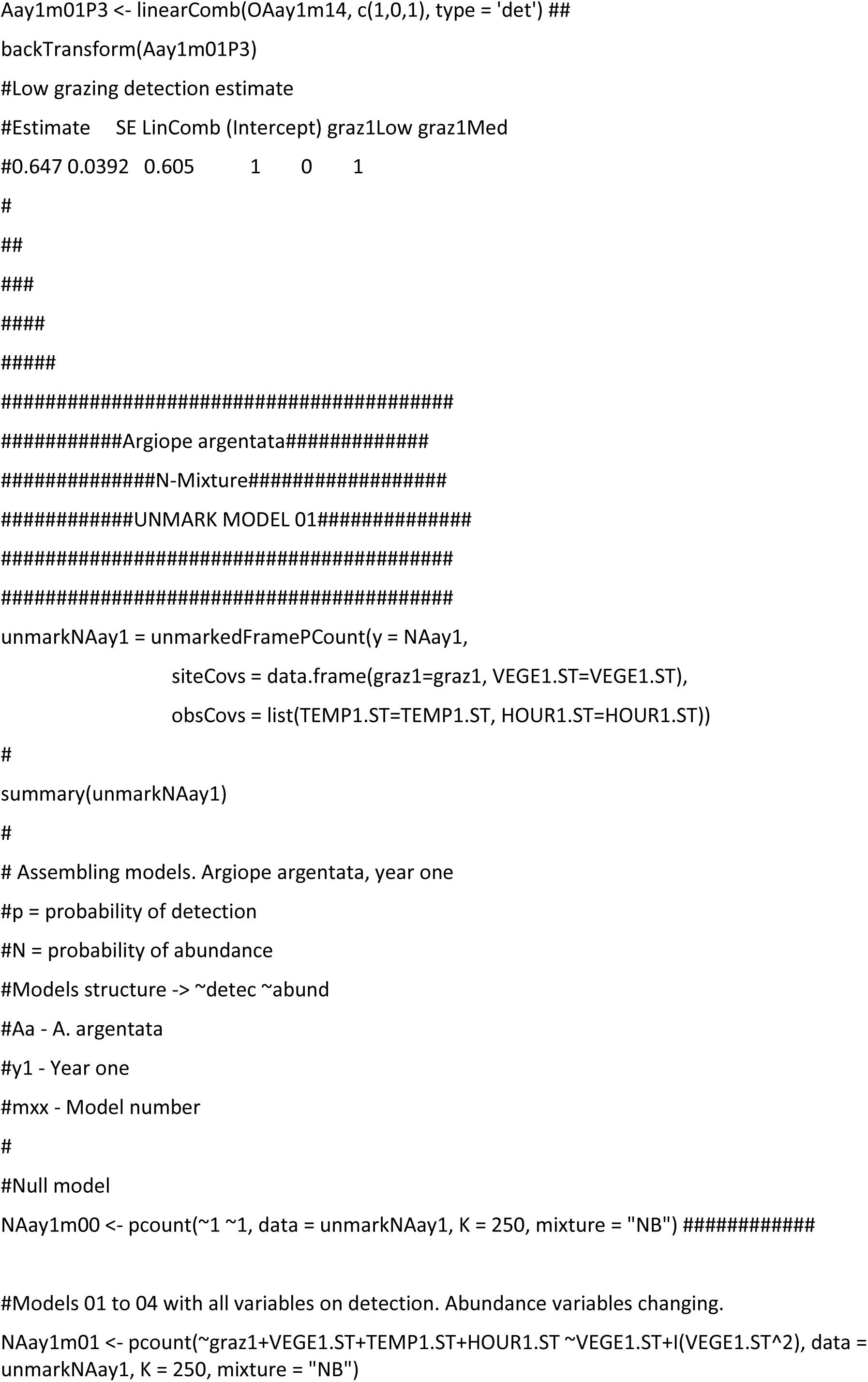
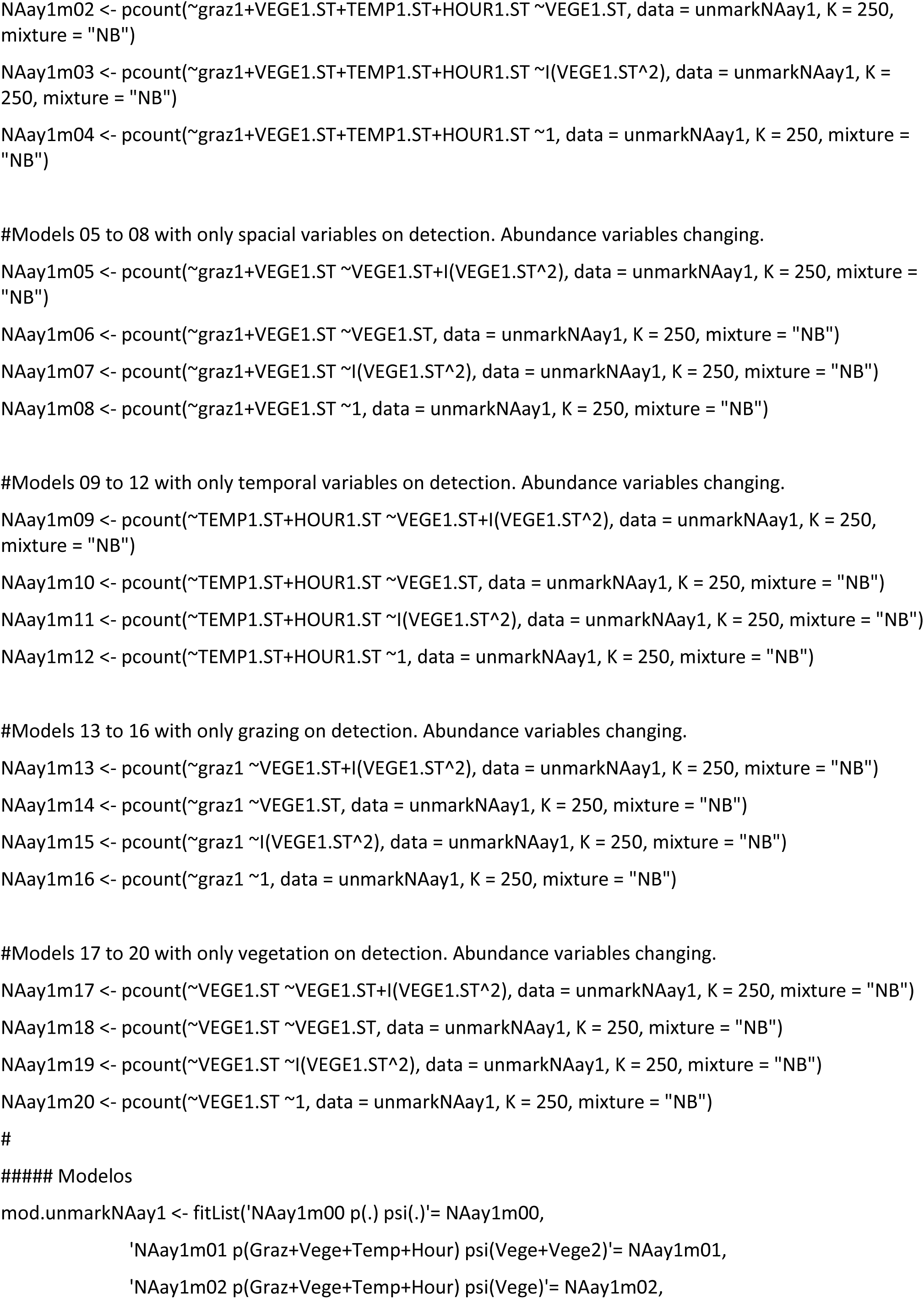
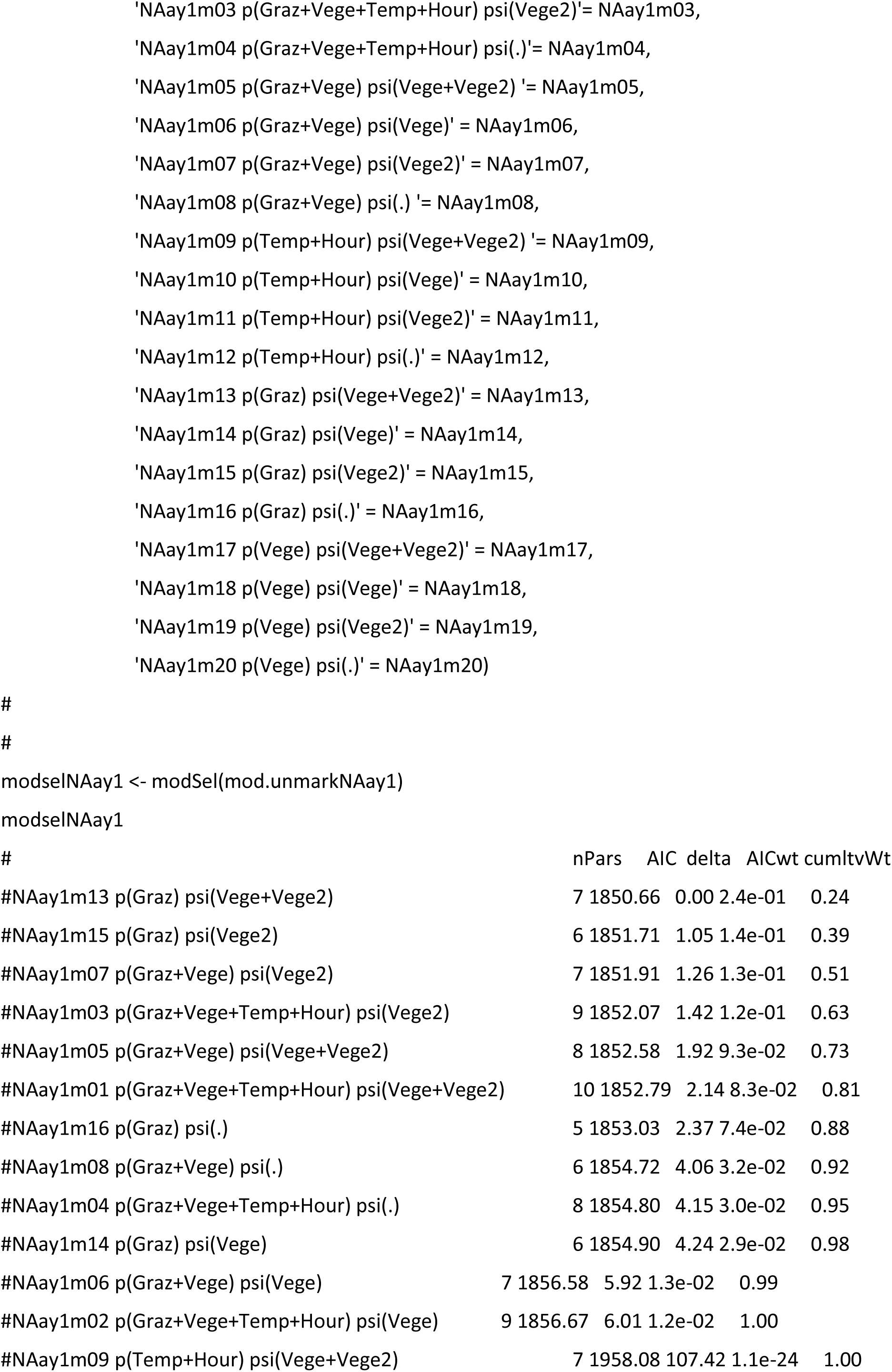
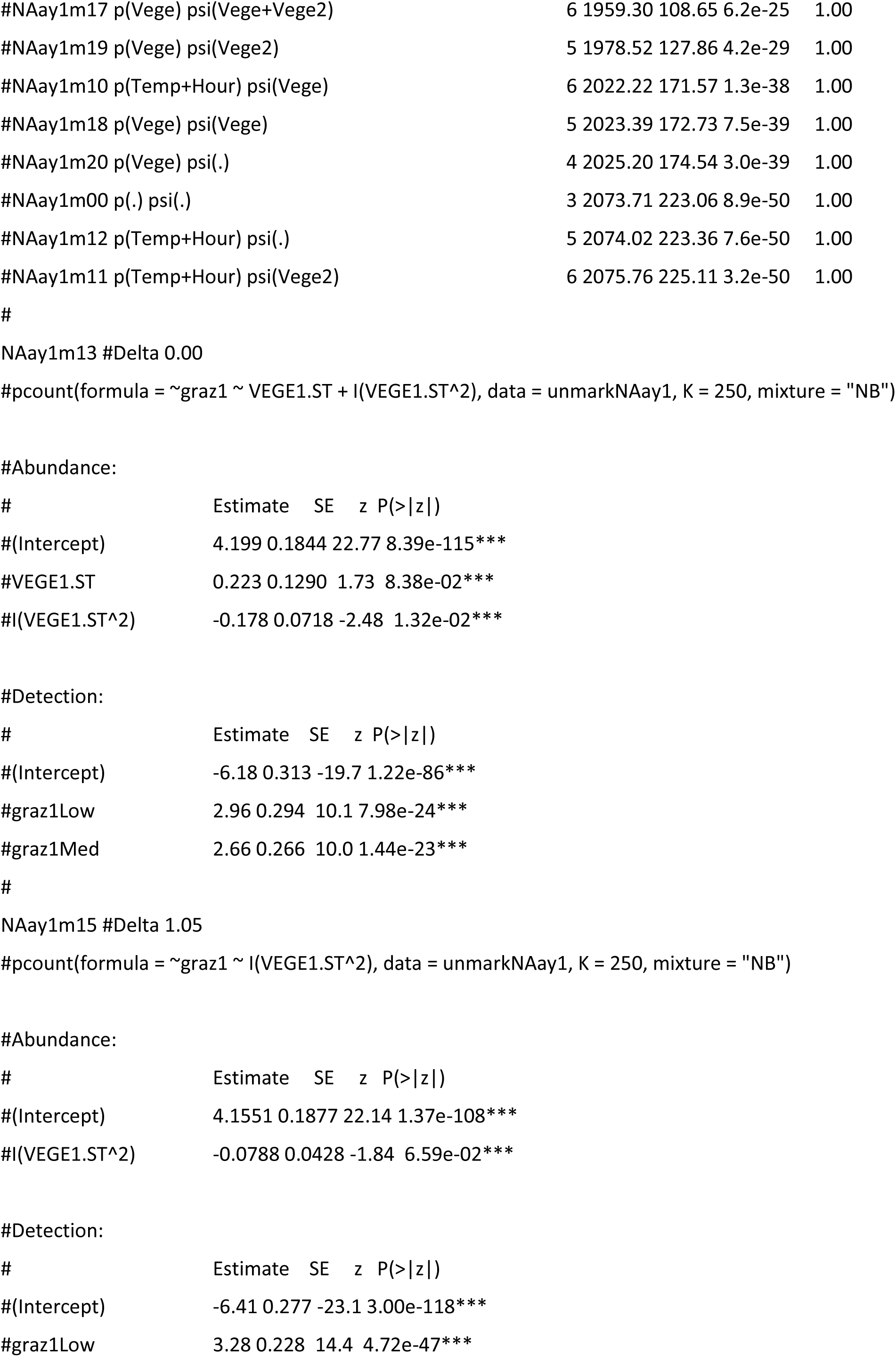
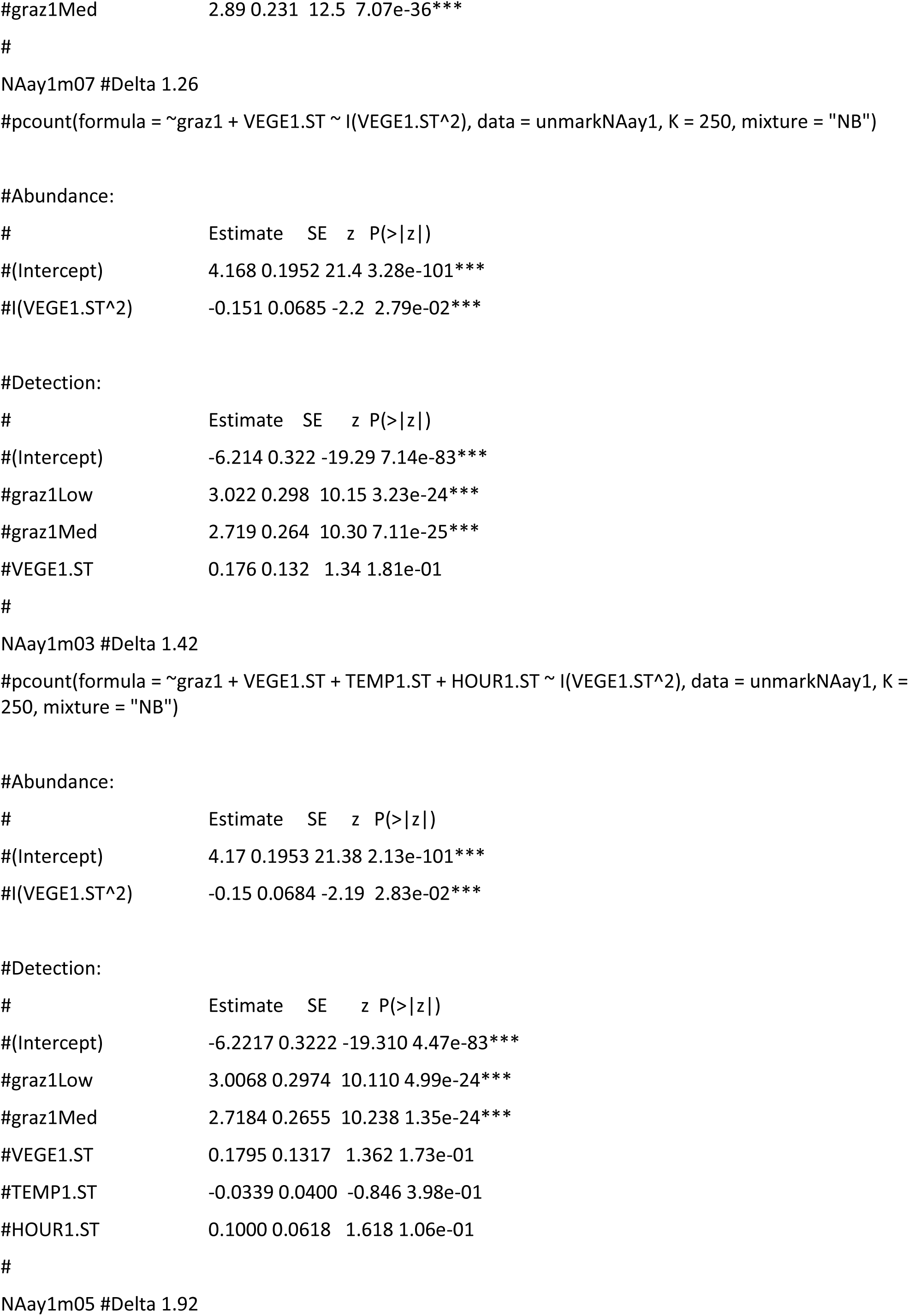
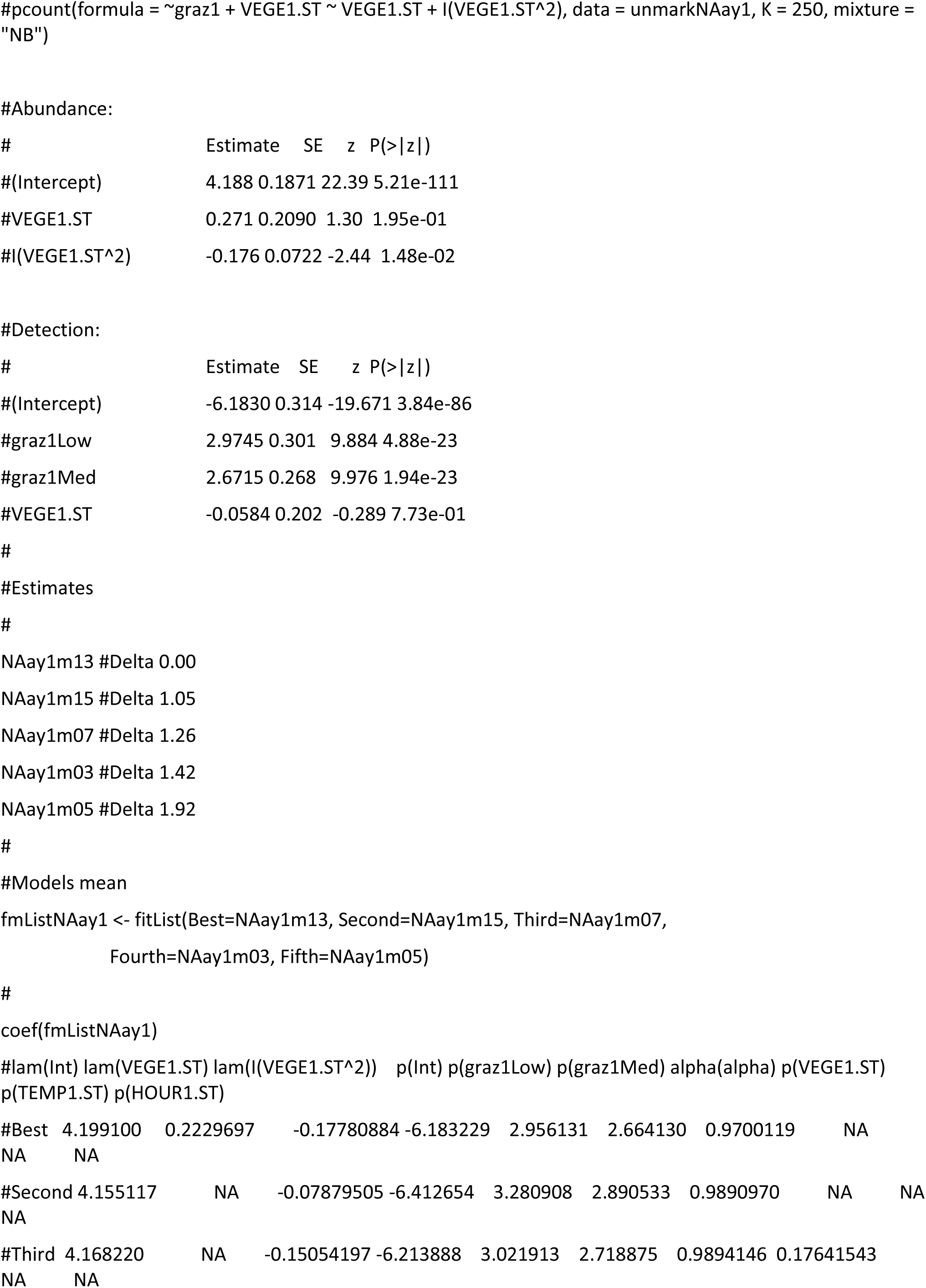
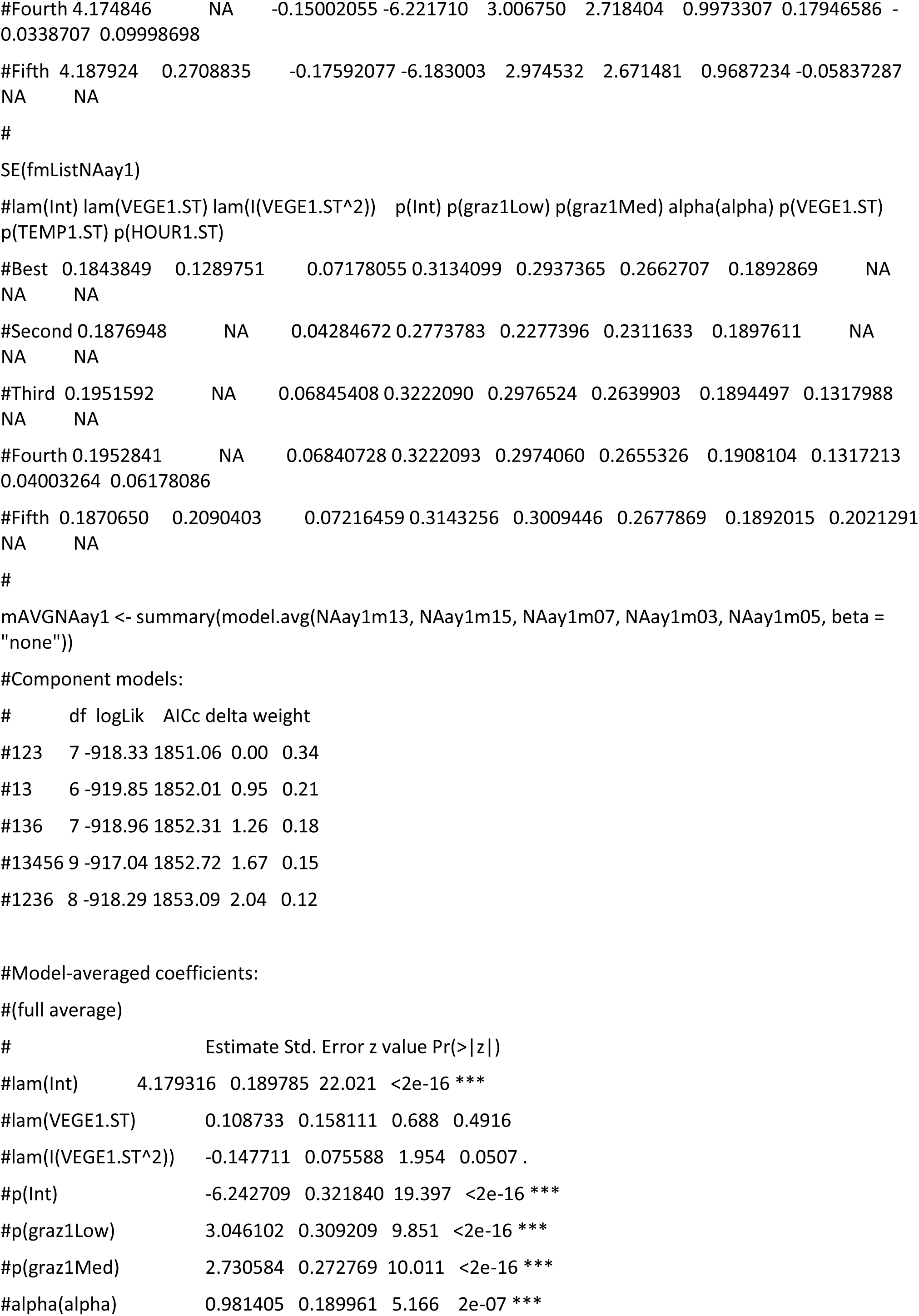
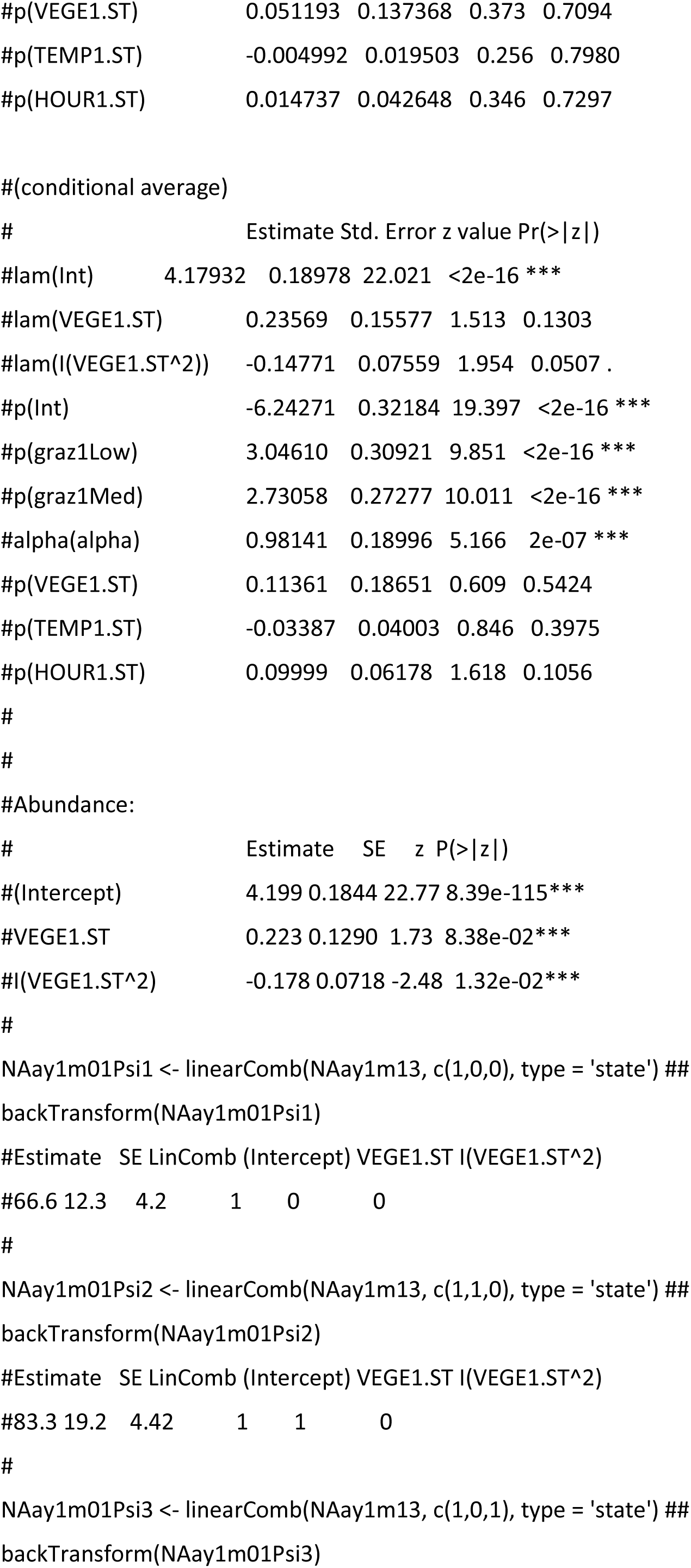
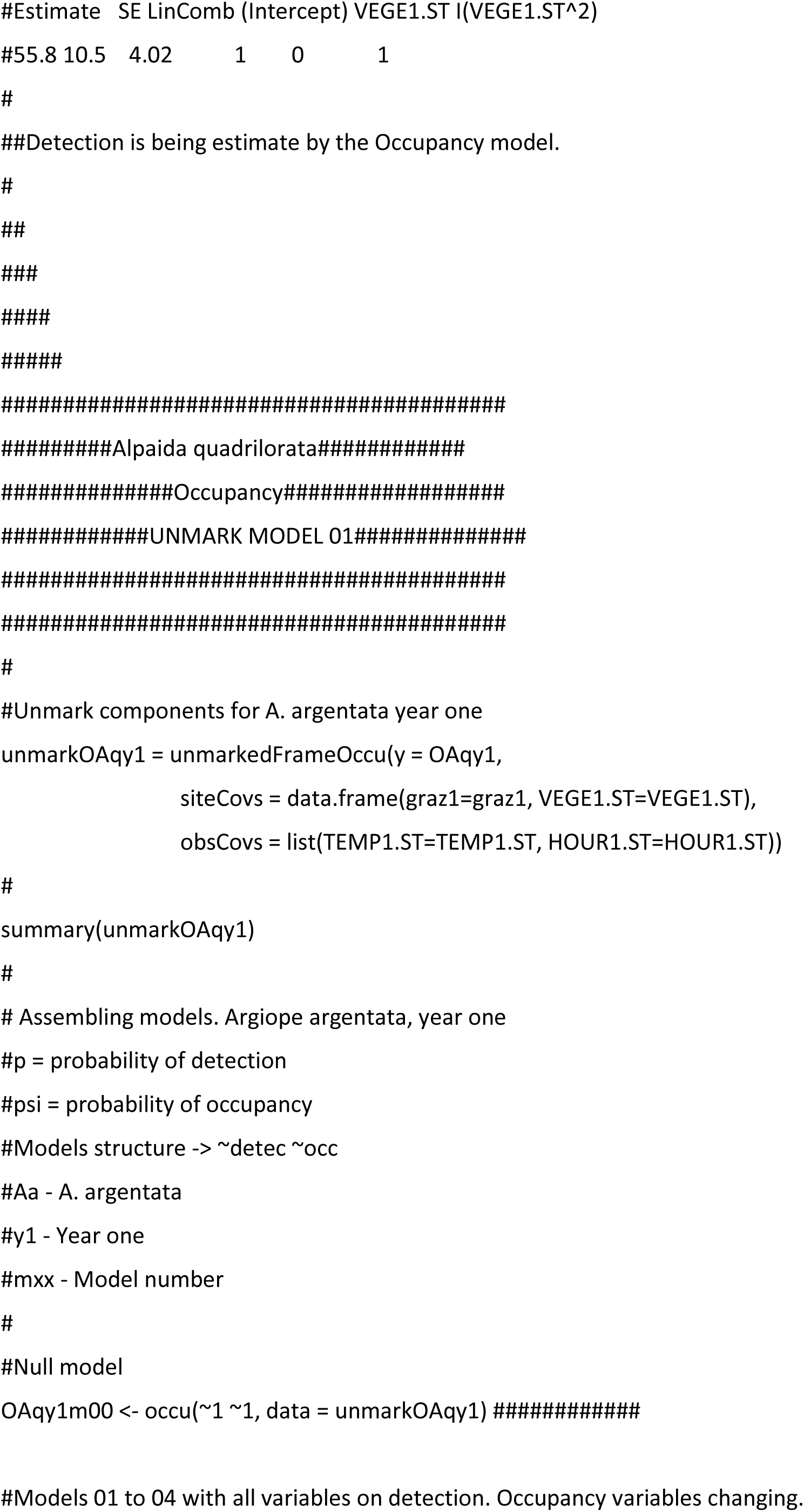
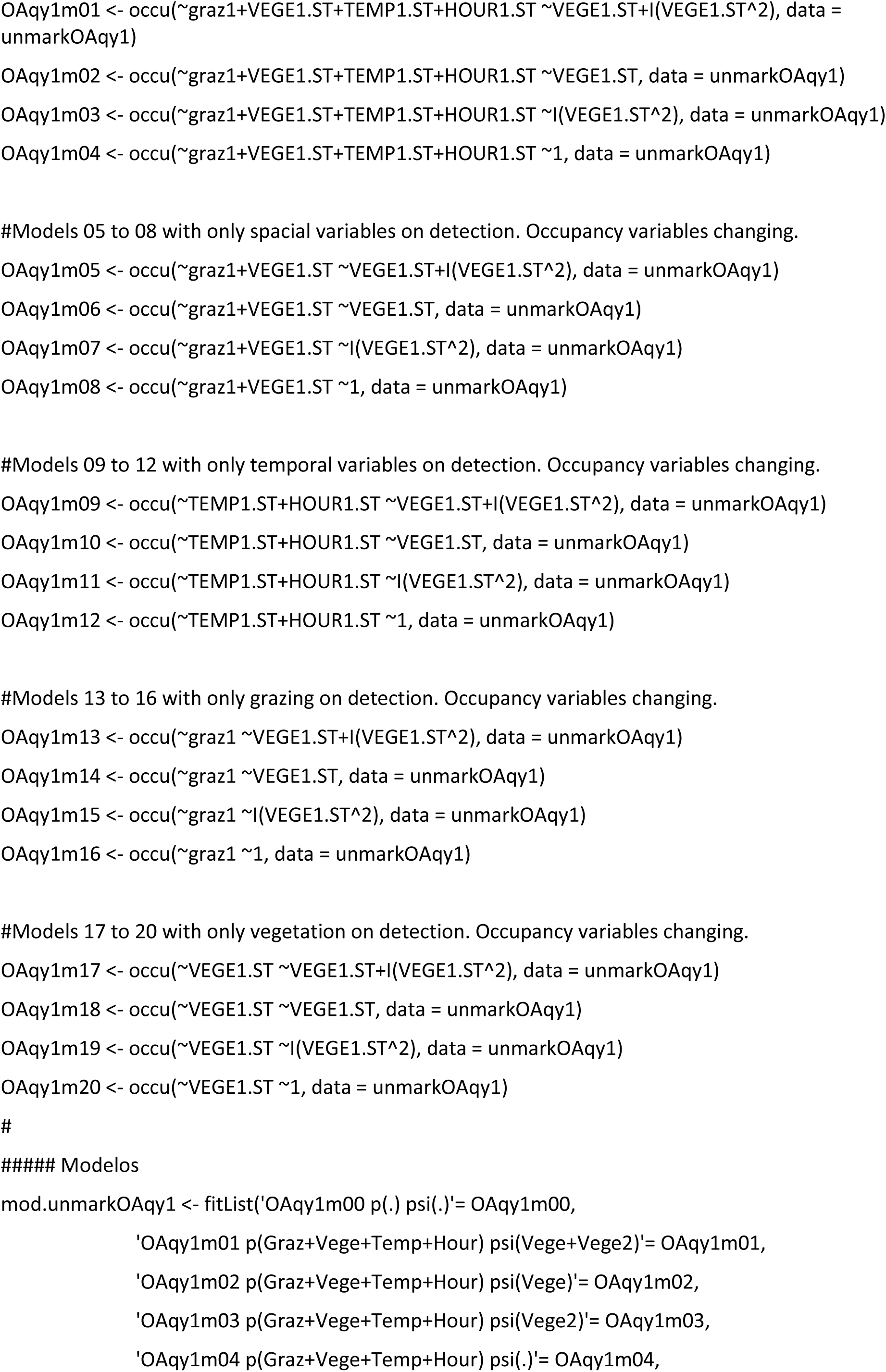
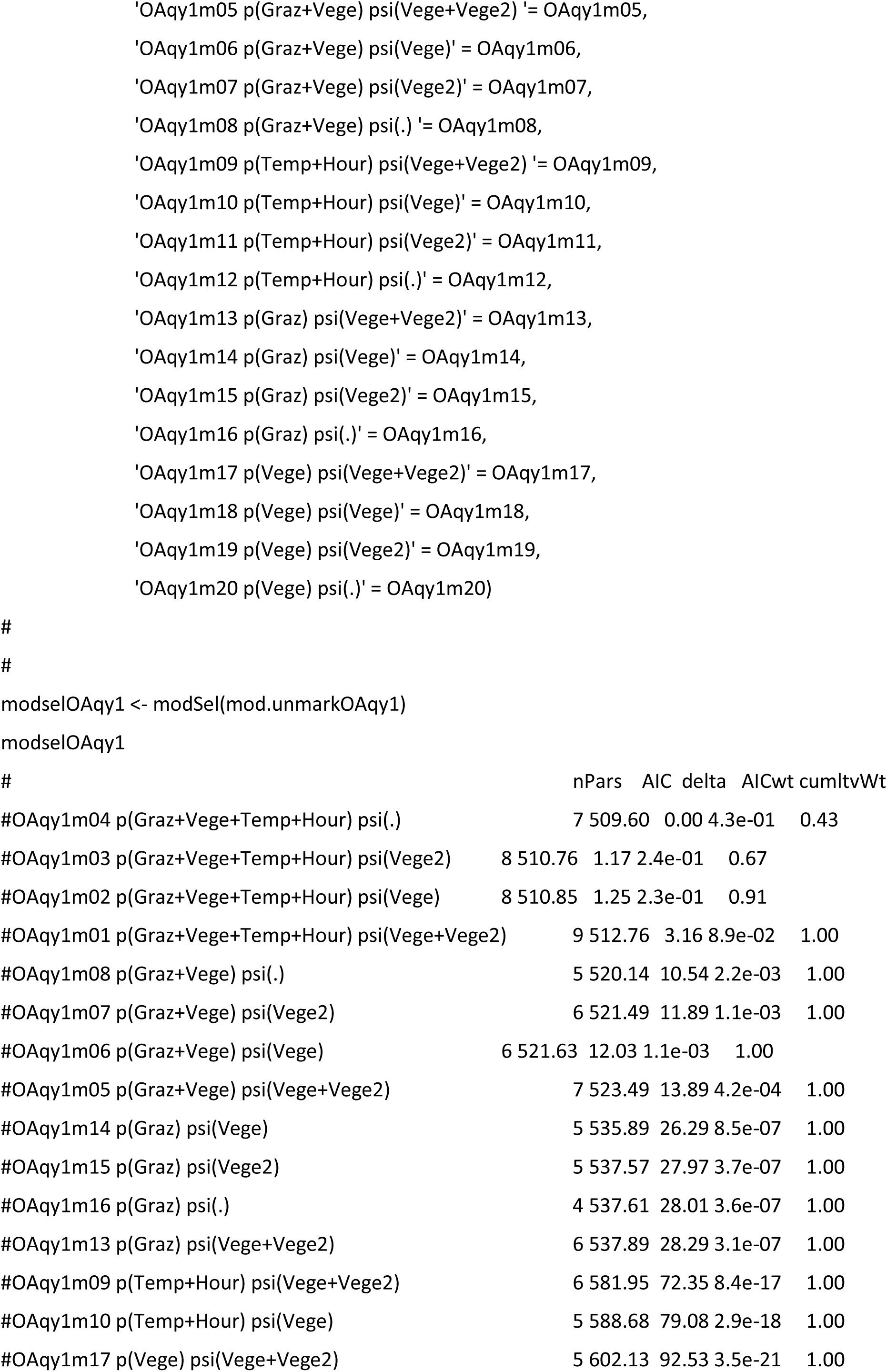
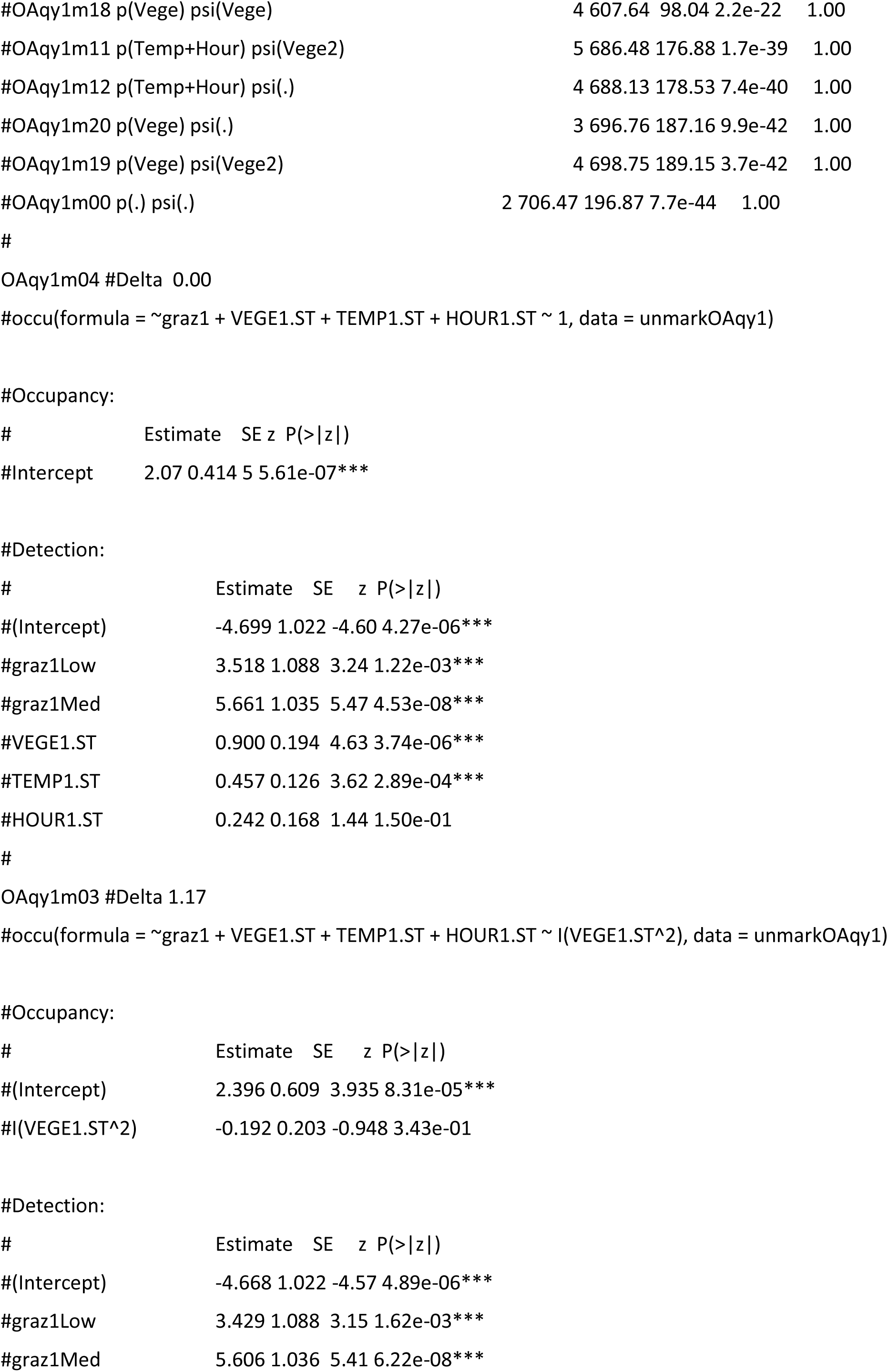
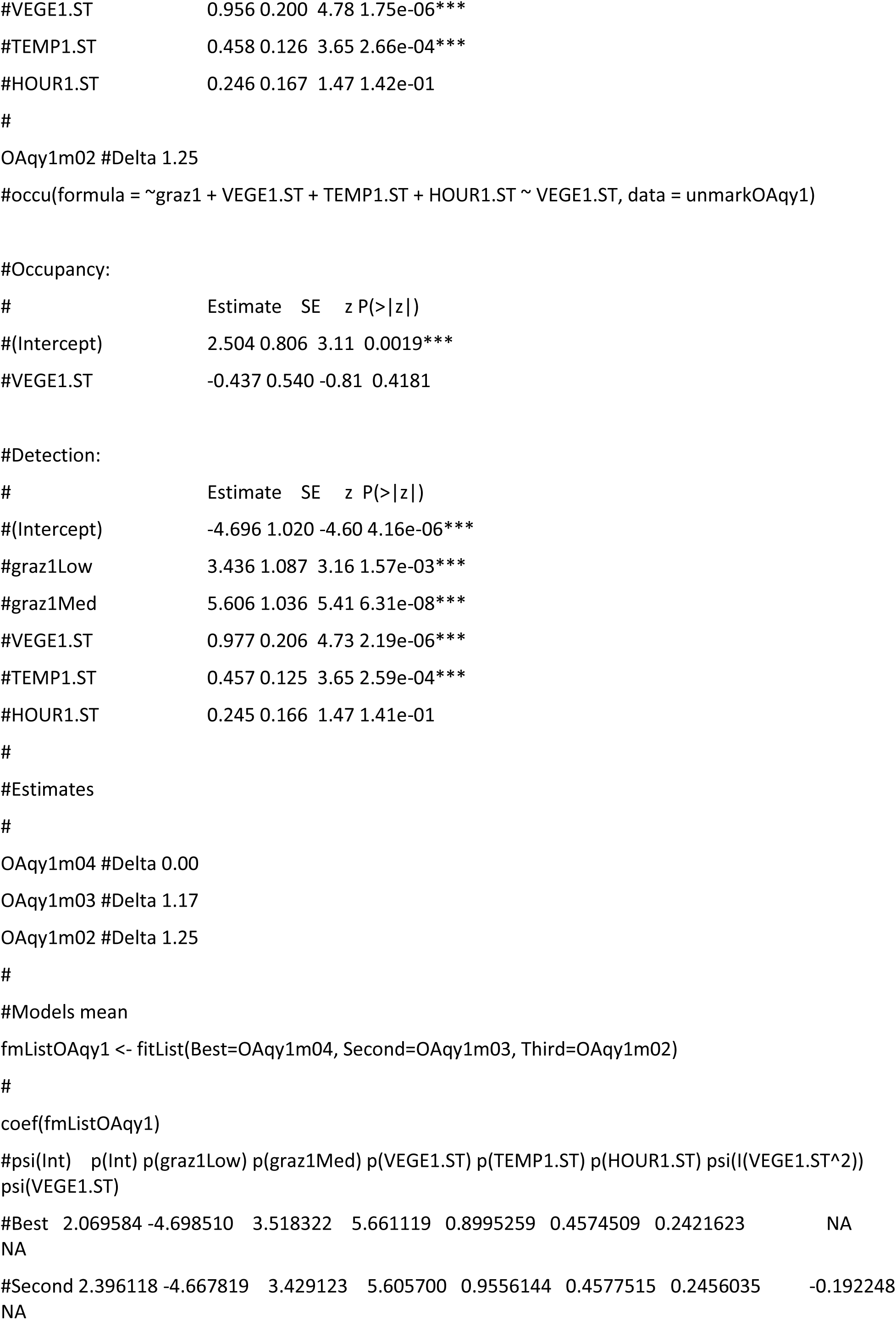
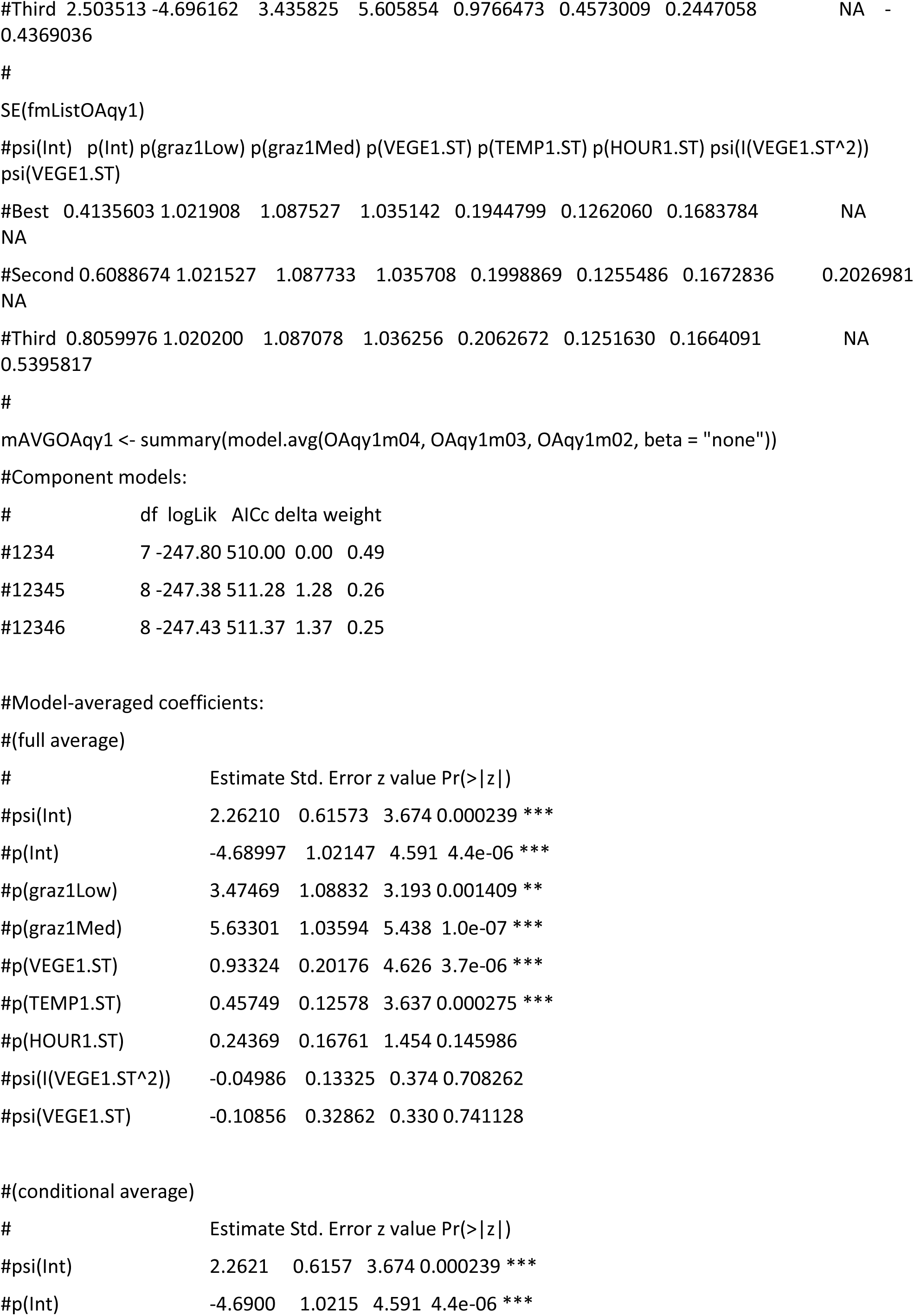
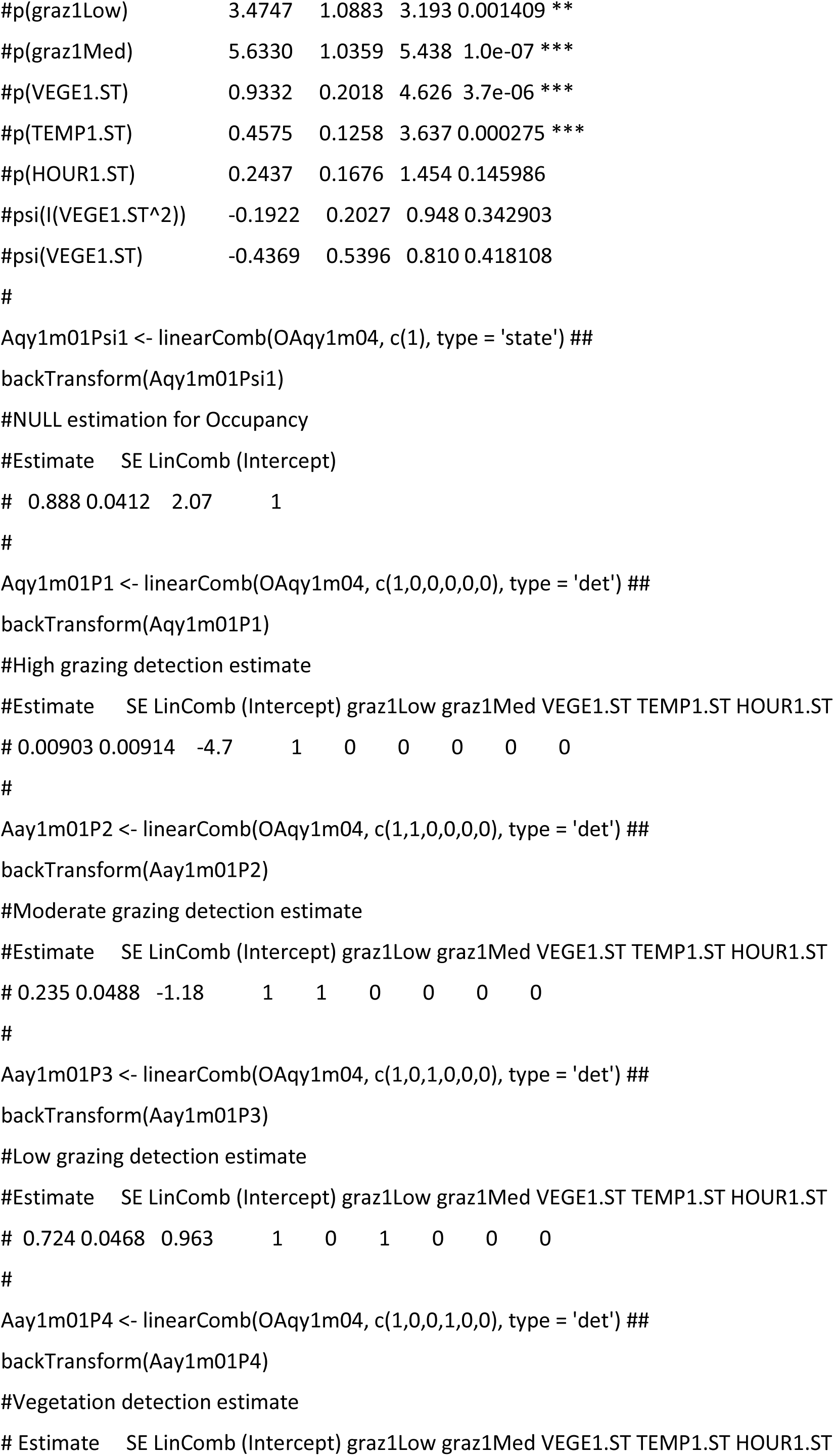
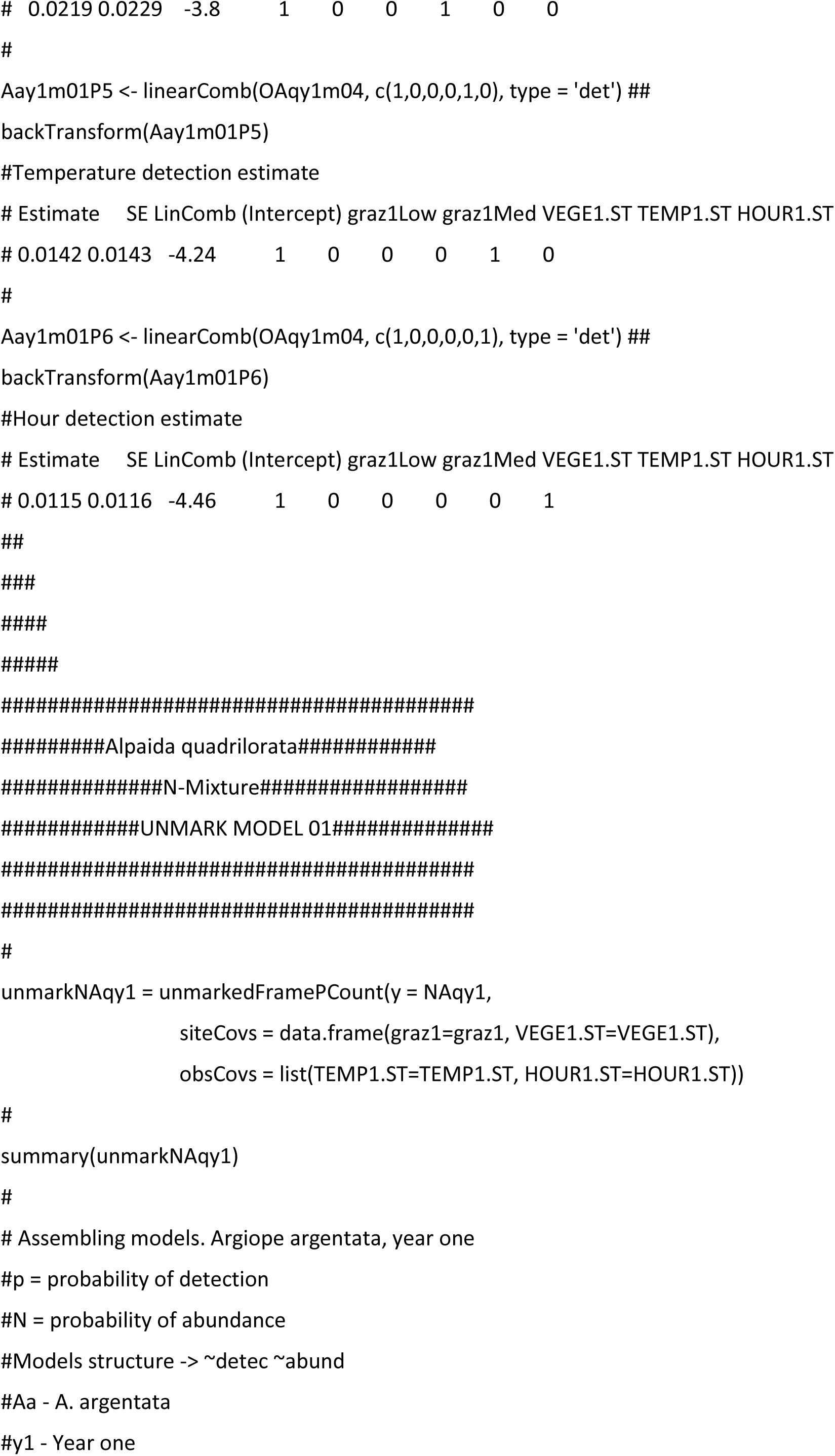
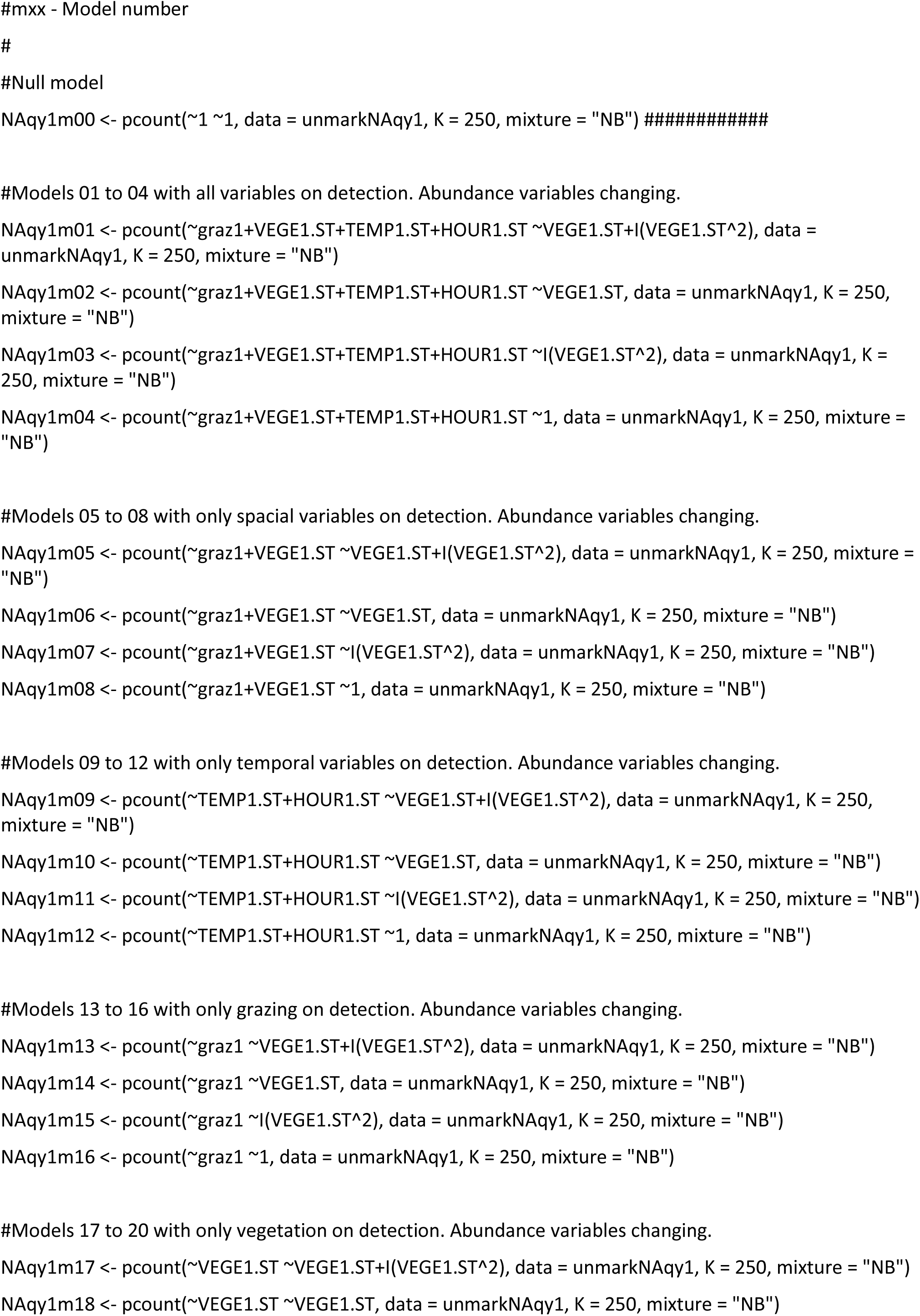
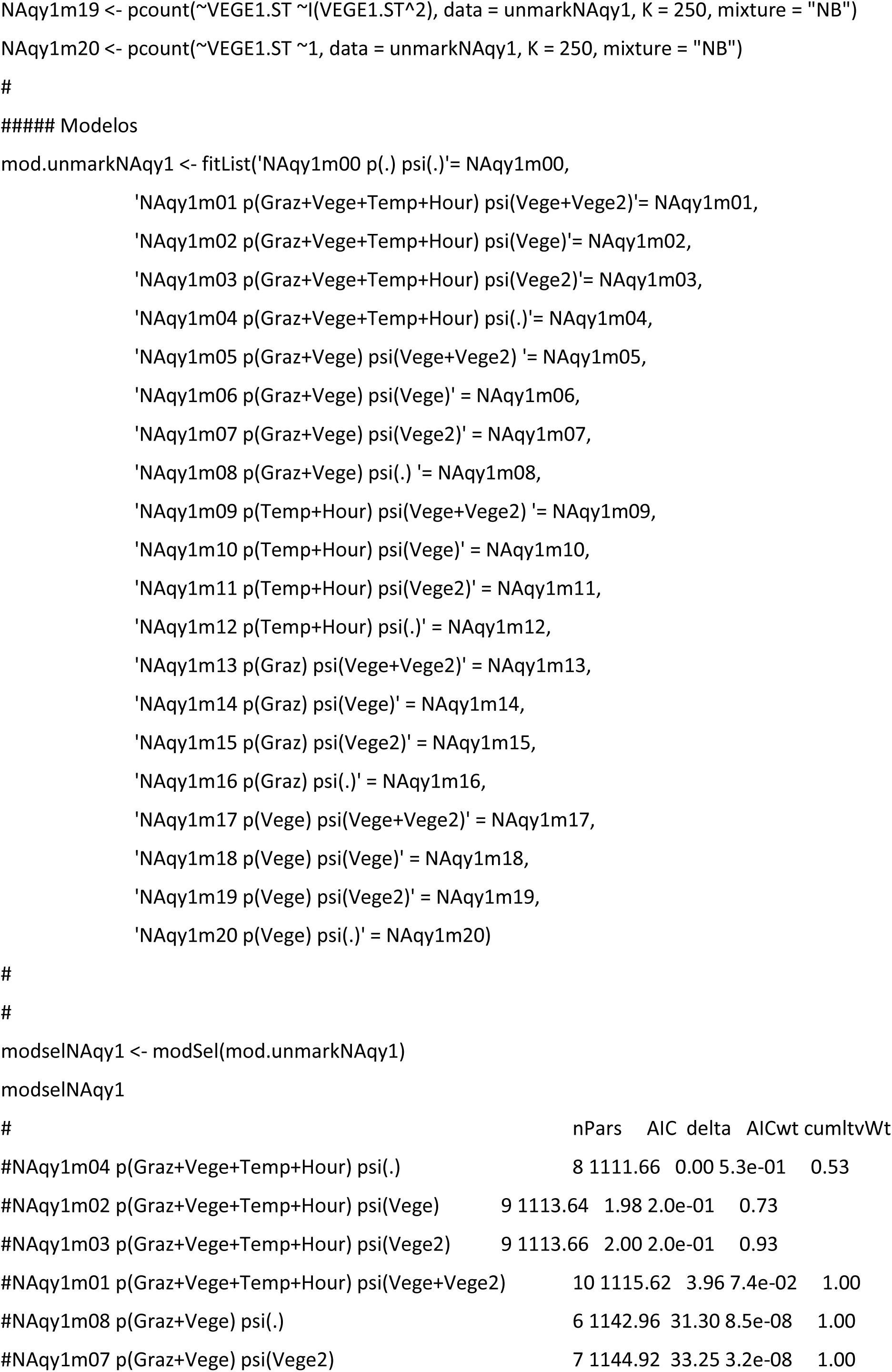
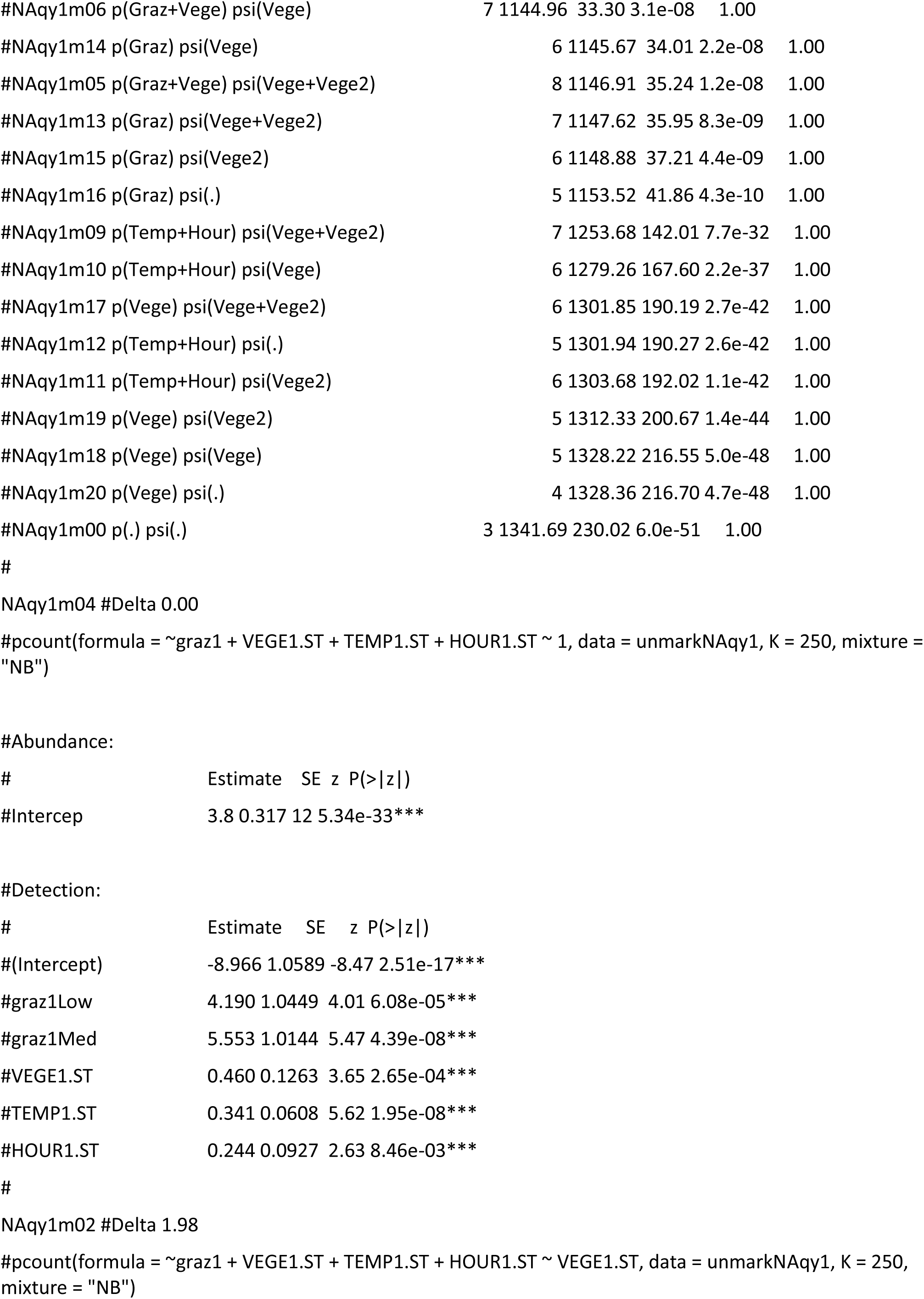
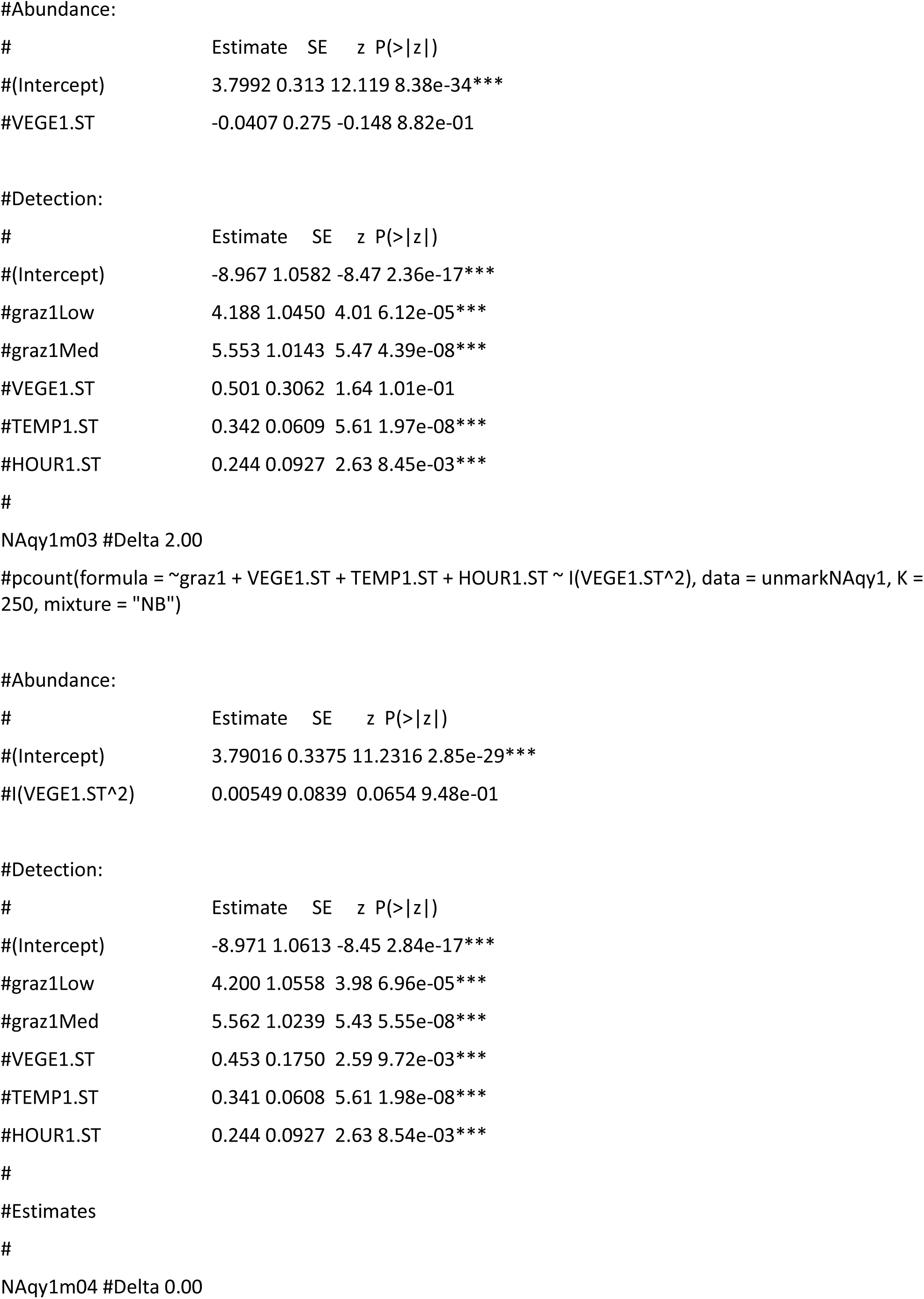
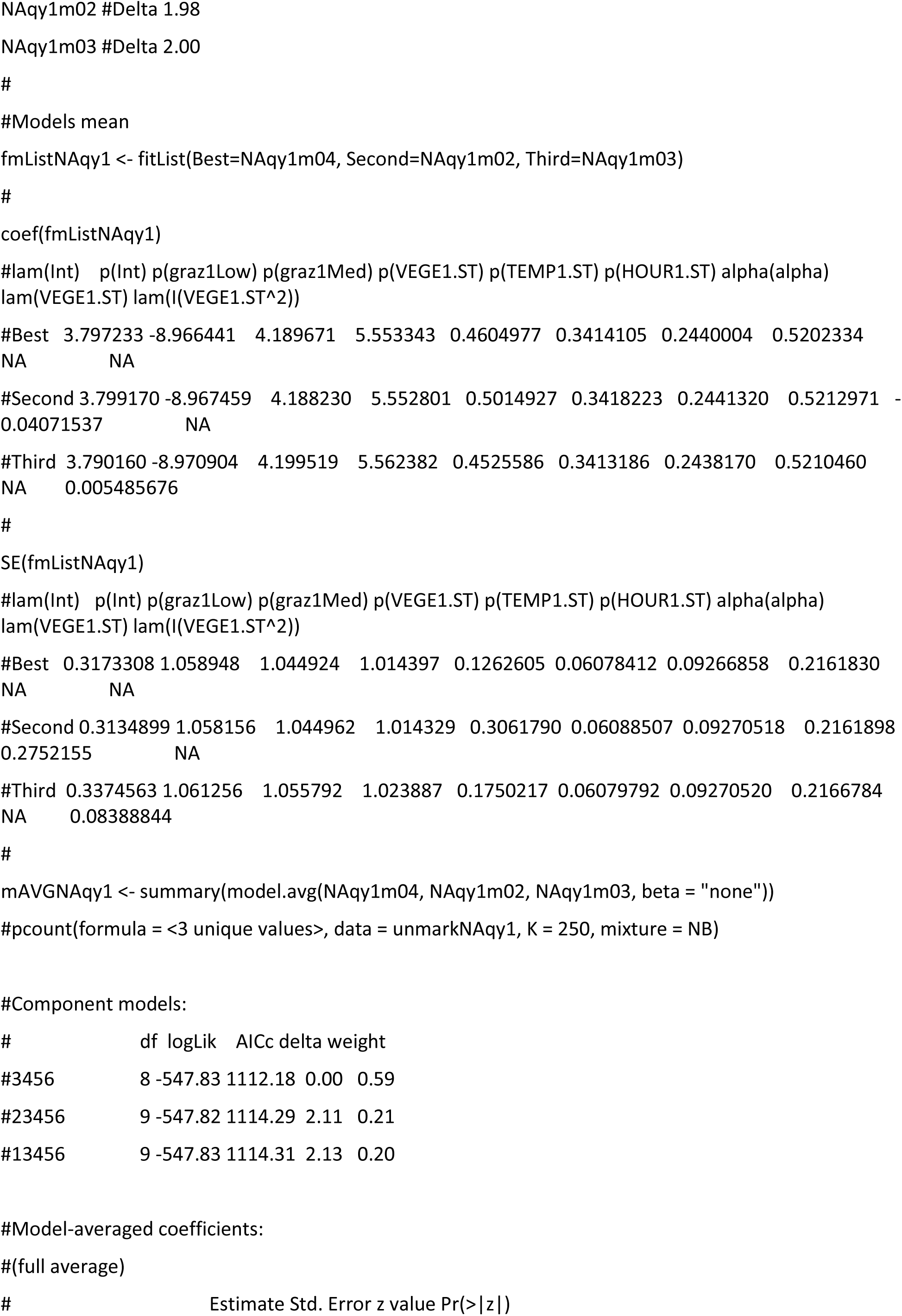
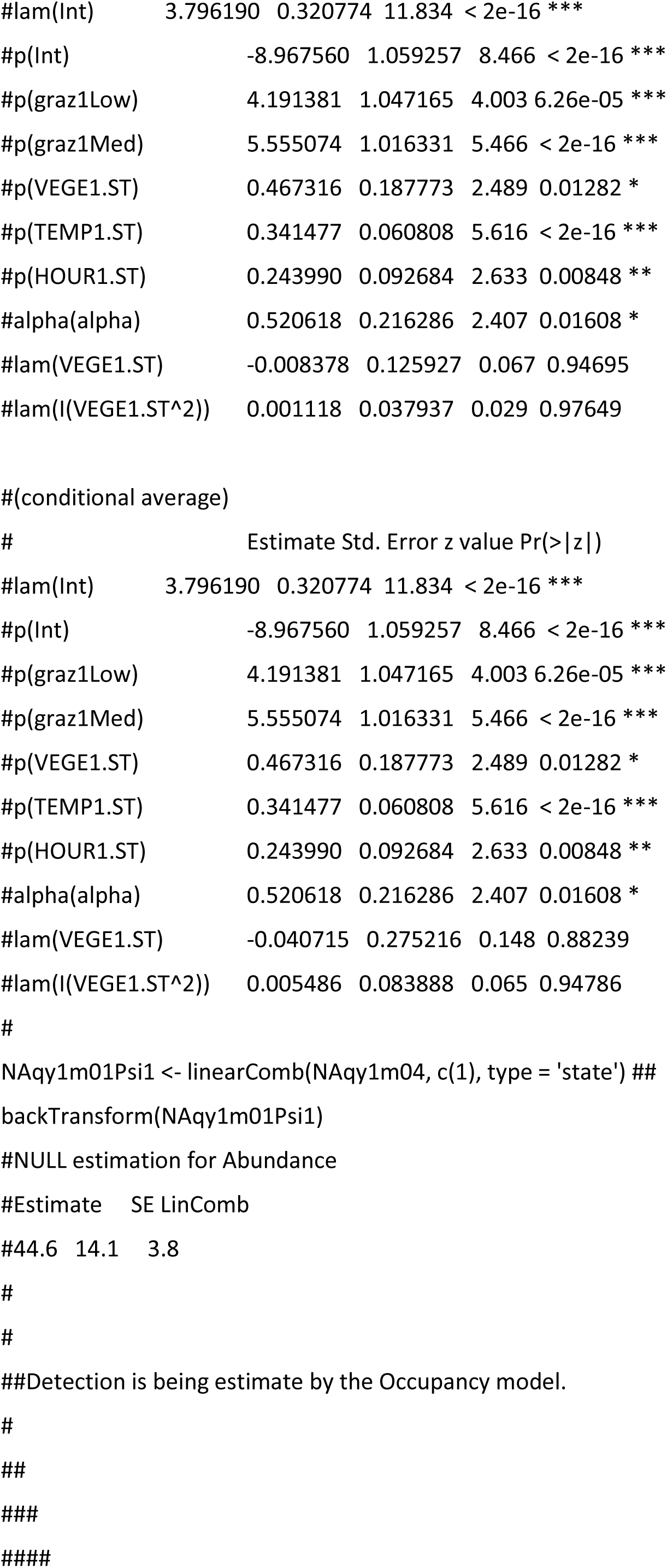
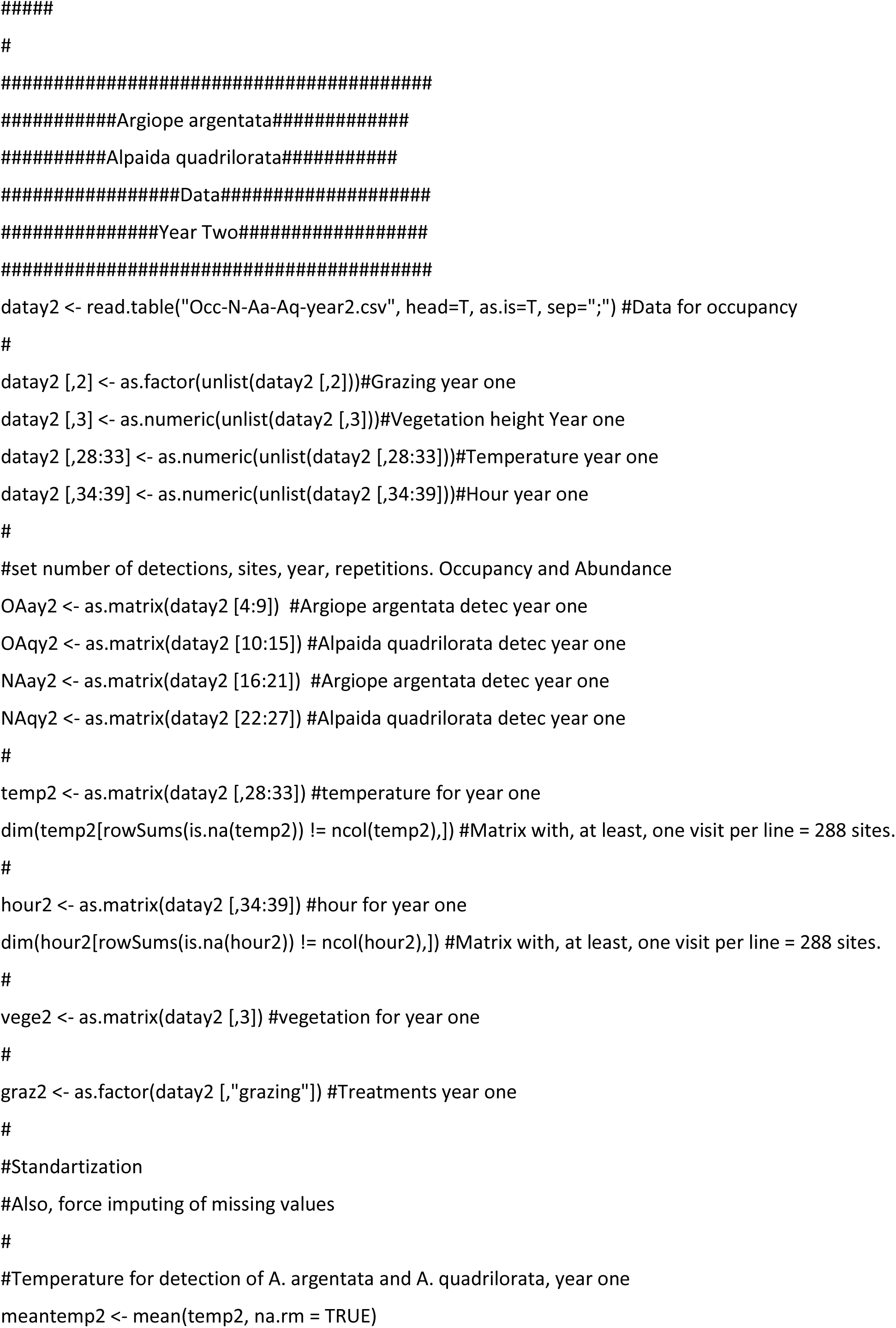
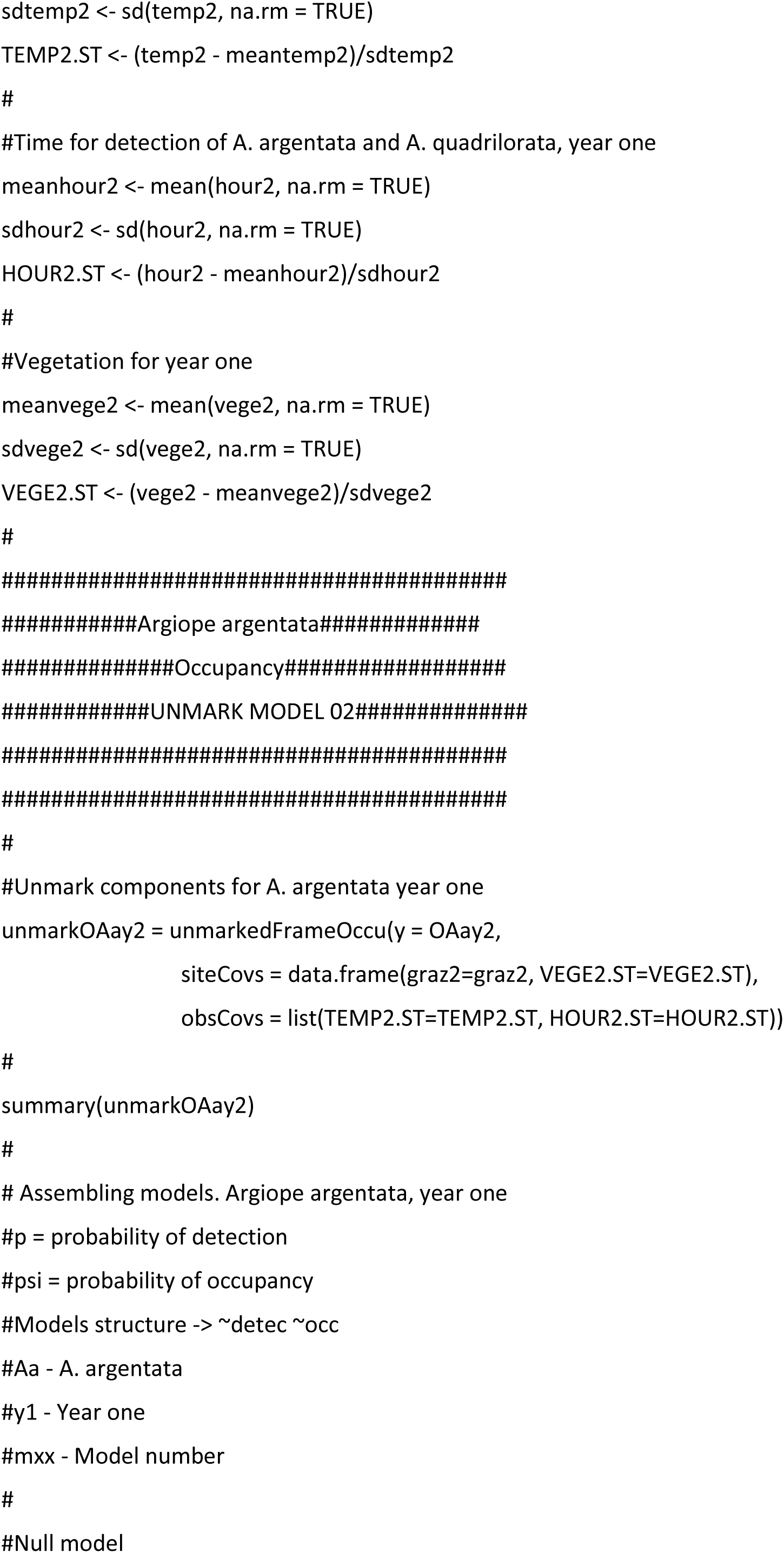
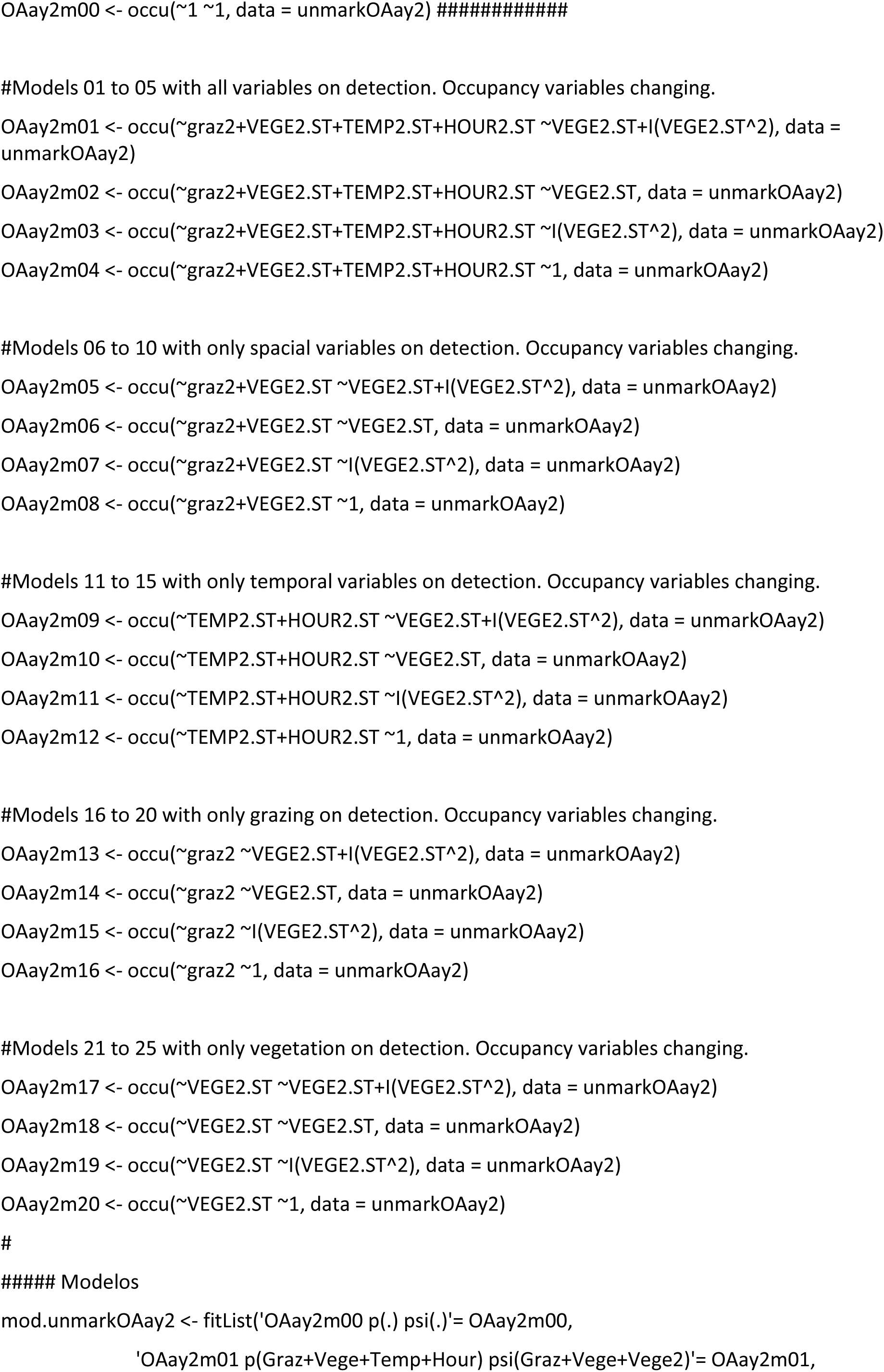
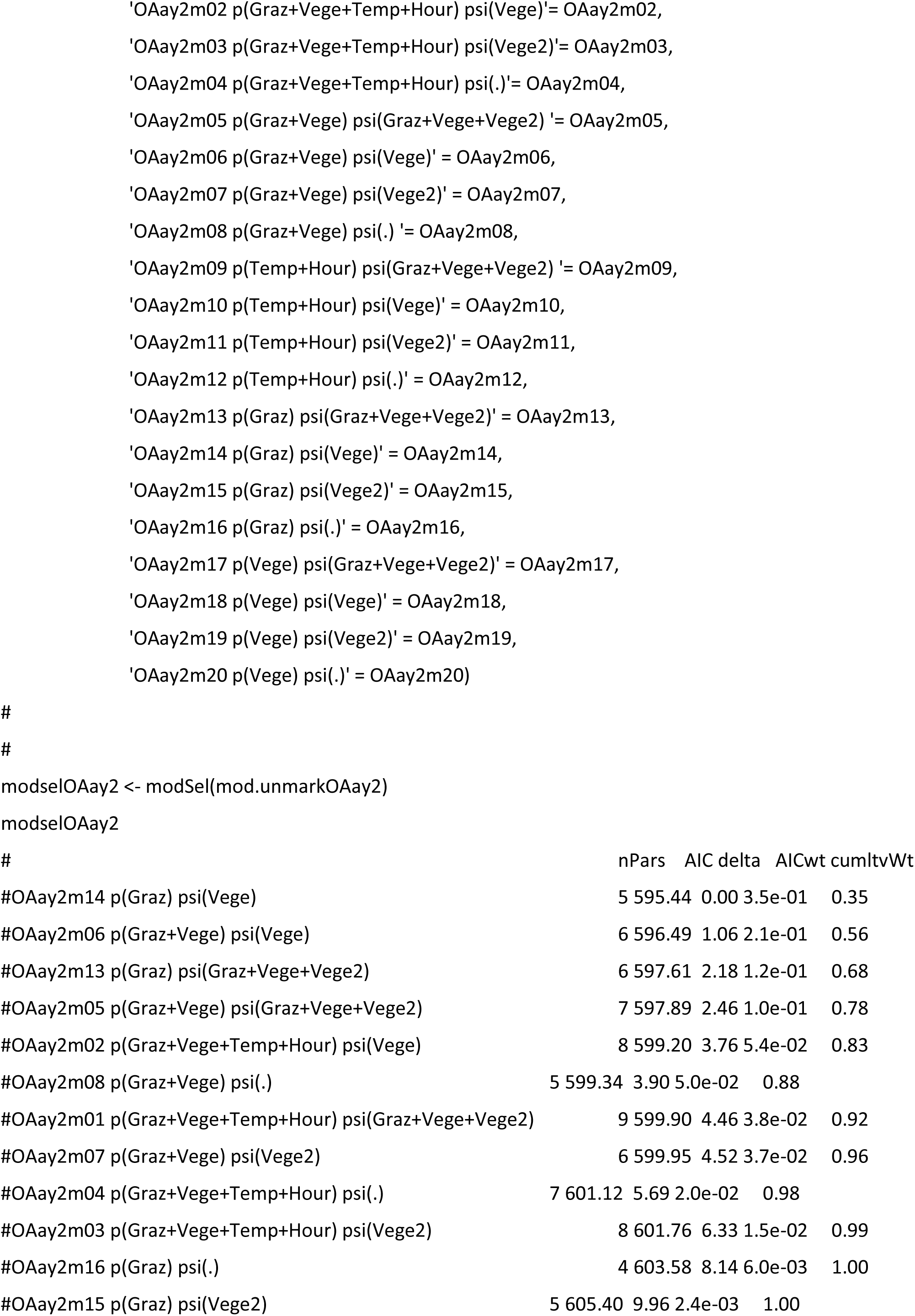
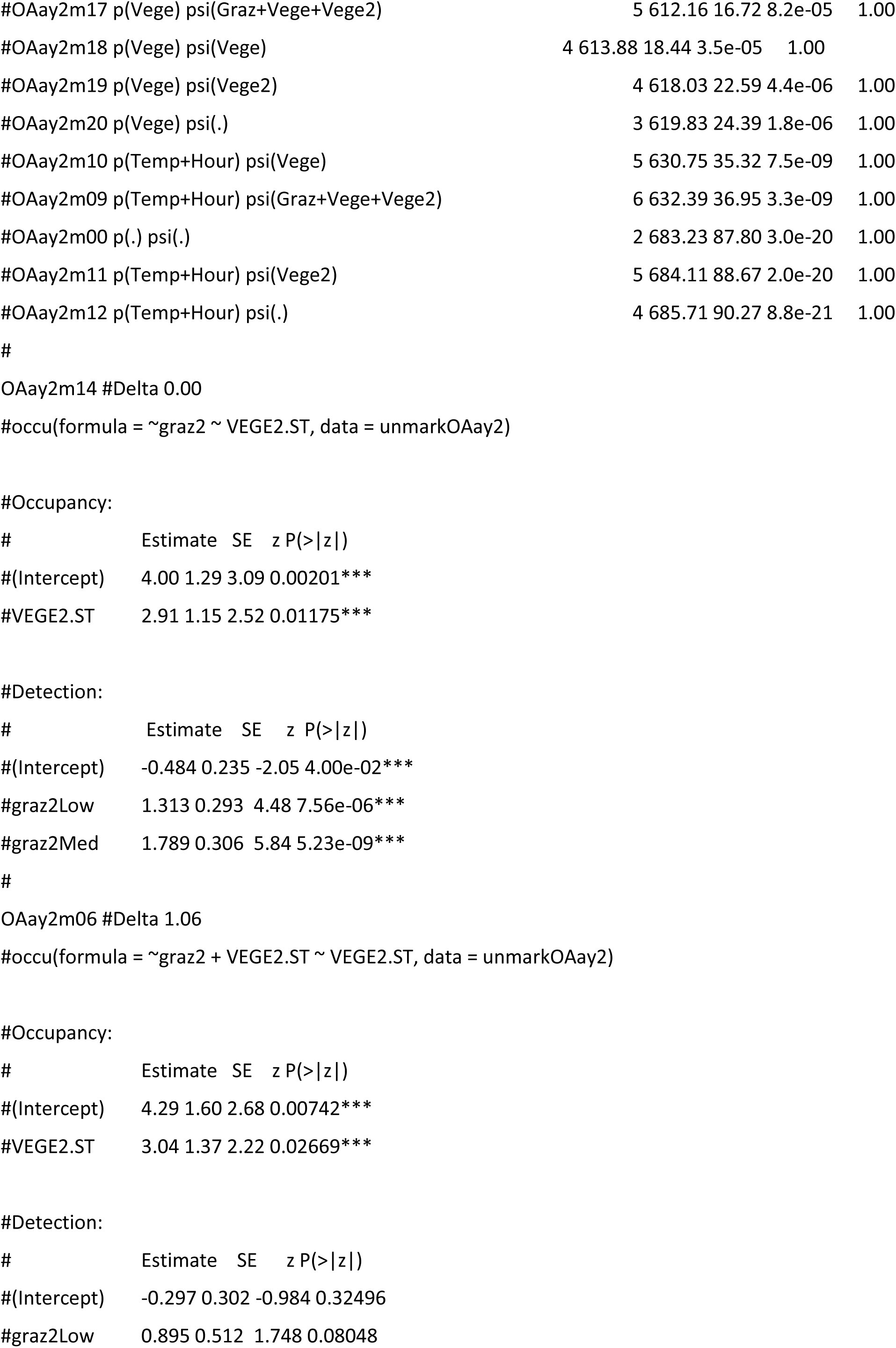
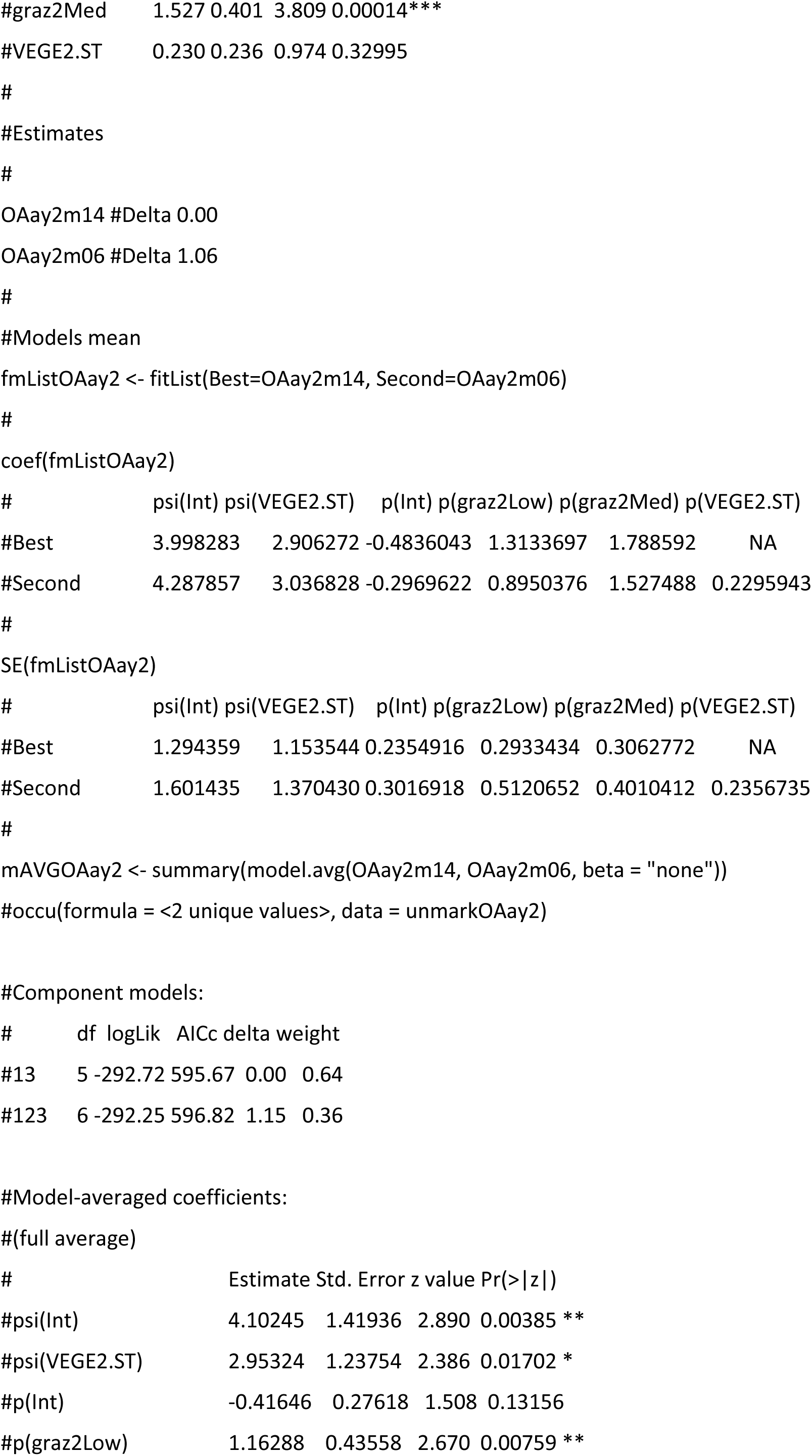
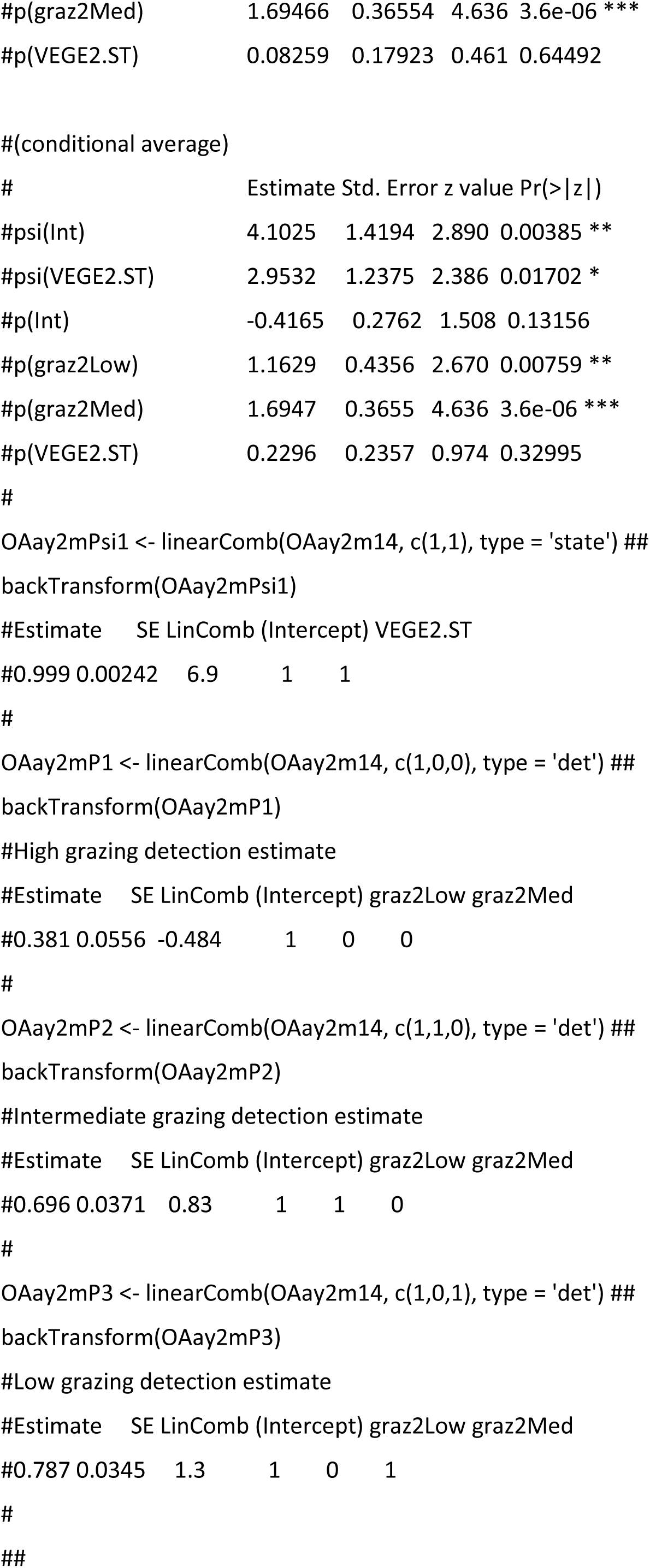
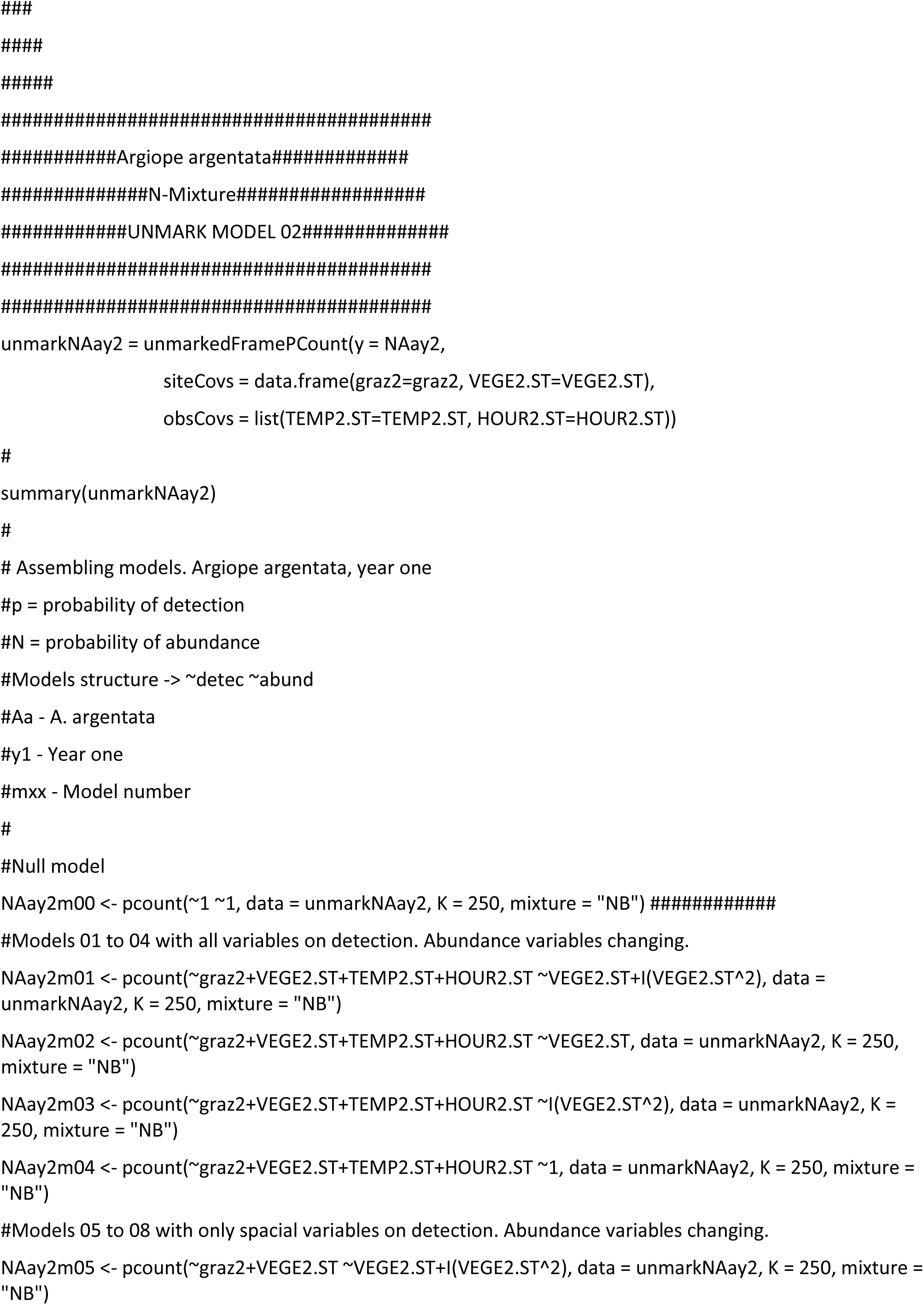
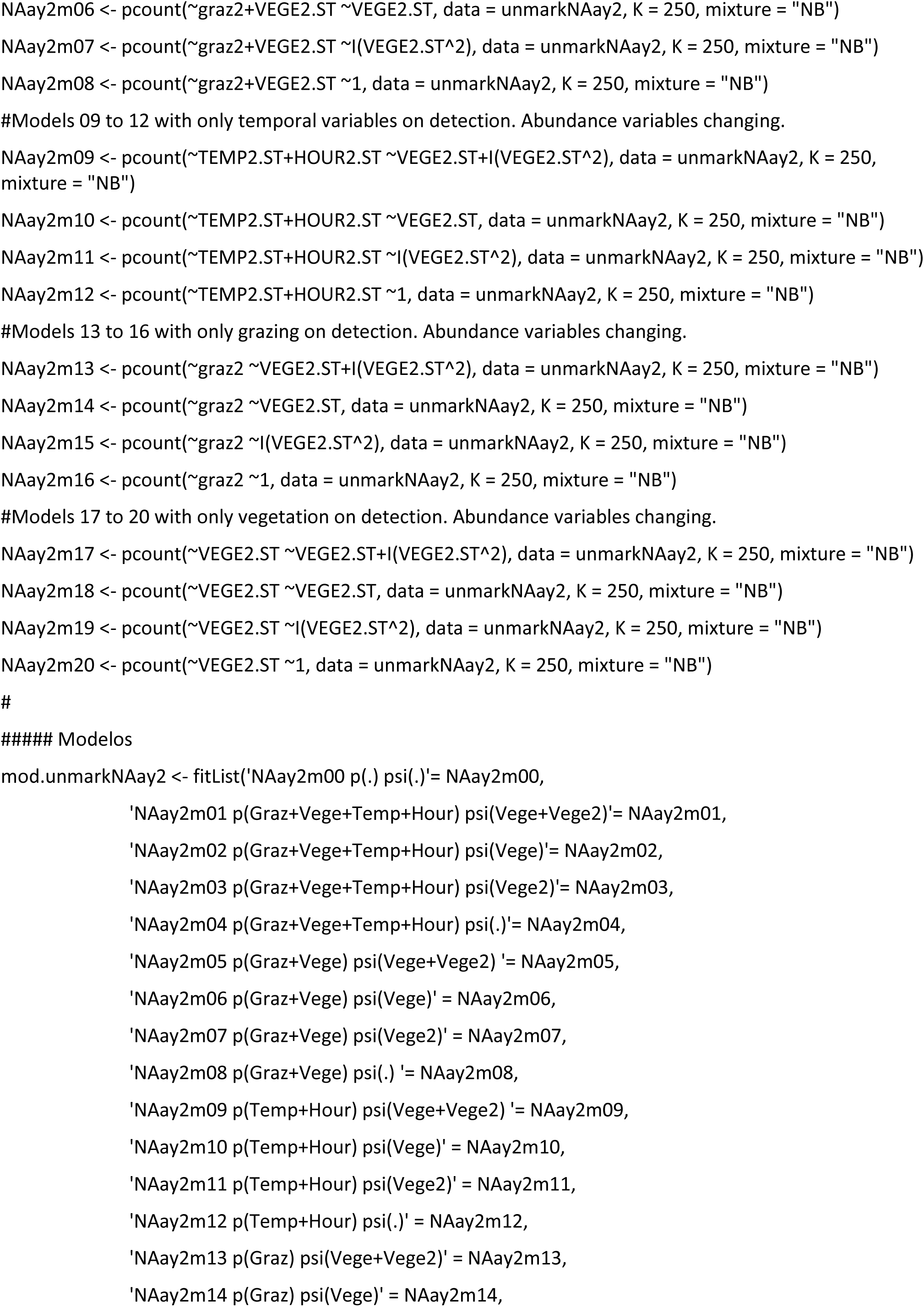
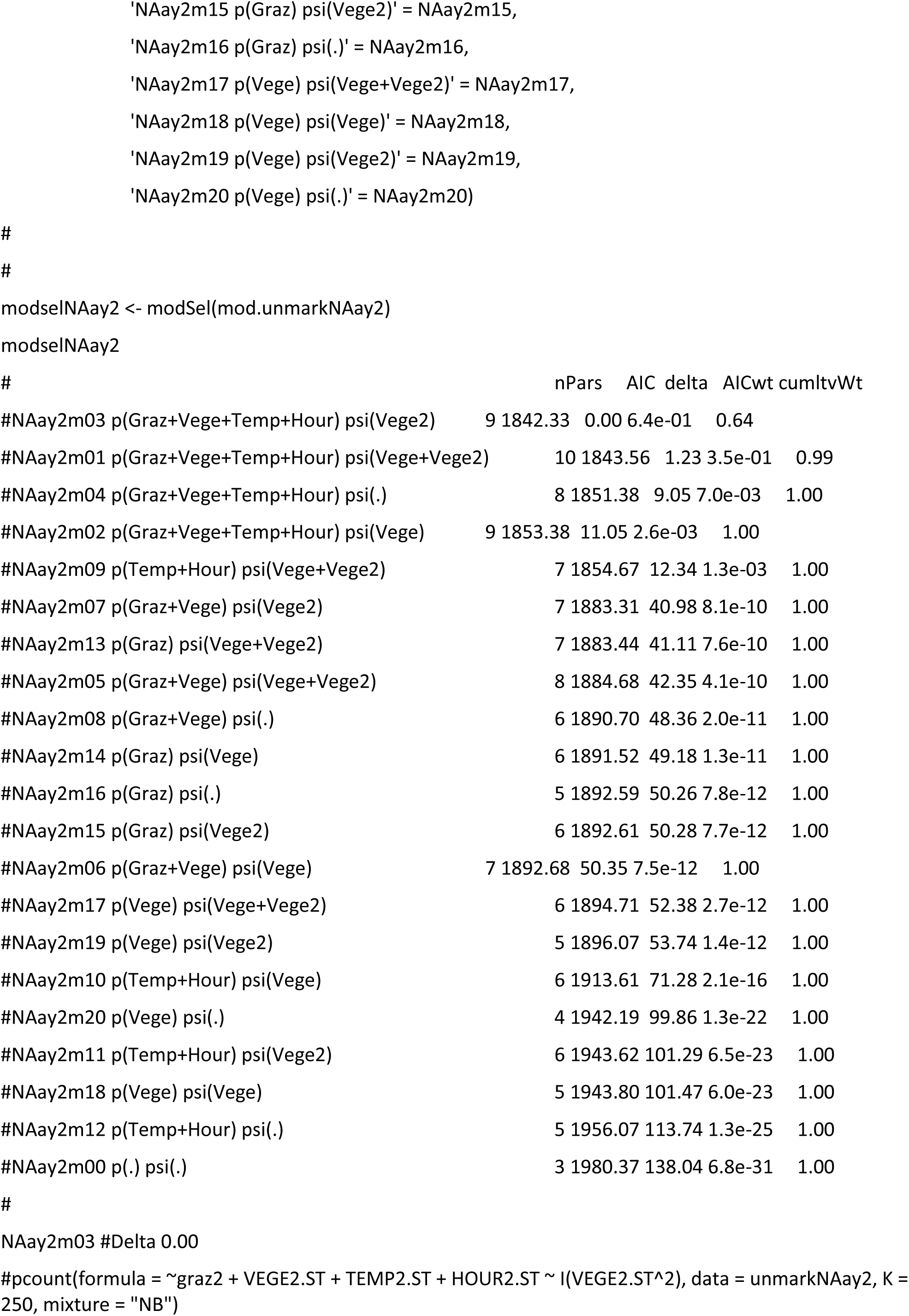
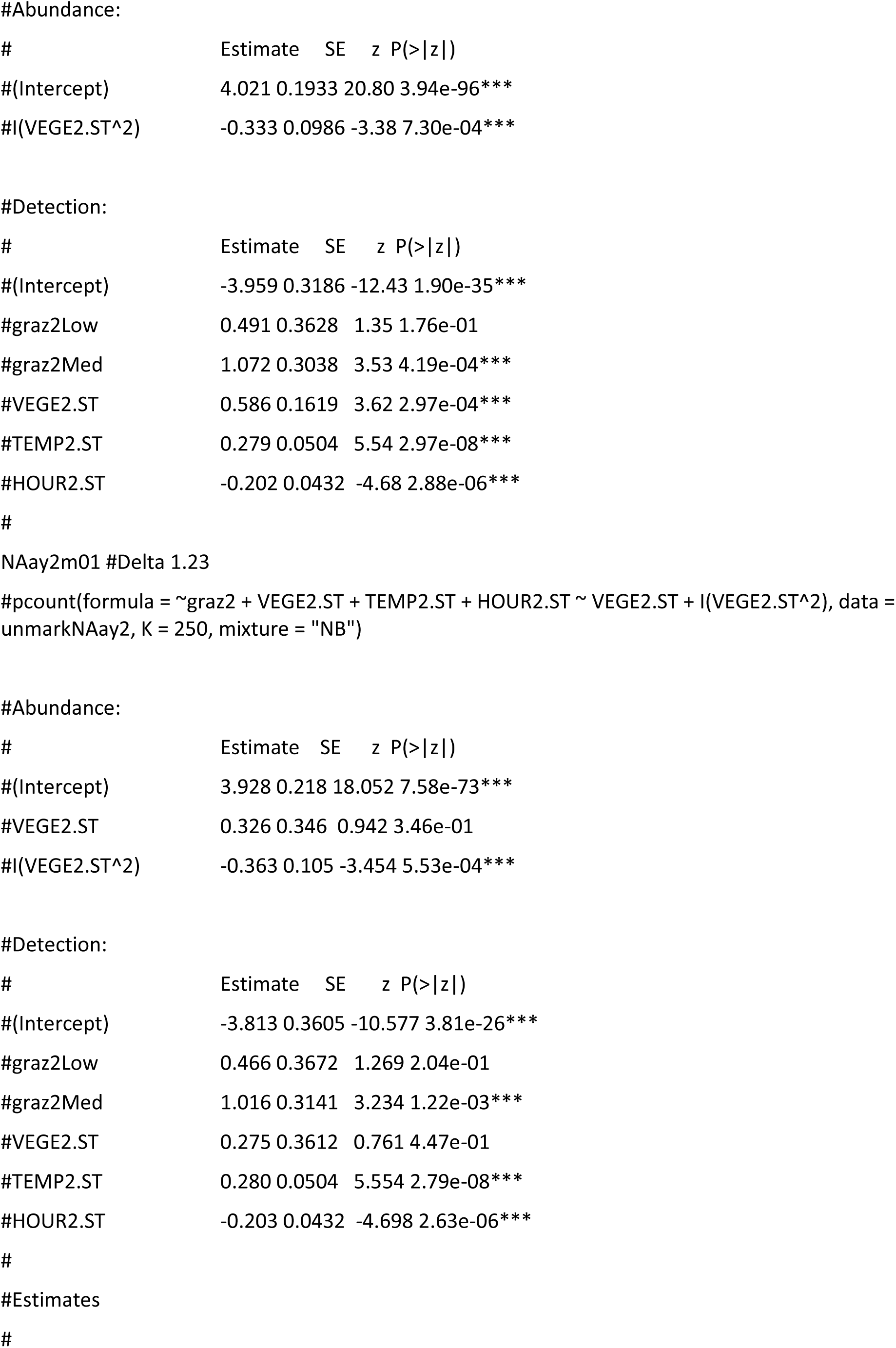
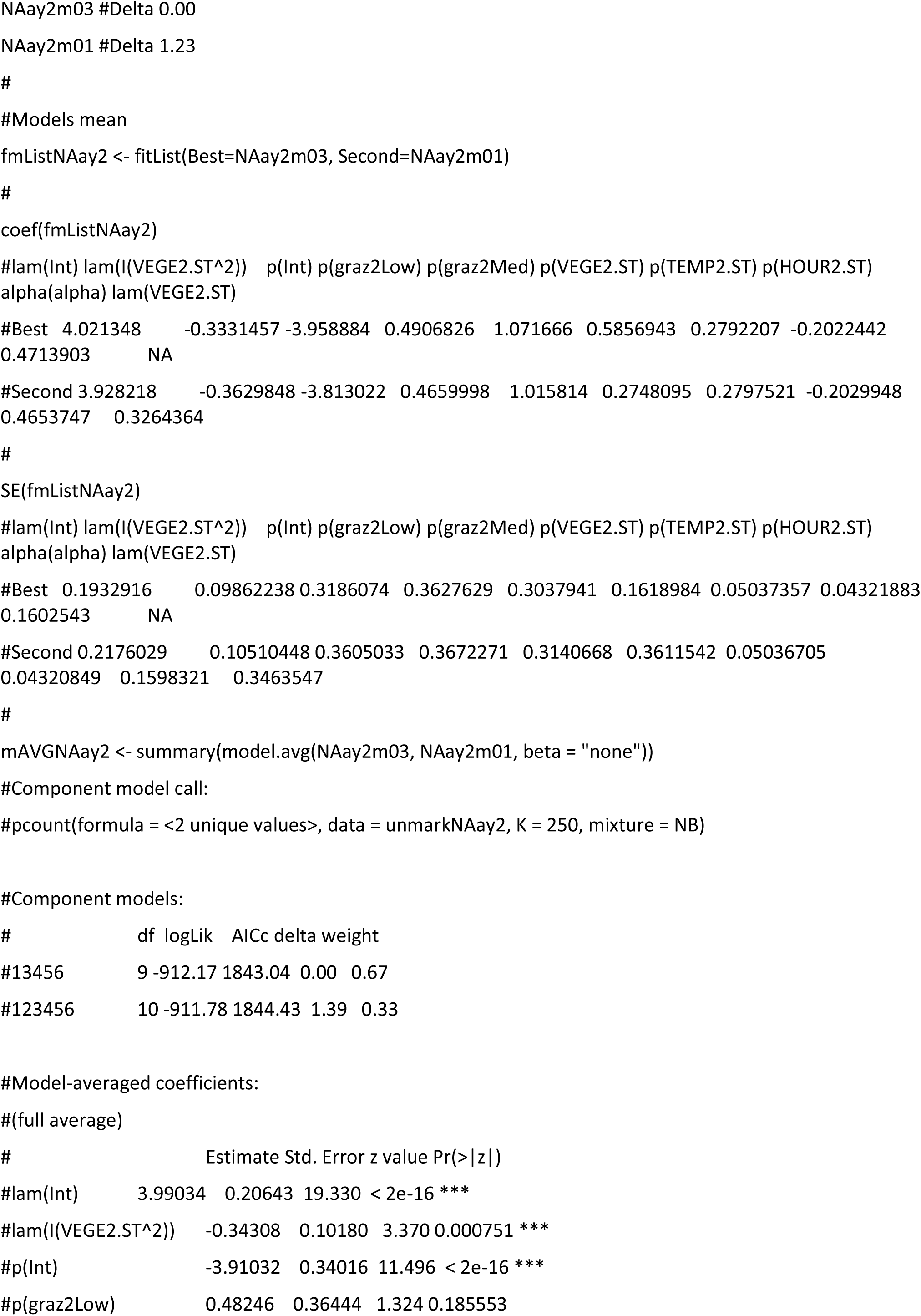
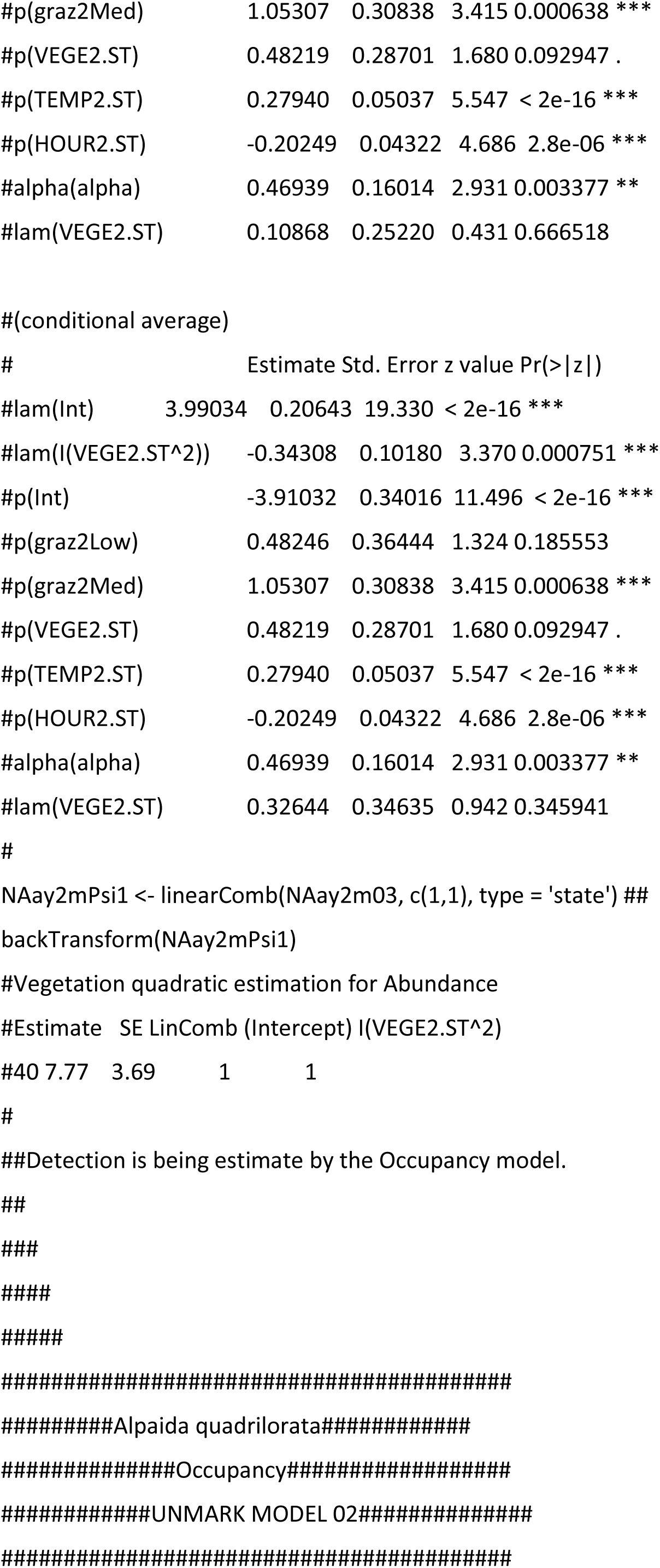
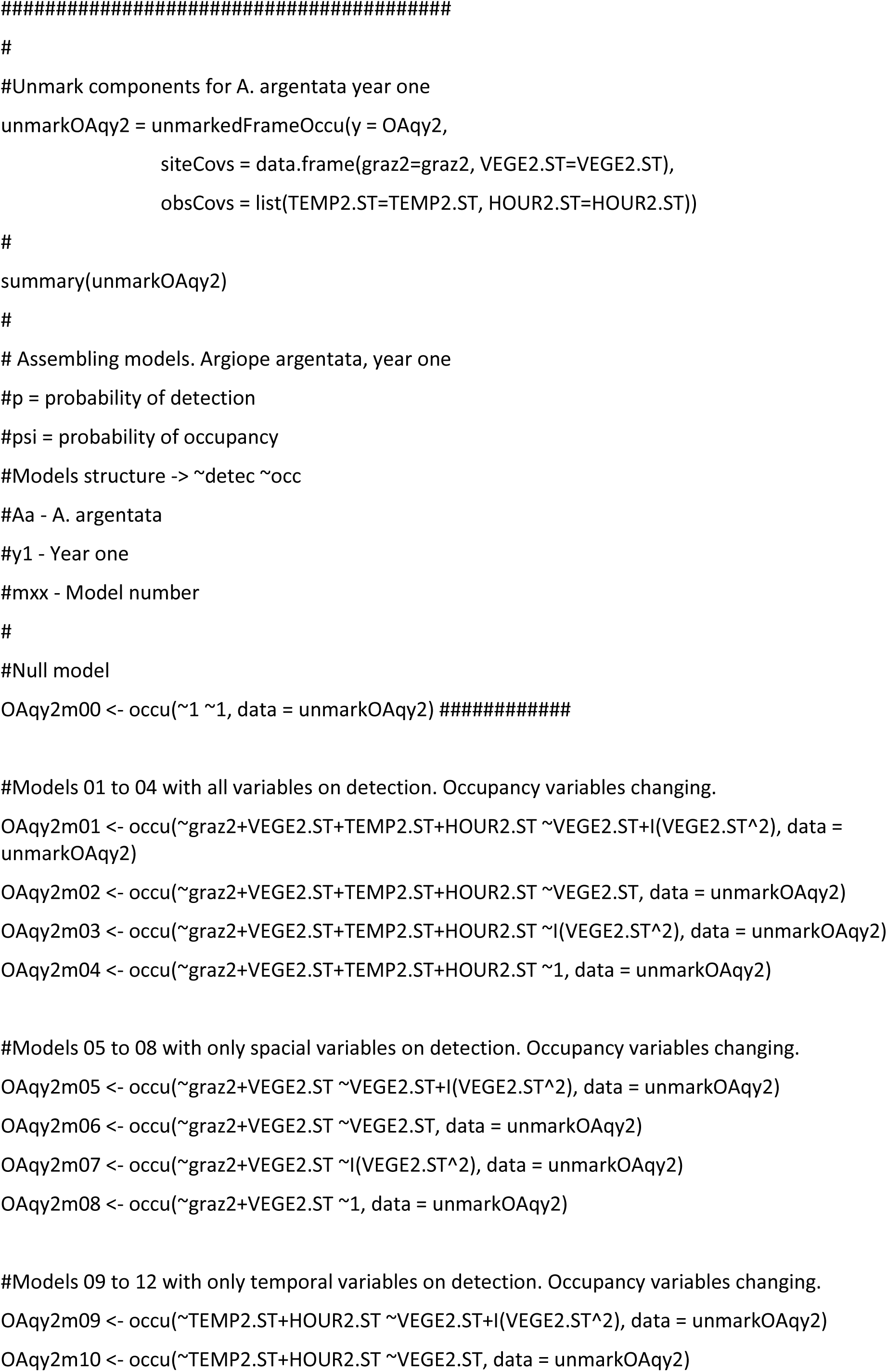
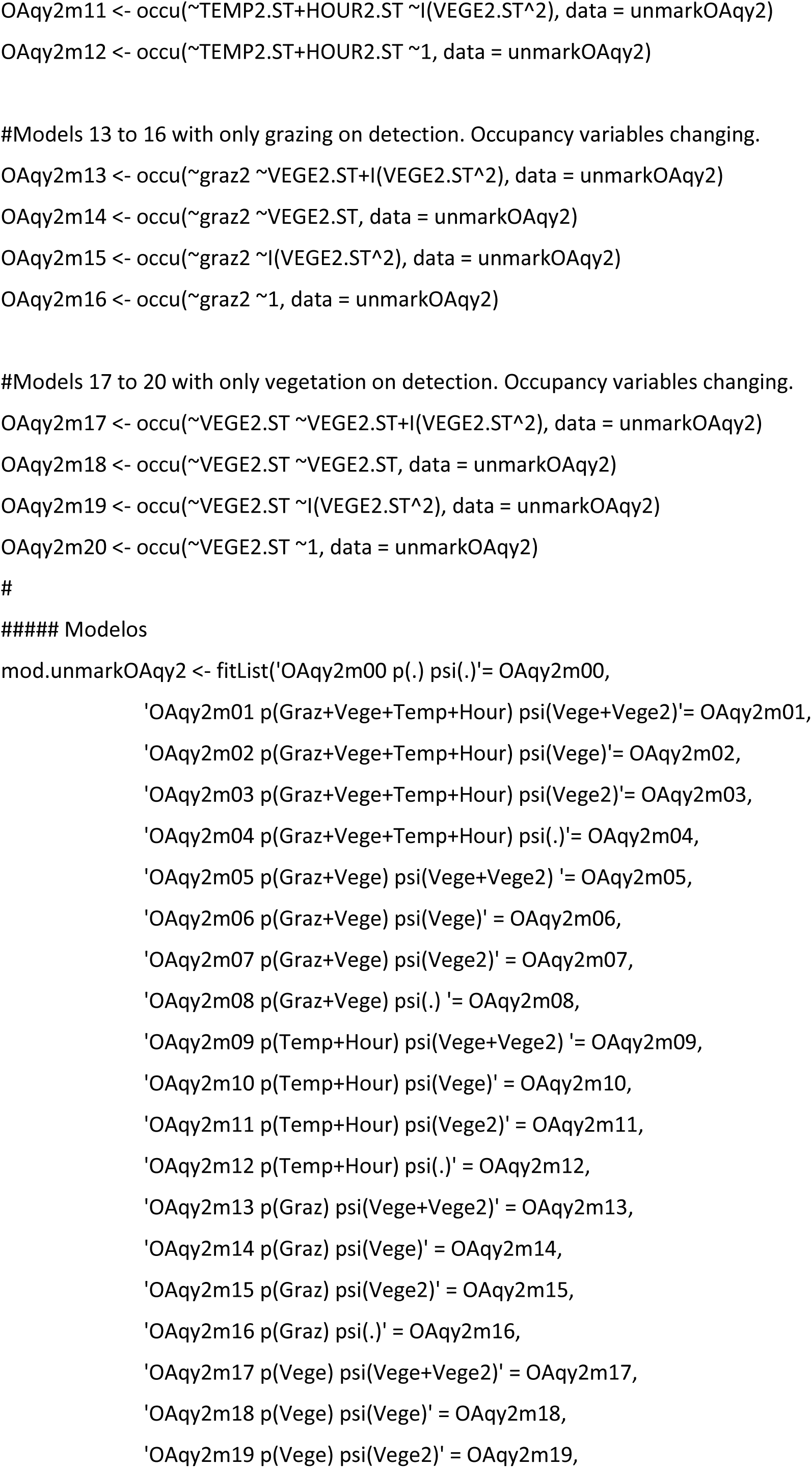
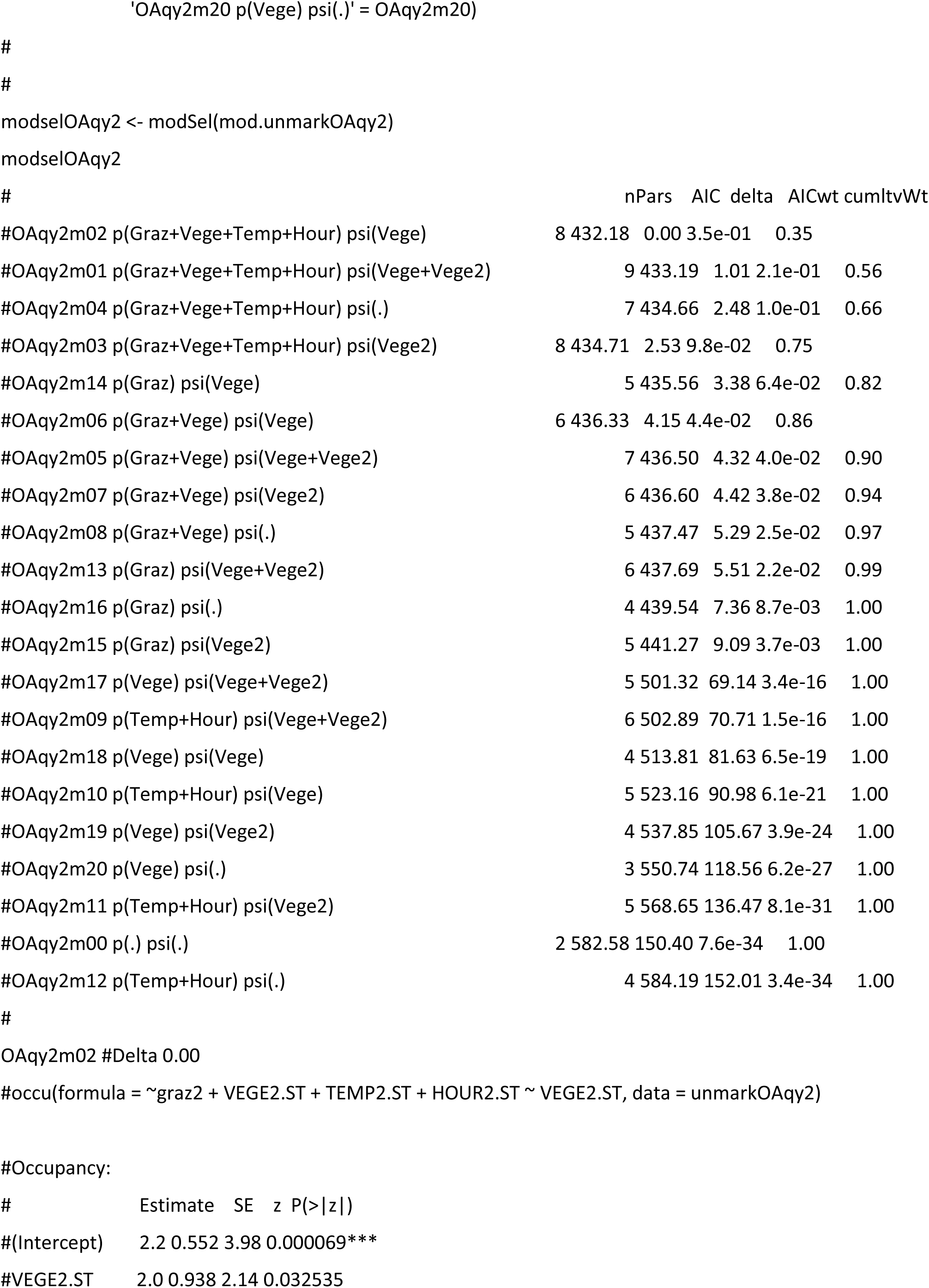
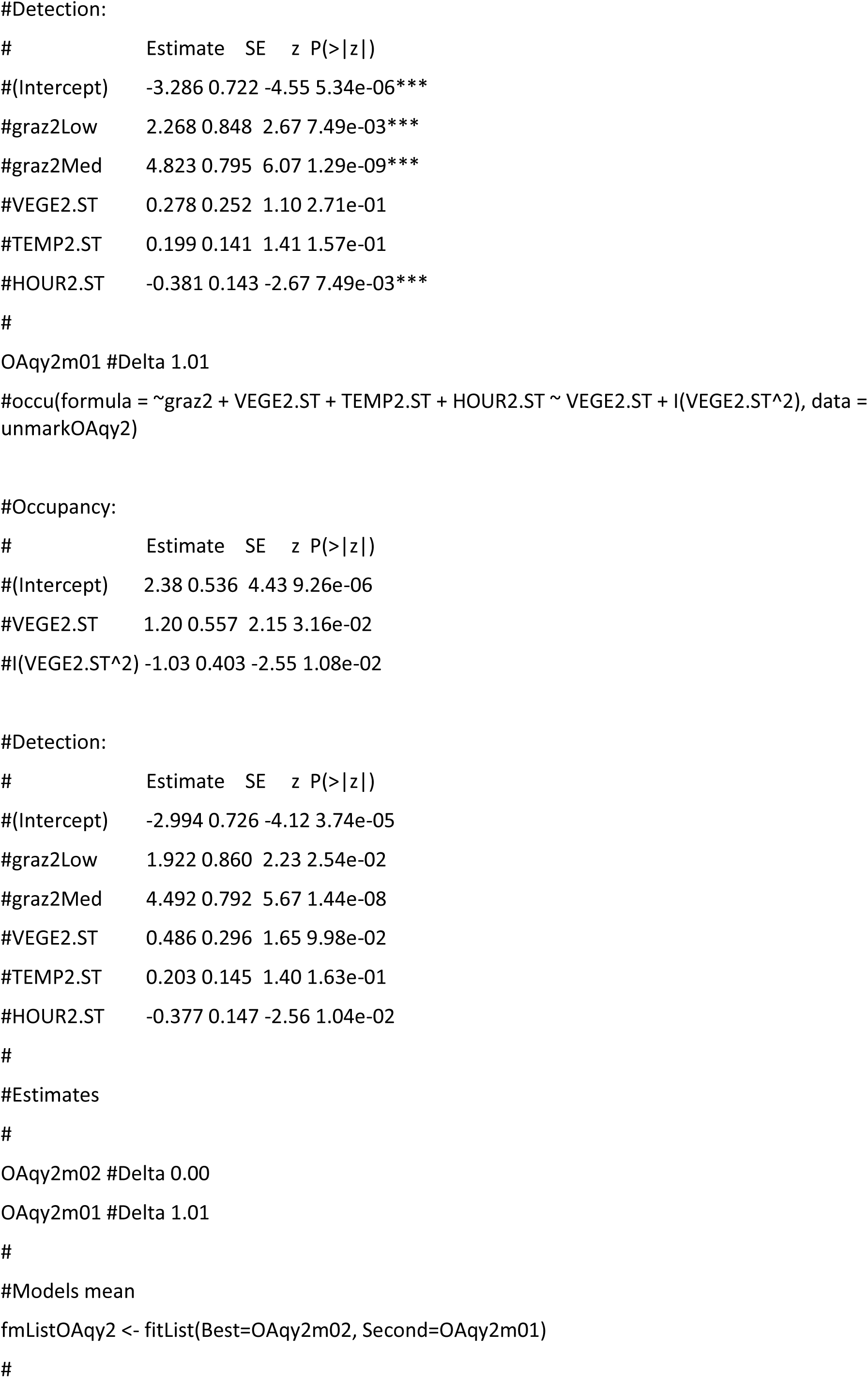
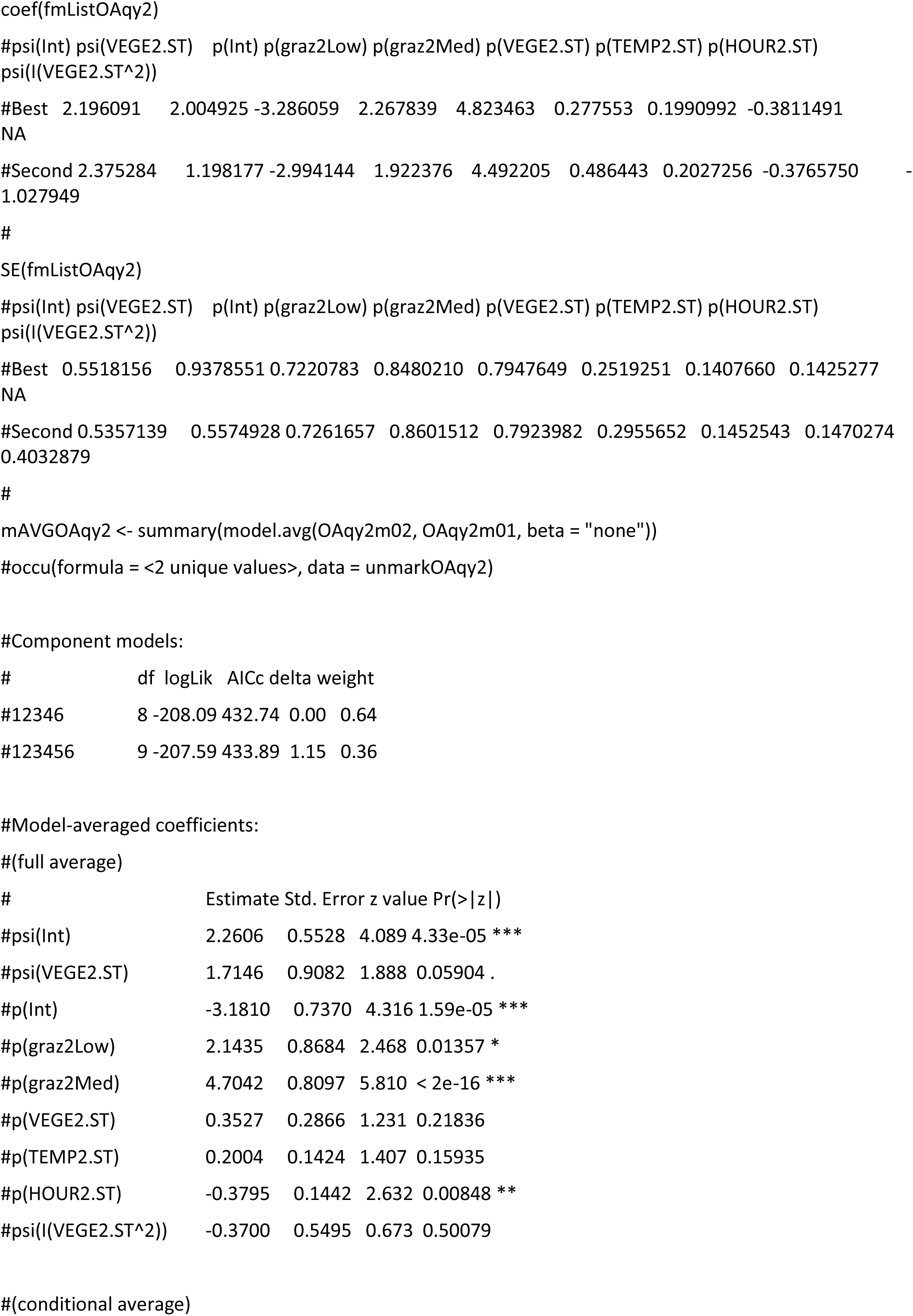
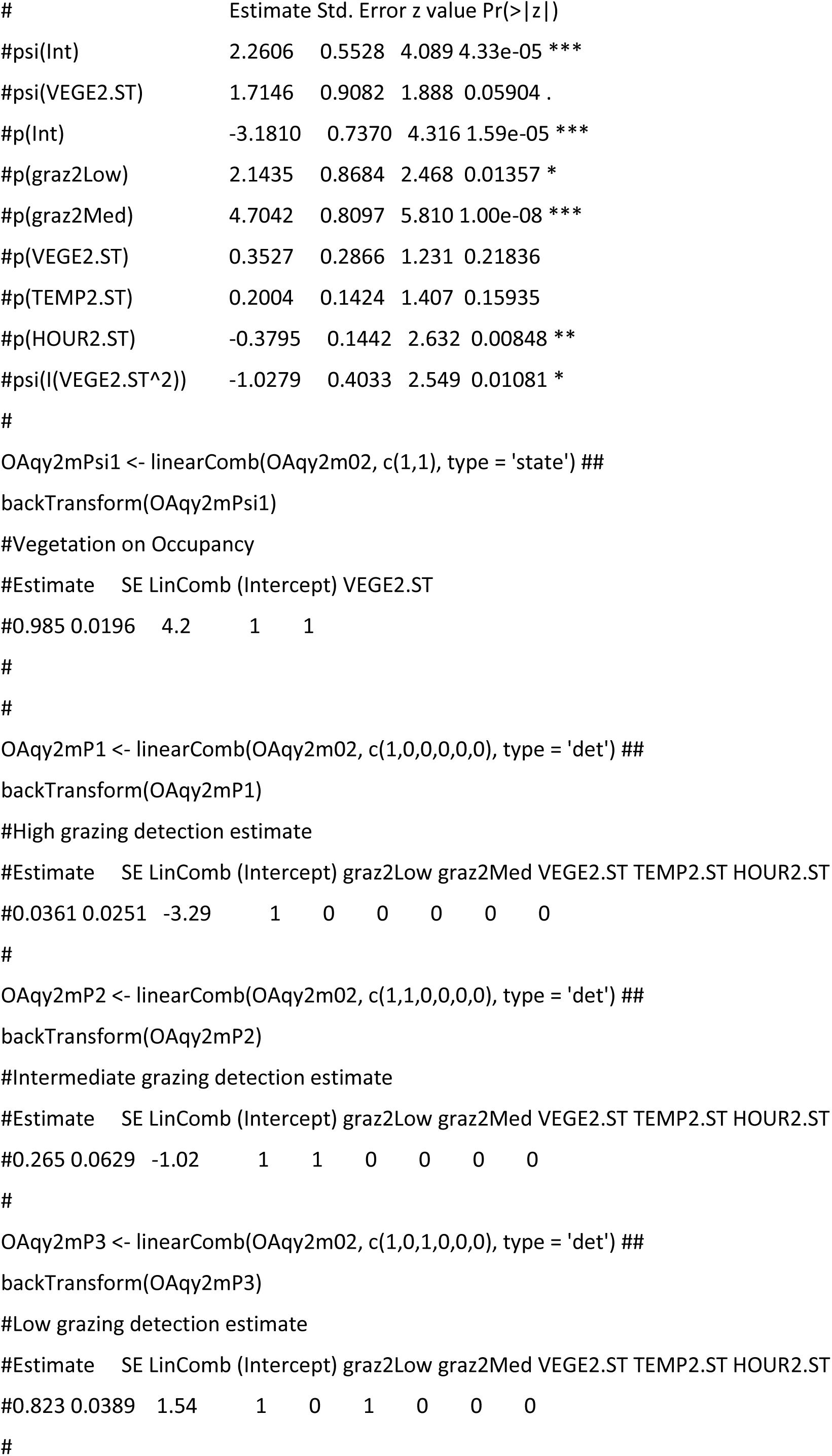
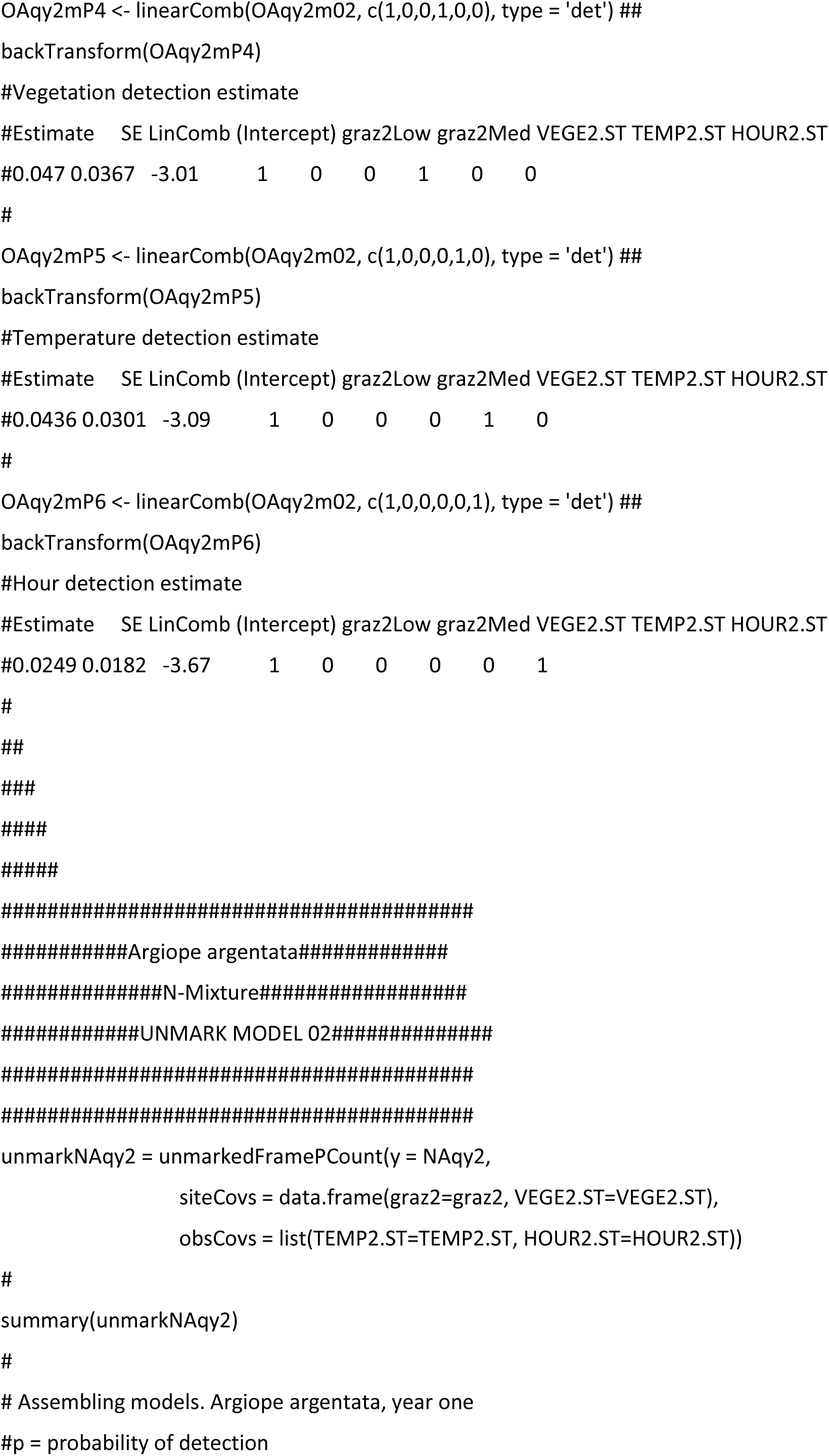
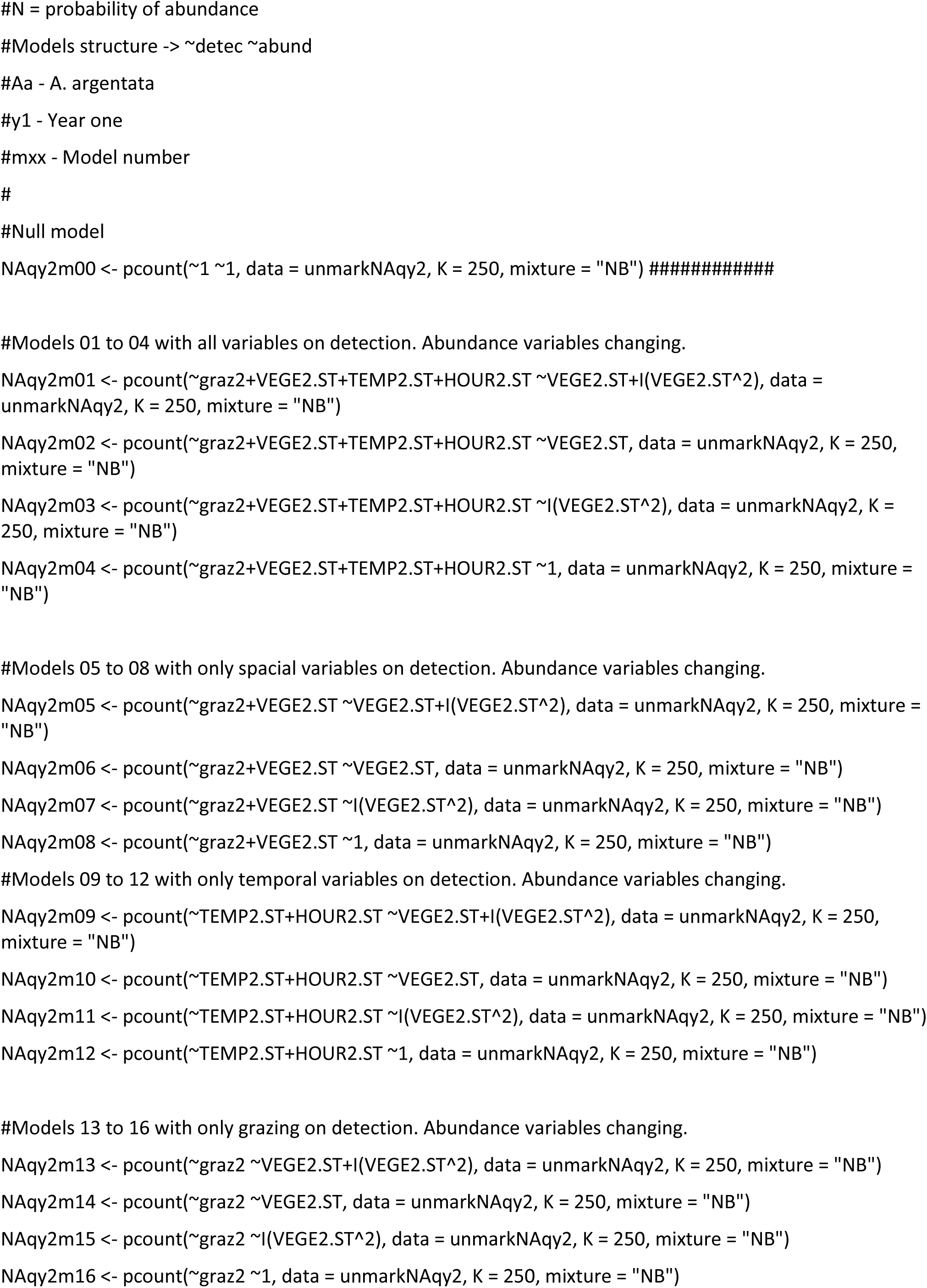
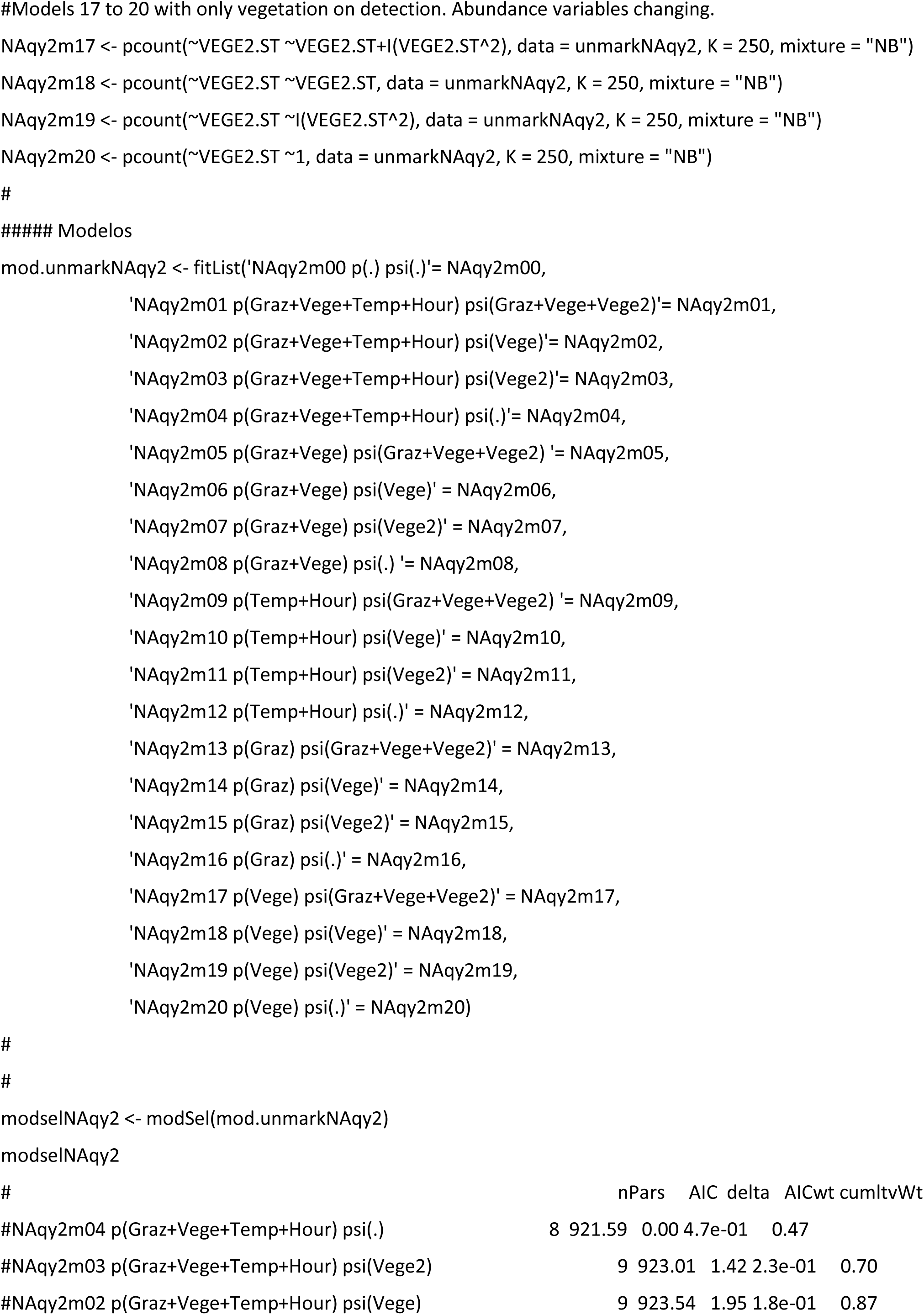
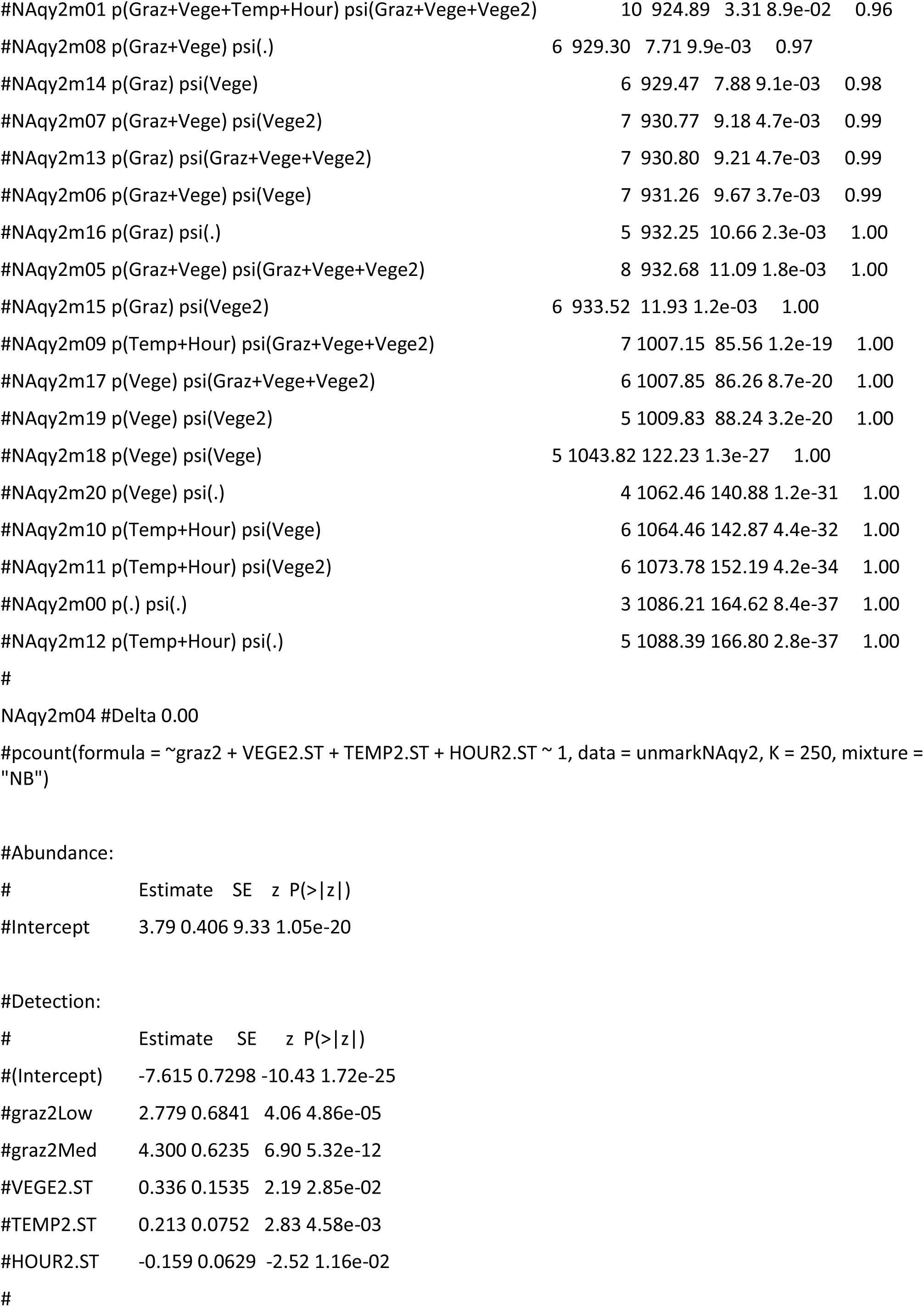
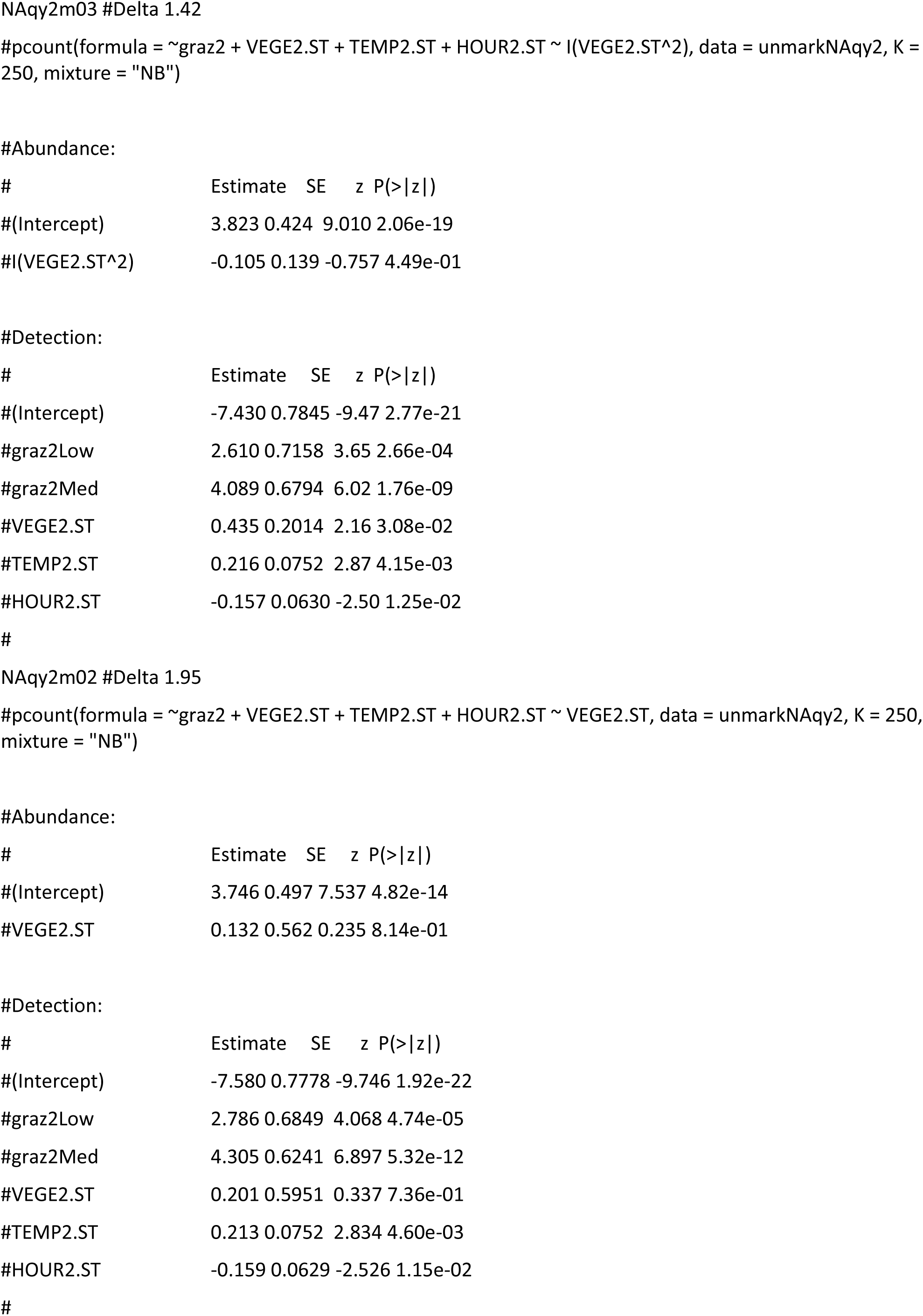
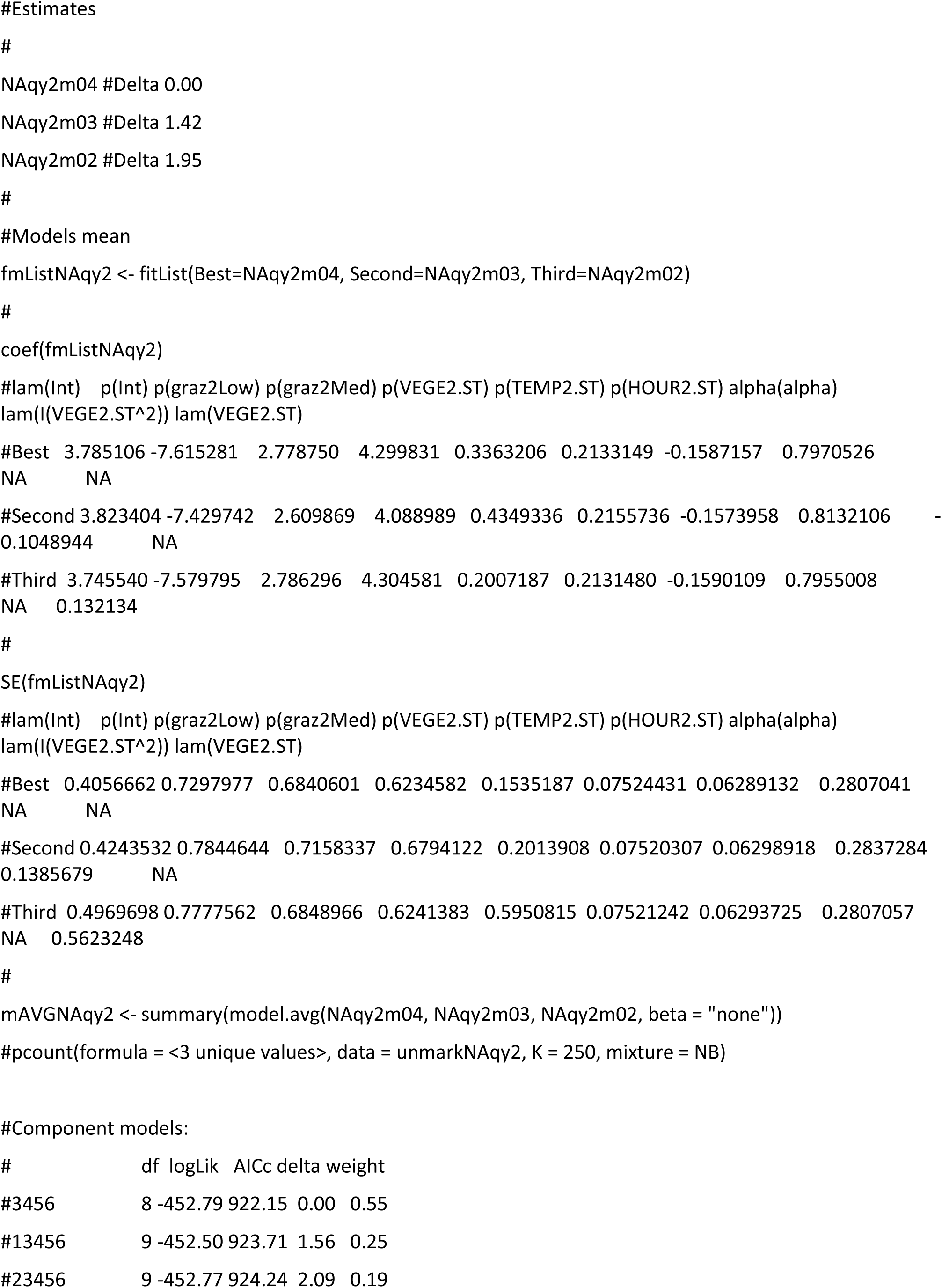
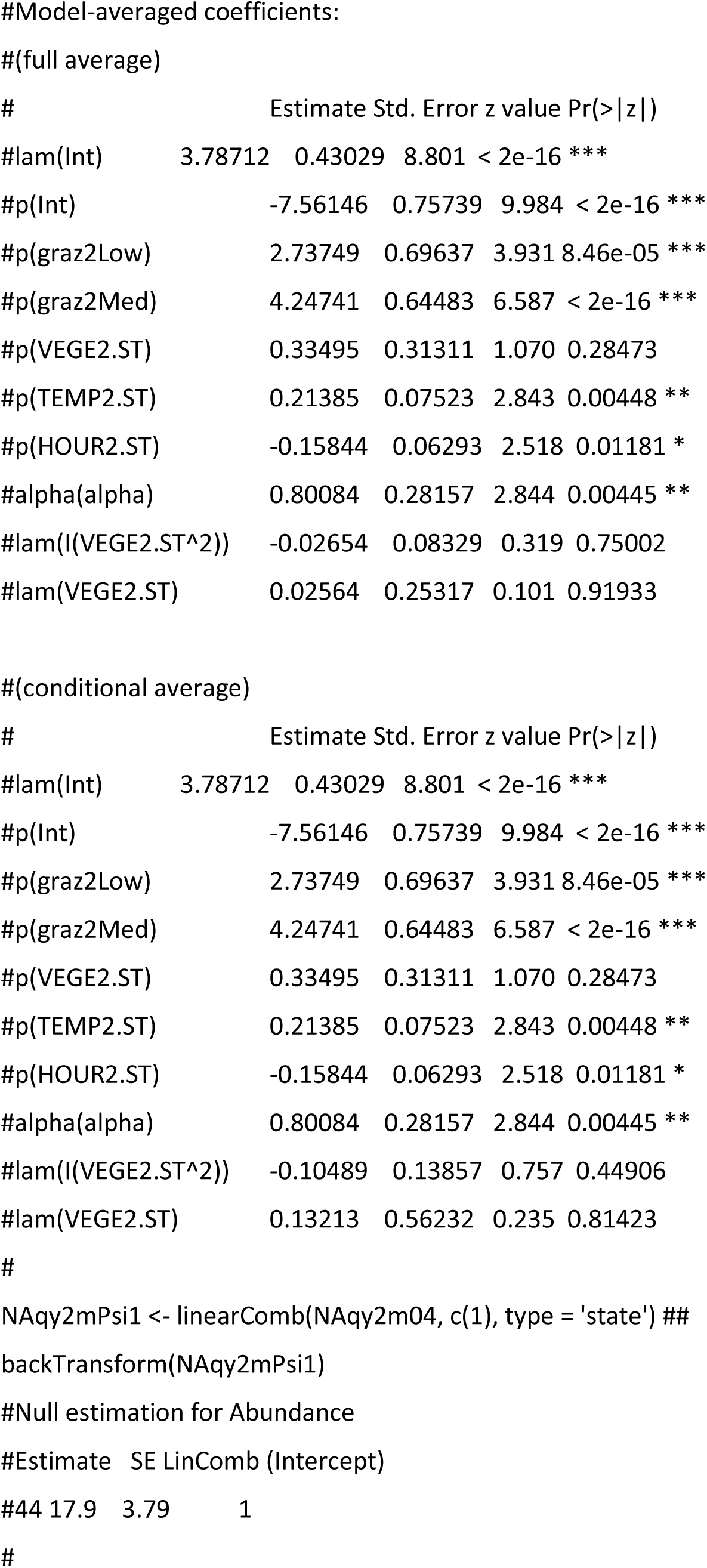
Orb-weaver spider species from the Araneidae family. Photos A and B represent *Argiope argentata* and photos C and D represent *Alpaida quadrilorata*.

